# Strain-release alkylation of Asp12 enables mutant selective GTP-state targeting of K-Ras(G12D)

**DOI:** 10.1101/2023.07.01.547359

**Authors:** Qinheng Zheng, Ziyang Zhang, Keelan G. Guiley, Kevan M. Shokat

## Abstract

K-Ras is the most commonly mutated oncogene in human cancer, yet direct small-molecule targeting of K-Ras mutants has been mostly unsuccessful until recently. The discovery of an allosteric pocket under Switch-II with covalent cysteine-crosslinking molecules has allowed for the development of targeted therapies that selectively engage the highly reactive acquired cysteine in the K-Ras(G12C) mutation without affecting the wild-type protein. Sotorasib and adagrasib, two advanced Switch-II Pocket inhibitors, have received FDA approval to treat K-Ras(G12C)-driven non-small cell lung cancer. However, the most frequent K-Ras mutation G12D particularly prevalent in pancreatic ductal adenocarcinoma has remained untargetable with covalent drugs due to the poor nucleophilicity of the somatic aspartate residue. Here we present a set of malolactone-based electrophiles which exploit ring strain to crosslink K-Ras(G12D) at the mutant aspartate to form stable covalent complexes. Structural insights from x-ray crystallography and exploitation of the stereoelectronic requirements for attack of the electrophile allowed development of a substituted malolactone which resisted attack by aqueous buffer but rapidly crosslink with the aspartate-12 of K-Ras in both GDP- and GTP-state. The signaling-competent GTP-state targeting allowed effective suppression of downstream signaling and proliferation of cancer cells harboring K-Ras(G12D) mutation, and tumor growth of cell line-derive xenograft in mice. Our results demonstrate the rational design of covalent inhibitors to target a non-catalytic carboxylic acid side chain in K-Ras(G12D) which has resisted traditional drug discovery efforts.

---

Oncogenic mutations of Ras are among the most common genetic alterations in human cancer, with an estimated disease burden of >3 million new patients per year worldwide^1^. Despite widespread appreciation of the importance of Ras in cancer, direct binding ligands which block downstream signaling were not reported until 2013^2^ due to the lack of obvious drug binding pockets in the protein. The only clinically approved K-Ras inhibitors are completely mutant specific as they rely on covalent recognition of the highly nucleophilic somatic cysteine residue of K-Ras(G12C). Recent pre-clinical reports^3,4^ of non-covalent K-Ras binding inhibitors have emerged which lack exclusive mutant specificity and exhibit varying degrees of biochemical preference for mutant K-Ras over the wild-type. The challenge in covalent targeting the most common single variant in K-Ras driven tumors (G12D), is that the carboxylic acid side chains of Asp and Glu are among the least nucleophilic heteroatom containing amino acid side chains. A further challenge for electrophiles capable of reacting with weak nucleophiles (Asp/Glu) is the need to withstand 55 M water and hypernucleophiles such as glutathione (GSH) in biological buffers. No suitable electrophiles for selectively targeting acidic amino acid side chains in aspartic or glutamic acids at physiological pH have been reported^5-8^ despite significant advances^9^ in targeting neutral or basic ones such as lysine^10-12^, tyrosine^13^, serine^14,15^, arginine^16^, methionine^17^, and histidine^18-20^. Here we report an electrophile which alkylates the mutant aspartate in K-Ras(G12D) in both GDP- and GTP-bound states, blocks effector interactions, and inhibits the growth of K-Ras(G12D)-driven cell lines.

## Results and Discussion

### Malolactones covalently modify Asp12 of K-Ras(G12D) mutant

To covalently target the mutant aspartate, we focused on electrophiles designed to exploit strain-release upon carboxylate attack of small 3- and 4-membered heterocyclic ring systems. Our^21^ and others^22^ exploration of electrophiles from the literature showed modest levels of K-Ras(G12D) crosslinking exhibited by 3-membered rings — *NH*-aziridine and epoxide electrophiles covalently modified K-Ras(G12D) by 40–60% over a course of 24 h (Supplementary Figure 1). A recent report^23^ demonstrated that structurally similar epoxides could label recombinant K-Ras(G12D) at extremely high concentration (1 mM) which precluded its use as a targeted covalent inhibitor in cells. Our further optimization proved unsuccessful due to the limited number of chemically stable modifications possible in 3-membered rings. A covalent K-Ras(G12D) inhibitor based on cyclophilin recruitment has been reported in a meeting abstract in the absence of structural information^8^.

The 4-membered ring electrophile, β-lactone, found in both natural products and synthetic drugs^14,24,25^ was attractive because the parent β-propiolactone, a potent acylating and alkylating reagent^26-32^, modified K-Ras(G12D) at multiple sites including the target Asp12 residue (Supplementary Figure 2). However, the [4.2.0]-fused β-lactones that covalently acylated a sibling K-Ras(G12S) mutation^14^ were not reactive with G12D (Supplementary Figure 3a), requiring exploration of another means to activate the nucleophilic attack of the β-lactone by Asp12.

Ambident electrophile β-lactones react with carboxylates with a preference at the β-carbon via an alkylation pathway at the β-carbon^25,33^ (Cf. alcohols such as serine side chain prefer acylation via carbonyl attack). The alkylation trajectory in the [4.2.0]-fused β-lactones at the bridgehead carbon was not accessible to Codon12 side chains (Supplementary Figure 3b). We reasoned that switching the fused ring system to a simple carbonyl bridge would, 1) geometrically enable the β-lactone electrophile to be poised for S_N_2 attack from the P-Loop Asp12, 2) activate the alkylation pathway by carbonyl π*-orbital participation, and 3) potentially provide further hydrogen bonding activation by the neighboring Lys16 shown to be important in activating of K-Ras(G12C) acrylamide electrophiles^34^.

A series of recently reported high-affinity non-covalent small molecules^35,36^, including MRTX1133^3,37^, which binds to K-Ras(G12D) and WT K-Ras^38^ provided the necessary scaffold for presenting the electrophilic fragment toward the Asp12. We prepared compound **1** as a racemate by coupling (±)-β-carboxyl-β-propiolactone, also known as malolactone, to the bicyclic piperazine group of a Switch-II Pocket (S-IIP) ligand scaffold using the carbonyl bridge design (Figure 1a). Using whole protein mass spectrometry, we assessed the reactions between recombinant K-Ras proteins and 10 μM **1** at 23 ºC (Figure 1b,c). Compound **1** reacted rapidly with K-Ras(G12D)•GDP with a half-life of 99 s (95% CI: 83–118 s) and fully modified K-Ras(G12D)•GDP in less than 15 minutes (Figure 1d). By contrast, **1** only achieved 6.5 ± 0.3% modification of K-Ras(G13D)•GDP and no detectable modification of K-Ras(wildtype)•GDP after 1 h. Interestingly, Compound **1** showed markedly reduced reactivity with K-Ras(G12E)•GDP, a non-natural mutant that positions its carboxylate nucleophile (Glu) further from the backbone than that in G12D, and K-Ras(G12S)•GDP. Although less potent, compound **1** was also active against the GppNHp-bound K-Ras(G12D). This is distinct from S-IIP K-Ras(G12C) inhibitors, which exclusively recognize the GDP-bound state^2,34,39-42^, whereas GTP-state recognition has so far only been observed for compounds with a different binding mechanism^8,43^. A recently reported K-Ras(G12C) inhibitor dependent on cyclophilin recruitment is capable of GTP-state recognition and showed faster in-cell signaling inhibition kinetics^43^. The covalent modification of Asp12 by compound **1** stabilized both K-Ras(G12D)•GDP and K-Ras(G12D)•GppNHp toward thermal denaturation (ΔTm = +10.3 ºC and +2.5 ºC, respectively). The adduct between K-Ras(G12D) and **1** was stable over pH 4.5– 7.5, and did not degrade in the presence of 1 vol% of dithiothreitol or hydrazine at 23 ºC showing its intrinsic resistance to nucleophiles in the bulk solution (Supplementary Figure 4).

**Figure 1.**
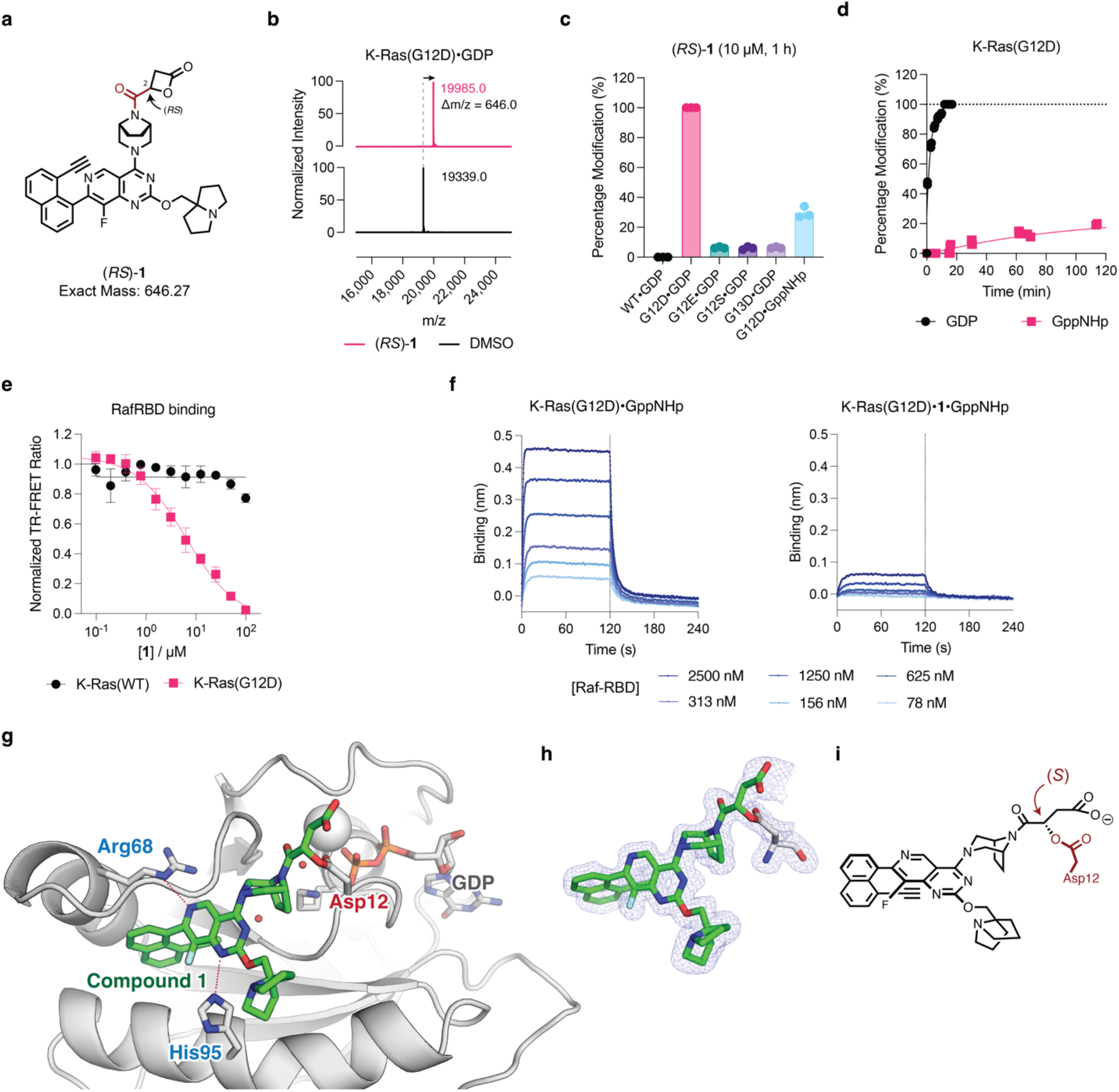
Malolactone (*RS*)-**1** is a selective, covalent inhibitor of K-Ras(G12D). **a**, Chemical structure of racemic (*RS*)-**1. b**, Deconvoluted protein mass spectra of Cyslight K-Ras(G12D)•GDP in the presence or absence of (*RS*)-**1**. Mass spectra are representative of two independent experiments. **c**, Compound (*RS*)-**1** selectively labeled K-Ras(G12D) not wildtype or other mutants. Data are presented as mean ± s.d. (*n* = 3). **d**, Kinetics of K-Ras(G12D) (200 nM) labeling with (*RS*)-**1** (10 μM) (*n* = 3, replicates are plotted as individual data points). **e**, Time-resolved fluorescence energy transfer dose-response of (*RS*)-**1** induced Ras-Raf•RBD binding disruption (*n* = 3). **f**, Biolayer interferometry dose-response of GST-Raf•RBD binding with K-Ras(G12D)•GppNHp without or with covalently labeled (*RS*)-**1. g**, Co-crystal structure of K-Ras(G12D)•GDP•**1. h**, 2*F*_o_ – *F*_c_ map for the covalently bound ligand **1** and Asp12 is depicted in blue mesh (1.0σ). **i**, Chemical structure of covalently bound ligand with the configuration of β-carbon assigned as *S*.

The ability of compound **1** to engage the GppNHp-bound state of K-Ras(G12D) prompted us to ask whether such a covalent modification could disrupt the interaction with effector proteins such as Raf. Using a time-resolved fluorescence energy transfer assay^44^, we found that compound **1** inhibited the interaction between Raf-RBD and K-Ras(G12D), but not wildtype K-Ras, in dose-dependent fashion (Figure 1e). We also measured the interaction between immobilized Raf-RBD with fully labeled K-Ras(G12D)•**1**•GppNHp using biolayer interferometry. Compared to unmodified K-Ras(G12D)•GppNHp, the K-Ras(G12D)•GppNHp•**1** complex showed substantially decreased binding to Raf-RBD (Figure 1f). To the best of our knowledge, this is the first example of a covalent S-IIP ligand directly impeding Ras-effector interaction in the GTP-state. Such a feature may be particularly advantageous for the G12D mutant, as its severely impaired GTP hydrolysis^45^ renders an abundant and persistent K-Ras population in the active GTP-state in the cell.

### X-ray crystal structure and kinetic analysis suggest an Asp12 O-alkylation mechanism

To better understand the covalent reaction in the S-IIP of KRAS (G12D) between **1** and Asp12, we solved a 1.7-Å crystal structure of the K-Ras(G12D)•GDP•**1** adduct (Figure 1g). We observed compound **1** in the S-IIP pocket adopting a conformation similar to that seen for K-Ras(G12C) inhibitors (Supplementary Figure 5), with clear electron density for the covalent ester bond between Asp12 and the compound as well as the free carboxyl group resulting from the ring opening. This high-resolution structure also allowed us to assign the stereochemistry of the adduct as *S* at the β-carbon (Figure 1h,i). Because we obtained the co-crystal using racemic (*RS*)-**1**, this *S* stereochemistry could in theory result from an S_N_2 attack on the β-carbon of *R* enantiomer of **1** or an attack on the carbonyl of the *S* enantiomer of **1** followed by an acyl transfer (Supplementary Figure 6a). To distinguish between these two possibilities, we prepared pure *R* and *S* enantiomers of a structural analog **2**. Monitoring the reaction rate with K-Ras(G12D)•GDP showed that the *R* enantiomer was significantly more reactive toward K-Ras(G12D)•GDP (Supplementary Figure 6b). Furthermore, we observed the same *R* >> *S* C_α_-enantioselectivity (i.e. C_2_ in malolactone system) in the context of K-Ras(G12D) labeling across a panel of strain-release electrophiles such as epoxides and aziridines which can only react via an S_N_2 mechanism (Supplementary Figure 1). Such enantioselectivity favoring a direct S_N_2 ring-opening mechanism aligns with literature observations with β-lactones^25,33^. The predominant S_N_2 preference of ring-opening reaction by Asp12 is distinct from the preference for attack by water, i.e. hydrolysis, which often takes place at the carbonyl in near-neutral buffers^46^.

### Mechanism- and structure-guided evolution of covalent K-Ras(G12D) active-state inhibitors

Despite being a potent covalent ligand of recombinant K-Ras(G12D), compound **1** was not stable in aqueous buffers at pH 7.4, precluding its use in cellular assays. We reasoned that its stability could be improved by attaching substituents at either or both sides of the α-carbon. Substitutions at the α-carbon will block the trajectory of incoming water nucleophiles from at least one side of the β-lactone ring (Figure 2a-lower path). The same modification of the electrophile was predicted to have little impact on the S_N_2 attack by Asp12 because of the pseudo-staggered conformation of the lowest unoccupied molecular orbital (LUMO) of the C–O bond and the α-carbon steric hindrance (Figure 2a-upper path).

**Figure 2.**
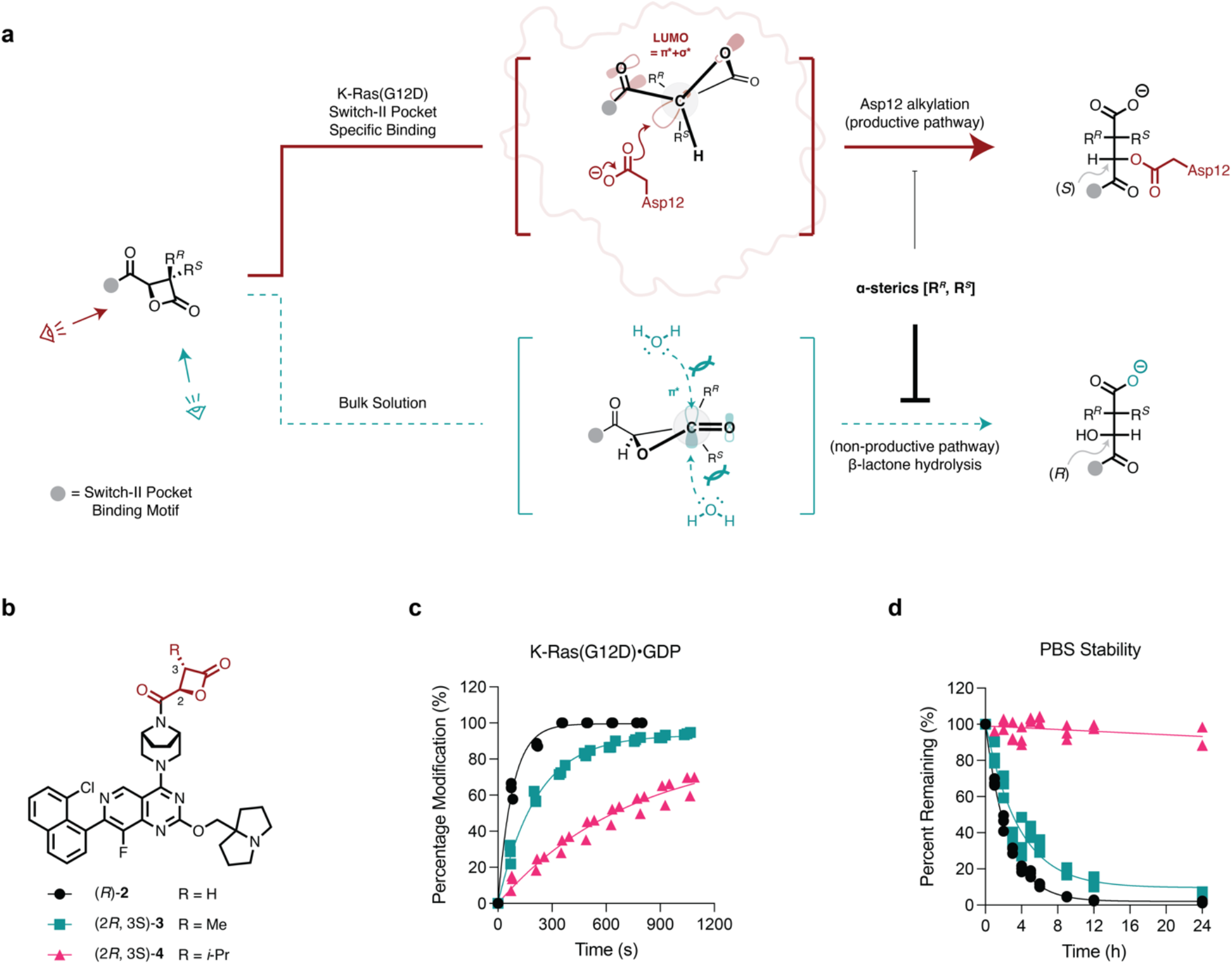
Sterically hindered β-lactones are stable and potent inhibitors of K-Ras(G12D). **a**, Mechanistic analysis of the Asp12-alkylating (red) and hydrolytic (teal) processes reveals the higher sensitivity of the latter to C3-substituents. **b**, Chemical structures of C3-substituted β-lactones (*R*)-**2**, (2*R*, 3*S*)-**3**, and (2*R*, 3*S*)-**4. c**, Labeling kinetics of 3-substituted β-lactones (*n* = 3, replicates are plotted as individual data points). **d**, Stability of 3-substituted β-lactones in phosphate-buffered saline (pH 7.4) by LC-MS (*n* = 3, replicates are plotted as individual data points).

To test this hypothesis, we prepared compounds **2**–**4** with increasing steric hinderance at the pro-*S* position of the α-carbon (Figure 2b). Compounds **2**–**4** maintained the ability to react with K-Ras(G12D), although the bulkier compounds exhibited decreased reaction rates (Figure 2c). Meanwhile, compound **4**, bearing an isopropyl group, resisted hydrolysis, with >90% of the material remaining intact after 24 h at 23 ºC and pH 7.4. By contrast, compounds **2** and **3** exhibited half-lives of 1.7 h and 2.5 h, respectively (Figure 2d).

With an improved isopropyl (*i*-Pr)-substituted malolactone electrophile, we further explored S-IIP ligands with higher-affinity, including the 8-ethynylnaphthyl ligand in Compound **1** [(2*R*,3*S*)-**5**] and MRTX1133 [(2*R*,3*S*)-**6**]^35-37^. The fast kinetics of GDP-state labeling persisted, Compound **5** and **6** labeled the GTP-state up to 200-fold faster than **4**, possibly due to the accessibility of the S-IIP of K-Ras(G12D)•GTP to the MRTX1133 scaffold. When tested at 10 μM, Compound **6** fully labeled 200 nM^47^ of K-Ras(G12D)•GppNHp within 5 min, which is, to our knowledge, the first S-IIP covalent molecule that has a preference for GTP-state of K-Ras (Figure 3b). Unlike MRTX1133 which prefers the GDP-state^48^, its (2*R*,3*S*)-3-isopropyl malolactone derivative Compound **6** preferred the GTP-state as assessed by intact protein mass spectrometry. The in-cell K-Ras(G12D) covalent labeling kinetics using a Western blot time course mirrored the recombinant protein results (Figure 3c), where Compound **6** labeled endogenous K-Ras(G12D) completely in homozygous *KRAS*^G12D/G12D^ cancer cell line SW1990 within 2 h as indicated by the gel mobility shift in the anti-Ras immunoblot, and concomitant reduction of the phospho-ERK levels. This is consistent with the higher biochemical potency we observed with K-Ras(G12D)•GppNHp, as cellular K-Ras(G12D) is enriched in the GTP-bound state as a result of its poor intrinsic and GAP mediated GTPase activity^45^.

**Figure 3.**
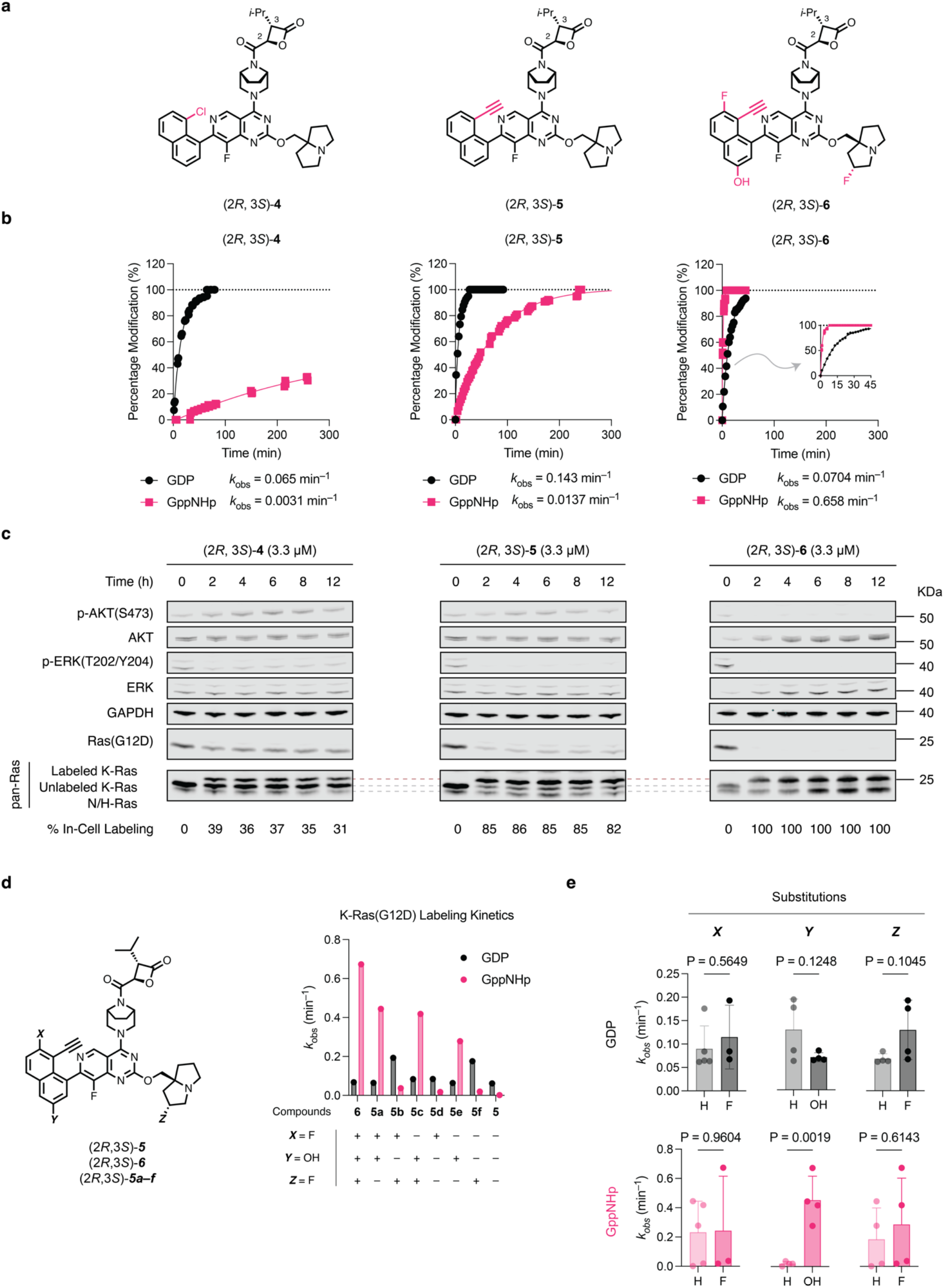
Rapid covalent modification of K-Ras(G12D)•GTP by β-lactone inhibitors is essential for in-cell target-engagement and oncogenic signaling suppression. **a**, Chemical structures of (2*R*, 3*S*)-**4**, (2*R*, 3*S*)-**5**, and (2*R*, 3*S*)-**6. b**, Recombinant K-Ras(G12D) labeling kinetics (*n* = 3, replicates are plotted as individual data points). **c**, Western blot time course of cellular K-Ras(G12D) covalent engagement and downstream signaling inhibition. **d**, Covalent K-Ras(G12D) labeling kinetics in both nucleotides by (2*R*, 3*S*)-**5**, and (2*R*, 3*S*)-**6**, and a compound analog series (2*R*, 3*S*)-**5a**–**5f. e**, Analysis of the determining substitution on covalent labeling kinetics by an unpaired t-test.

To understand the significantly improved GTP-state labeling kinetics from (2*R*,3*S*)-**5** to (2*R*,3*S*)-**6**, we synthesized six compounds [(2*R*,3*S*)-**5a** – (2*R*,3*S*)-**5f**, Figure 3D] exploiting their difference in the S-IIP binding core substitution pattern. Measuring the pseudo-first-order kinetic rates with 200 nM K-Ras(G12D) and 10 μM compounds, GTP-state labeling was more sensitive to these substitution changes with higher coefficient of variation (*CV*_GppNHp_ = 1.1, *CV*_GDP_ = 0.5). We grouped compounds with or without certain substitution, and analyzed the means of pseudo-first-order rates using student’s t-test (Figure 3E). While the two fluorine substitutions on the naphthyl ring or the bicyclic tertiary amine were dispensable, naphthyl 3-OH substitution was critical for fast GTP-state covalent labeling (*P* = 0.0019). Besides the gain of GTP-state labeling kinetics, the 3-hydroxy in the naphthyl ring significantly stabilized the covalent complex by 5–9 ºC (ΔΔTm, 3-hydroxy group versus 3-hydrogen, Supplementary Table 6). We reason that the hydrogen bonding (2.7 Å, as measured in PDB access code 7T47) between naphthyl 3-OH and Asp69 side chain is critical for S-IIP inhibitors to access the GTP-state and stabilize the complex. Similar interaction exists in previous GTP-state-selective cyclic peptide KD2 ^44^ and K-Ras(G12C) inhibitors ARS-853, ARS-107^34^, highlighting an underappreciated pharmacophore for active state targeting of K-Ras.

### α,α-Dimethyl malolactone (R)-7 selectively inhibited K-Ras(G12D) cancer cell lines

Despite efficient labeling of K-Ras(G12D) by Compounds **4**–**6** in cells, these compounds did not show significant mutant-selectivity in terms of cell growth inhibition in a survey of seven cancer cell lines including three *KRAS*^G12D^ mutant lines and four with either wildtype *KRAS* or non-G12D *KRAS* mutations (Supplementary Figure 7). We reasoned that Compounds **4**–**6** may possess cellular targets that are critical for cell survival other than K-Ras(G12D). We ruled out inhibition of WT K-Ras based on the effect on A375 cells which possess a downstream BRAF (V600E) activating mutation which should bypass K-Ras(WT) inhibition. We next turned to the monosubstituted malolactone electrophile in **4**–**6** which resembles that found in the natural product belactosin C^49-51^, a potent covalent inhibitor of the eukaryotic proteasome (Supplementary Figure 8). We confirmed that compounds **4**–**6** inhibited proteasome in a cell-based fluorescent assay and induced accumulation of poly-ubiquitinated proteins in HEK293 cells (Supplementary Figure 8). Because proteasome inhibition leads to cell growth inhibition independent of K-Ras(G12D) mutation, we sought to eliminate this off-target activity.

Through comparison of the co-crystal structure of a belactosin C derivative bound to the yeast proteasome (PDB: 3TDD) and that of compound **1** bound to K-Ras(G12D)•GDP, we hypothesized that introduction of a second substitution at the pro-*R* position would block proteasome binding and further improve hydrolytic stability (Supplementary Figure 9). Based on this design approach we also reasoned that the larger pro-*S* isopropyl substitution introduced previously would not be required if both hydrogens were substituted and therefore designed a 3,3-*gem*-dimethyl substituted malolactone, (*R*)-**7**. As predicted, (*R*)-**7** with a double substituted electrophile showed excellent stability (*t*_1/2_ > 24 h) in the presence of reduced glutathione in PBS at 37 ºC for 24 h (Supplementary Figure 10). (*R*)-**7** did not inhibit proteasome or induce protein poly-ubiquitination in HEK293 cells (Supplementary Figure 8). We further show that (*R*)-**7** was highly selective for K-Ras(G12D) and did not induce more than 20% inhibition in a panel of 482 kinases commonly used in safety profiling (Table S10). Compound (*R*)-**7** showed comparable labeling kinetics to **5**; it completely labeled K-Ras(G12D)•GDP and K-Ras(G12D)•GppNHp within 30 min and 300 min, respectively (Figure 4b). Compound (*R*)-**7** showed high mutant selectivity, labeling only the GDP- and GppNHp-state of G12D, but not the GDP-state of WT, G12E, G12S, or G13D despite elongated incubation time (Figure 4d). Compound (*R*)-**7** also stabilized both GDP and GppNHp-bound K-Ras(G12D) by 18.6 and 11.7 ºC, respectively (Supplementary Figure 11).

**Figure 4.**
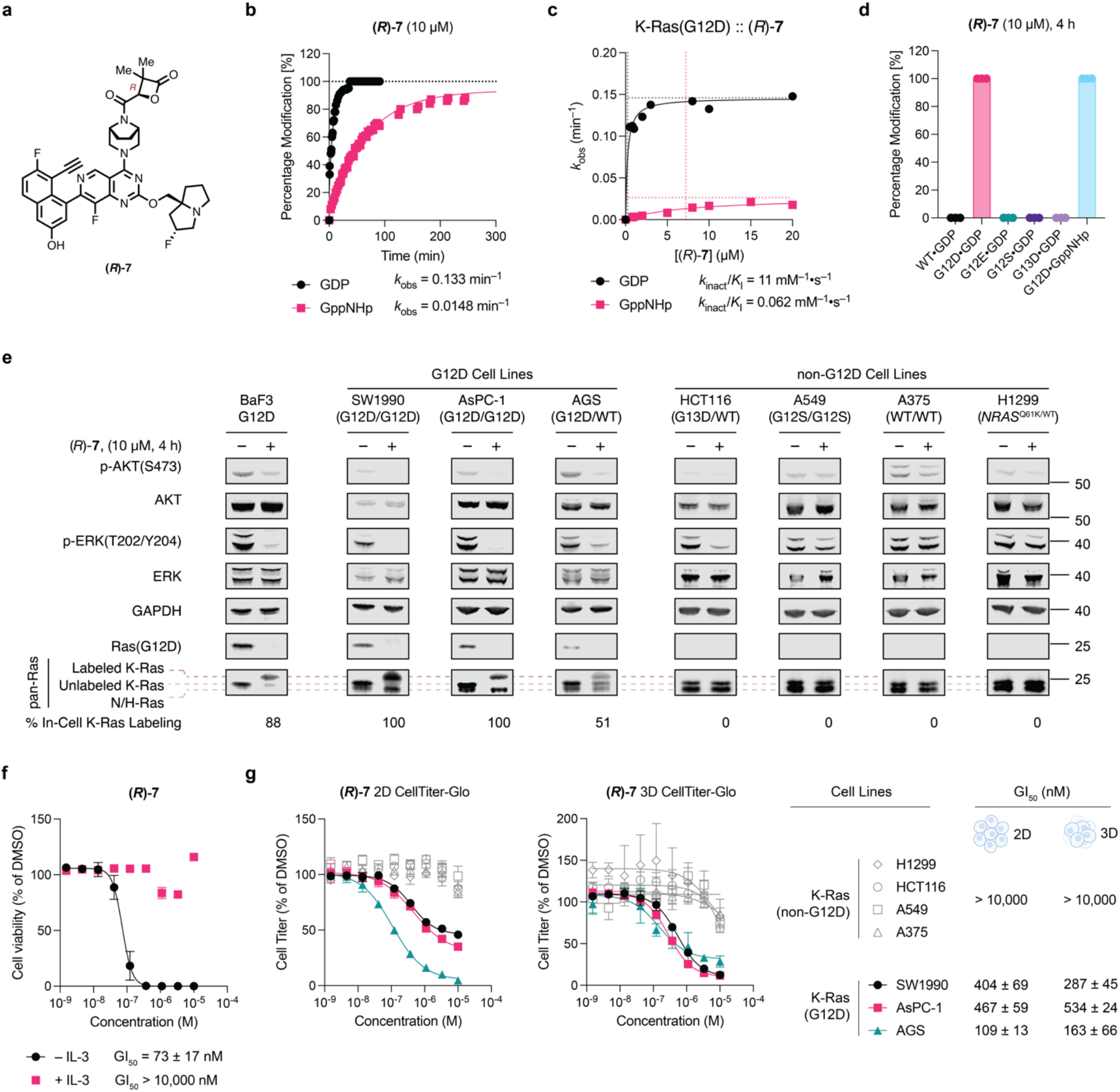
Covalent K-Ras(G12D) inhibitor (*R*)-**7** selectively inhibits cell growth in cancer cell lines harboring *KRAS*^G12D^ mutation and tumor growth in mice bearing SW1990 xenograft. **a**, Chemical structure of (*R*)-**7. b**, Pseudo-first-order K-Ras(G12D) labeling kinetics of (*R*)-**7**. Conditions: K-Ras(G12D) (200 nM), (*R*)-**7** (10 μM), room temperature. **c**, Second-order K-Ras(G12D) labeling kinetics of (*R*)-**7. d**, Covalent labeling selectivity against K-Ras wildtype and mutants (*n* = 3). **e**, Immunoblot of Ba/F3:K-Ras(G12D), SW1990 AsPC-1, AGS, HCT116, A549, A375, and H1299 cells treated with DMSO or 10 μM (*R*)-**7** for 4 h. **f**, Relative growth of Ba/F3:K-Ras(G12D) cells (with or without 10 ng/mL IL-3) after treatment with (*R*)-**7** for 72 h. Data are presented as mean ± s.d. (*n* = 3) and are representative of two independent experiments. **g**, Relative growth of cancer cell lines with (black, red, or teal) or without (grey) *KRAS*^G12D^ mutation after treatment with (*R*)-**7** for 72 h (2D) or 120 h (3D). Data are presented as mean ± s.d. (*n* = 3) and are representative of two independent experiments.

The mutant-selective, in-cell covalent labeling was assessed by immunoblots, where (*R*)-**7** labeled endogenous K-Ras(G12D) completely in homozygous *KRAS*^G12D/G12D^ cell lines SW1990 and AsPC-1, only half in heterozygous *KRAS*^G12D/WT^ AGS cell line, and none in non-G12D mutation cell lines (Figure 4e). Compound (*R*)-**7** inhibited growth of *KRAS*^*G12D*^-transformed Ba/F3 cells with an GI_50_ of 73 ± 17 nM. The same cells co-treated with interleukin-3 (IL-3, 10 ng/mL) lost sensitivity to (*R*)-**7** up to 10 μM suggesting the observed cell growth inhibition was due to K-RAS (G12D) inhibition. This compound further showed significantly biased inhibition profile toward K-Ras(G12D) mutation (SW1990, AsPC-1 and AGS) from non-G12D mutation cancer cell lines (H1299, HCT116, A549, A375) in 2D-adherent monolayer cultures. Compound (*R*)-**7** showed minimal toxicity to the latter cell lines at concentrations up to 10 μM. Similar cell growth inhibition potency as well as selectivity was observed in 3D-spheroid suspensions^39^.

Further, we tested three non-covalent analogs of our lead molecule (*R*)-**7**, including its hydrolyzed product and two non-lactone cyclic analogs, β-lactam and cyclobutanone. None of these molecules inhibited growth of K-Ras(G12D) cancer cell lines as potently as (*R*)-**7** (Supplementary Figure 12) highlighting the importance of covalency for mutant-selectivity and cellular potency. Together, our data demonstrate that strain release can be exploited to trap a common somatically mutated aspartate residue on the surface of the oncogene K-Ras(G12D). Despite the much weaker nucleophilicity of aspartate compared to cysteine, the second order rate constant (*k*_inact_/*K*_I_) for covalent bond formation of (*R*)-**7** with K-Ras(G12D)•GDP (11 mM^−1^ s^−1^) is comparable to those reported for approved K-Ras(G12C) inhibitors^34,40,41^ — 14 mM^−1^ s^−1^ and 35 mM^−1^ s^−1^, for sotorasib and adagrasib, respectively. The highly tunable malolactone ring system allowed selective detuning of water attack while preserving efficient attack by the weaker aspartate nucleophile. Compound (*R*)-**7** is a covalent ligand of K-Ras(G12D) with potent and mutant-selective cellular activity against K-Ras(G12D) cancers.

## Methods

### Recombinant protein expression and purification

K-Ras(wildtype), K-Ras(G12D), K-Ras(G12E), K-Ras(G13D), K-Ras (G12D) Cyslight DNA sequences encoding human K-Ras (wildtype, aa 1-169), human K-Ras (G12D, aa 1-169), human K-Ras (G12E, aa 1-169), human K-Ras (G13D, aa 1-169), and human K-Ras G12D Cyslight (G12D/C51S/C80L/C118S, aa 1-169) were codon optimized, synthesized by Twist Biosciences and cloned into pJExpress411 vector using the Gibson Assembly method^7^. The resulting construct contains N-terminal 6xHis tag and a TEV cleavage site (ENLYFQG). The proteins were expressed and purified following previously reported protocols.^2,8^ Briefly, chemically competent BL21(DE3) cells were transformed with the corresponding plasmid and grown on LB agar plates containing 50 μg/mL kanamycin. A single colony was used to inoculate a culture at 37 ºC, 220 rpm in terrific broth containing 50 μg/mL kanamycin. When the optical density reached 0.6, the culture temperature was reduced to 20 ºC, and protein expression was induced by the addition of IPTG to 1 mM. After 16 h at 20 ºC, the cells were pelleted by centrifugation (6,500 x g, 10 min) and lysed in lysis buffer [20 mM Tris 8.0, 500 mM NaCl, 5 mM imidazole] with a high-pressure homogenizer (Microfluidics, Westwood, MA). The lysate was clarified by high-speed centrifugation (19,000 x g, 15 min) and the supernatant was used in subsequent purification by immobilized metal affinity chromatography (IMAC). His-TEV tagged protein was captured with Co-TALON resin (Clonetech, Takara Bio USA, 2 mL slurry/liter culture) at 4 ºC for 1 h with constant end-to-end mixing. The loaded beads were then washed with lysis buffer (50 mL/liter culture) and the protein was eluted with elution buffer [20 mM Tris 8.0, 300 mM NaCl, 300 mM imidazole]. To this protein solution was added His-tagged TEV protease (0.05 mg TEV/mg Ras protein) and GDP (1 mg/mg Ras protein), and the mixture was dialyzed against TEV Cleavage Buffer [20 mM Tris 8.0, 300 mM NaCl, 1 mM EDTA, 1 mM DTT] at 4 ºC using a 10K MWCO dialysis cassette until LC-MS analysis showed full cleavage (typically 16-24 h). MgCl_2_ was added to a final concentration of 5 mM, and the mixture was incubated with 1 mL Ni-NTA (Qiagen) beads at 4 ºC for 1 h to remove TEV protease, any residual His-tagged proteins and peptides. The protein solution was diluted 1:10 v/v with 20 mM Tris 8.0 and further purified with anion exchange chromatography (HiTrapQ column, GE Healthcare Life Sciences) using a NaCl gradient of 50 mM to 500 mM in 20 mM Tris 8.0. Nucleotide loading was performed by mixing the ion exchange-purified protein with an excess of GDP (5 mg/liter culture) or GppNHp (5 mg/liter culture) and 5 mM EDTA at 23 ºC for 30 min. The reaction was stopped by the addition of MgCl_2_ to 10 mM. For GppNHp, an additional calf intestine phosphatase treatment was performed as follows to ensure high homogeneity of the loaded nucleotide. The protein buffer was exchanged into Phosphatase Buffer [32 mM Tris 8.0, 200 mM ammonium sulfate, 0.1 mM ZnCl_2_] with a HiTrap Desalting Column (GE Healthcare Life Sciences). To the buffer-exchanged protein solutions, GppNHp was added to 5 mg/mL, and Calf Intestine Phosphatase (NEB) was added to 10 U/mL. The reaction mixture was incubated on ice for 1 h, and MgCl_2_ was added to a final concentration of 20 mM. After nucleotide loading, the protein was concentrated using an 10K MWCO centrifugal concentrator (Amicon-15, Millipore) to 20 mg/mL and purified by size exclusion chromatography on a Superdex 75 10/300 GL column (GE Healthcare Life Sciences). Fractions containing pure Ras protein were pooled and concentrated to 20 mg/mL and stored at –78 ºC. In our hands, this protocol gives a typical yield of 5–15 mg/liter culture.

### Crystallization

K-Ras(G12D) Cyslight (G12D/C51S/C80L/C118S) bound by GDP purified by size exclusion chromatography was diluted to 20 μM in Reaction Buffer (20 mM HEPES 7.5, 150 mM NaCl, 1 mM MgCl_2_). Compound **1** was added as a 10 mM solution in DMSO to a final concentration of 50 μM. The mixture was allowed to stand at 23 ºC until LC-MS analysis of the reaction mixture showed full conversion to a single covalent adduct. The reaction mixture was concentrated using a 10K MWCO filter device and the adduct was purified by size exclusion chromatography (Superdex75, 20 mM HEPES 7.5, 150 mM NaCl, 1 mM MgCl_2_) and concentrated to 20 mg/mL. For crystallization, 0.1 μL of the protein was mixed with 0.1 μL well buffer containing 0.1 M MES 6.5, 25% PEG4K. Crystals were grown at 20 ºC in a 96-well plate using the hanging-drop vapor diffusion method. Maximal crystal growth was achieved after 7 days. The crystals were transferred to a cryoprotectant solution (0.1 M MES 6.5, 25% PEG4K, 25% Glycerol) and flash-frozen in liquid nitrogen.

### X-Ray Data Collection and Structure Determination

Dataset was collected at the Advanced Light Source beamline 8.2.2 with X-ray at a wavelength of 0.999907 Å. The dataset was indexed and integrated using iMosflm^52^, scaled with Scala^53^ and solved by molecular replacement using Phaser^54^ in CCP4 software suite^55^. The crystal structure of GDP-bound K-Ras(G12C)-MRTX849 adduct (PDB code: 6USZ) was used as the initial model. The structure was manually refined with Coot^56^ and PHENIX^57^. Data collection and refinement statistics are listed in Supplementary Table 1. In the Ramachandran plot of the final structure, 98.21% and 1.49% of the residues are in the favored regions and allowed regions, respectively.

### Cell culture

AsPc-1 (CRL-1682), SW1990 (CRL-2172), H1299 (or NCI-H1299, CRL-5803), HCT-116 (CCL-247), A549 (CRM-CCL-185), A375 (or A-375, CRL-1619), HEK-293 (CRL-1573) cells were obtained from ATCC and maintained in high-glucose (4.5 g/L) DMEM (Gibco 11995073) supplemented with 4 mM L-glutamine, 1 mM sodium pyruvate, and 10% heat-inactivated fetal bovine serum (Axenia Biologix). AGS (CRL-1739) cells were obtained from ATCC and maintained in Ham’s F12-K (Gibco 21127022) supplemented with 10% heat-inactivated fetal bovine serum (Axenia Biologix). Ba/F3 cells were a gift from Dr. Trevor Bivona (UCSF) and were maintained in RPMI-1640 (Gibco 11875093) supplemented with 10% heat-inactivated fetal bovine serum (Axenia Biologix) and 10 ng/mL recombinant mouse interleukin-3 (Gibco PMC0031).

Cells were passed for at least two generations after cryorecovery before they were used for assays. All cell lines were tested mycoplasma negative using MycoAlert™ Mycoplasma Detection Kit (Lonza).

### Gel electrophoresis and immunoblot

cells were treated with drugs at 40-60% confluency at a final DMSO concentration of 1%. At the end of treatment period, cells were chilled on ice. Unless otherwise indicated, adherent cells were washed once with ice-cold PBS (1 mL), scraped with a spatula, and pelleted by centrifugation (500 x g, 5 min). Suspension cells were pelleted by centrifugation (500 x g, 5 min), washed with 1 mL ice-cold PBS, and pelleted again. Cells were lysed in RIPA buffer supplemented with protease and phosphatase inhibitors (mini cOmplete and phosSTOP, Roche) on ice for 10 min.

Concentrations of lysates were determined with protein BCA assay (Thermo Fisher) and adjusted to 2 mg/mL or lowest available concentration with additional RIPA buffer. Samples were mixed with 5x SDS Loading Dye and denatured at either room temperature for 30 min for Ras band shift experiment, or 95 ºC for 5 min for other non-Ras proteins immunoblotting.

Unless otherwise noted, SDS-PAGE was run with Novex 12% Bis-Tris gel (Invitrogen) in MOPS running buffer (Invitrogen) at 200 V for 60 min following the manufacturer’s instructions. Protein bands were transferred onto 0.2-μm nitrocellulose membranes (Bio-Rad) using a wet-tank transfer apparatus (Bio-Rad Criterion Blotter) in 1x TOWBIN buffer with 10% methanol at 75V for 45 min. Membranes were blocked in 5% BSA–TBST for 1 h at 23 ºC. Primary antibody binding was performed with the indicated antibodies diluted in 5% BSA–TBST at 4 ºC for at least 16 h. After washing the membrane three times with TBST (5 min each wash), secondary antibodies (goat anti-rabbit IgG-IRDye 800 and goat anti-mouse IgG-IRDye 680, Li-COR) were added as solutions in 5% BSA–TBST at the dilutions recommended by the manufacturer. Secondary antibody binding was allowed to proceed for 1 h at 23 ºC. The membrane was washed three times with TBST (5 min each wash) and imaged on a Li-COR Odyssey fluorescence imager.

### Preparation of Mouse Stem Cell Virus (MSCV)

pMSCV-Puro plasmids containing full length human *KRAS* genes (wildtype, G12D) were constructed using standard molecule biology techniques by inserting the *KRAS* gene fragment between the BamHI and XhoI sites. Transfection-grade plasmids were prepared using ZymoPure II Plasmid Midiprep kit. EcoPack 293 cells (Takara Bio) were plated in 6-well plates (3 x 10^5^/mL, 2 mL). The next day, cells were transfected with 2.5 μg pMSCV plasmid using lipofectamine 3000 following the manufacturer’s instructions. The cells were incubated for 66 h, and then the virus-containing supernatants were collected and passed through a 0.22-μm syringe filter. The harvested virus was used immediately for spinfection of Ba/F3 cells or stored at –80 ºC.

### Generation of stable Ba/F3 transductants

1 mL of MSCV-containing supernatant (*vide supra*) was added to one well of a 6-well plate containing 1 x 10^6^ Ba/F3 cells in 1 mL of media comprised of 60% RMPI 1640, 40% heat-inactivated FBS, 10 ng mouse IL-3 and 4 μg polybrene. Cells were spinfected by centrifugation at 2,000 g for 90 minutes at room temperature and then placed in the incubator for 24 hours. After 1 day, the cells were diluted into 10 mL culture medium (RPMI 1640 + 10% heat-inactivated FBS, 10 ng/mL mouse IL-3) and recovered for a second day after spinfection. On the third day after spinfection, cells were pelleted at 500 x g for 5 min and resuspended in 10 mL selection medium (RPMI 1640 + 10% heat-inactivated FBS, 10 ng/mL mouse IL-3, 1.25 μg/mL puromycin). Cells were maintained under puromycin selection for 4-7 days, splitting as required to maintain density <2 x 10^6^ cells/mL. After 7 days, cells were pelleted, washed once with IL-3 free culture medium (RPMI 1640 + 10% heat-inactivated FBS) and pelleted again before resuspending at 2-4 x 10^5^ cells/mL in IL-3 free culture medium. Cells were maintained under these conditions for 7 days, passaging as needed to maintain density < 2 x 10^6^ cells/mL. Growth was monitored (Countess II Cell Counter) over these 7 days to confirm that an IL-3 independent population has been achieved.

### Differential scanning fluorimetry (DSF)

The protein of interest was diluted with SEC Buffer [20 mM HEPES 7.5, 150 mM NaCl, 1 mM MgCl_2_] to 2 μM. This solution was dispensed into wells of a white 96-well PCR plate in triplicate (25 μL/well). Fluorescence was measured at 0.5-ºC temperature intervals every 30 s from 25 ºC to 95 ºC on a Bio-Rad CFX96 qPCR system using the FRET setting. Each data set was normalized to the highest fluorescence and the normalized fluorescence reading was plotted against temperature in GraphPad Prism 9.0. *T*_m_ values were determined as the temperature(s) corresponding to the maximum(ma) of the first derivative of the curve. Proteins crosslinked with small molecules were desalted using Zeba™ Spin Desalting Columns (Thermo) before DSF *T*_m_ measurement.

### Detection of covalent modification of K-Ras by whole-protein mass spectrometry

Test compounds were prepared as 100x stock solutions in DMSO. K-Ras proteins were diluted with SEC Buffer (20 mM HEPES 7.5, 150 mM NaCl, 1 mM MgCl_2_) to 1 μM. In a typical reaction, 1 μL 100x compound stock was mixed with 99 μL diluted K-Ras protein, and the resulting mixture was incubated for the desired amount of time. The extent of modification was assessed by electrospray mass spectrometry using a Waters Xevo G2-XS system equipped with an Acquity UPLC BEH C4 1.7 μm column. The mobile phase was a linear gradient of 5–95% acetonitrile/water + 0.05% formic acid. For kinetic measurements, a 2x compound solution was first prepared in SEC Buffer, which was then mixed with 400 nM K-Ras(G12D) protein at 1:1 (v/v) ratio. Injection time stamps were used to calculate elapsed time.

### Detection of covalent modification of K-Ras by tandem mass spectrometry

K-Ras(G12D) protein (1 μM, 100 μL) in PBS 7.4 was treated with β-propiolactone (1 mM or 10 mM) at 23 ºC for 1 h. The reaction buffer was exchanged into Digestion Buffer (20 mM Tris 8.0, 2 mM CaCl_2_) using a Zeba 0.5-mL desalting column (7K MWCO, Thermo Scientific). 80 μL of the resulting protein solution was mixed with 2 μL 200 mM DTT. The mixture was heated at 56 ºC for 30 min. After cooling to 23 ºC, 4 μL 200 mM iodoacetamide was added. After 15 min at 23 ºC, 2.1 μL 200 mM DTT was added. After an additional 5 min, 500 ng trypsin was added to the mixture, and the samples were incubated at 37 ºC overnight. 5 μL 10% formic acid was added to stop the digestion (final formic acid concentration: 0.5% v/v). The tryptic peptides were enriched and desalted using OMIX C18 tips (Agilent) following the manufacturer’s instructions. Peptides (0.5% of total) were resolved on an Easy-Spray nano-HPLC column (Thermo Fisher ES800A, 150 mm length, 3 μL particle size, 100-Å particle size) over a 54-min gradient of 2–37% acetonitrile–water + 0.1% formic acid and analyzed by a Q-Exactive hybrid quadrupole-orbitrap mass spectrometer (MS1 resolution 70,000, AGC target: 3e6, range: 350-1500 m/z; MS2 resolution 17,500, AGC target: 5e4, max injection time: 120 ms, Top 10, NCE: 25%, dynamic exclusion: 15 s). Peptides were searched against the K-Ras(G12D) sequence using MaxQuant (v.2.0.3.0, https://www.maxquant.org/), with β-propiolactone (C_3_H_4_O_2_) as a variable modification on serine, threonine, lysine, aspartate, tyrosine, glutamate and histidine residues. Peptides were identified with a false discovery rate cutoff of 0.01. Only the peptides with sufficient MS2 fragment information to assign the modification site with >0.9 probability were used for analysis.

### Stability of β-lactone compounds in PBS

10 μM solution of β-lactone compound in PBS, pH 7.4 was prepared by diluting 1 μl of 10 mM DMSO stock solution into 999 μl of PBS pH 7.4 in the presence or absence of 5 mM reduced glutathione (GSH). At specified time points, aliquots were taken, and the amount of intact compound was analyzed on a Waters Xevo G2-XS Quadrupole-TOF system equipped with an Acquity UPLC BEH C18 1.7 μm column for multiple-reaction monitoring (MRM) for the 1+ precursor of respective compound (precursor > precursor) with target enhancement and low collision energy (0–2 eV).

### 2D cell viability assay

Cells were seeded into 96-well white flat bottom plates (1,000 cells/well) (Corning) and incubated overnight. Cells were treated with the indicated compounds in a nine-point threefold dilution series (100 μL final volume) and incubated for 72 h. Cell viability was assessed using a commercial CellTiter-Glo (CTG) luminescence-based assay (Promega). The 96-well plates were equilibrated to room temperature before the addition of diluted CTG reagent (100 μL) (1:4 CTG reagent:PBS, containing 1% Triton X-100). Plates were placed on an orbital shaker for 30 min before recording luminescence using a Spark 20M (Tecan) plate reader.

### 3D cell viability assay

Cells were resuspended in fresh culture medium to a concentration of (1,000 cells/well) and plated (90 μL/well) in Corning Costar # 3474, 96-well Clear Flat Bottom Ultra-Low Attachment Microplate. Cells were treated with the indicated compounds in a nine-point threefold dilution series (100 μL final volume) and incubated for 120 h^39^. Cell viability was assessed using a commercial CellTiter-Glo (CTG) luminescence-based assay (Promega) as described in 2D cell viability assay.

## Data availability

Atomic coordinates and structure factors for the reported crystal structures have been deposited with the Protein Data Bank (PDB), with the following accession numbers: K-Ras(G12D)•GDP•1, 8T4V. Source data of uncropped, unprocessed gel images are provided with this paper.

## Acknowledgements

Q.Z. is the Connie and Bob Lurie Fellow of the Damon Runyon Cancer Research Foundation (DRG-2434-21). Z.Z. is a Damon Runyon Fellow supported by the Damon Runyon Cancer Research Foundation (DRG-2281-17). K.Z.G. was supported as an HHMI Fellow by the Damon Runyon Cancer Research Foundation (DRG-2399-20). K.M.S. would like to acknowledge support from HHMI. We would like to thank J. Taunton and Z. Knight for helpful advice.

## Author contributions

Q.Z., Z.Z., and K.M.S. conceptualized this study, wrote the original draft, reviewed and edited the manuscript. Q.Z., and Z.Z. synthesized, characterized, tested compounds in biochemical and cell-based assays, and analyzed data. Z.Z., and K.Z.G. collected and analyzed crystallization data.

## Competing interests

K.M.S., Z.Z., and Q.Z. are inventors on patents related to covalent K-Ras(G12D) inhibitors reported here. K.M.S. is an inventor on patents owned by University of California San Francisco covering KRAS targeting small molecules licensed to Araxes, Erasca and Novartis. K.M.S. has consulting agreements for the following companies, which involve monetary and/or stock compensation: Revolution Medicines, Black Diamond Therapeutics, BridGene Biosciences, Denali Therapeutics, Dice Molecules, eFFECTOR Therapeutics, Erasca, Novartis, Genentech/Roche, Janssen Pharmaceuticals, Kumquat Biosciences, Kura Oncology, Mitokinin, Type6 Therapeutics, Nested Therapeutics, Vevo, Wellspring Biosciences (Araxes Pharma), Ikena, Initial Therapeutics and BioTheryX.

## Materials & Correspondence

Correspondence and material requests should be addressed to K.M.S. and Z.Z.

## Notes

### Summary of Updates

Figure 3 updated

## Reference

1 Prior, I. A., Hood, F. E. & Hartley, J. L. The Frequency of Ras Mutations in Cancer. Cancer Res 80, 2969–2974 (2020). 10.1158/0008-5472.CAN-19-3682

2 Ostrem, J. M., Peters, U., Sos, M. L., Wells, J. A. & Shokat, K. M. K-Ras(G12C) inhibitors allosterically control GTP affinity and effector interactions. Nature 503, 548–551 (2013). 10.1038/nature12796

3 Hallin, J. et al. Anti-tumor efficacy of a potent and selective non-covalent KRASG12D inhibitor. Nature Medicine 28, 2171–2182 (2022). 10.1038/s41591-022-02007-7

4 Kim, D. et al. Pan-KRAS inhibitor disables oncogenic signalling and tumour growth. Nature (2023). 10.1038/s41586-023-06123-3

5 Ma, N. et al. 2 H -Azirine-Based Reagents for Chemoselective Bioconjugation at Carboxyl Residues Inside Live Cells. Journal of the American Chemical Society 142, 6051–6059 (2020). 10.1021/jacs.9b12116

6 McGrath, N. A., Andersen, K. A., Davis, A. K. F., Lomax, J. E. & Raines, R. T. Diazo compounds for the bioreversible esterification of proteins. Chemical Science 6, 752–755 (2015). 10.1039/c4sc01768d

7 Jun, J. V., Petri, Y. D., Erickson, L. W. & Raines, R. T. Modular Diazo Compound for the Bioreversible Late-Stage Modification of Proteins. Journal of the American Chemical Society 145, 6615–6621 (2023). 10.1021/jacs.2c11325

8 Knox, J. E. et al. in American Association for Cancer Research (AACR) Annual Meeting.

9 Zanon, P. R. A. et al. Profiling the Proteome-Wide Selectivity of Diverse Electrophiles. (2021). 10.26434/chemrxiv.14186561.v1

10 Abbasov, M. E. et al. A proteome-wide atlas of lysine-reactive chemistry. Nat Chem 13, 1081–1092 (2021). 10.1038/s41557-021-00765-4

11 Yang, T. et al. Reversible lysine-targeted probes reveal residence time-based kinase selectivity. Nat Chem Biol 18, 934–941 (2022). 10.1038/s41589-022-01019-1

12 Wan, X. et al. Discovery of Lysine-Targeted eIF4E Inhibitors through Covalent Docking. Journal of the American Chemical Society 142, 4960–4964 (2020). 10.1021/jacs.9b10377

13 Chen, W. et al. Arylfluorosulfates Inactivate Intracellular Lipid Binding Protein(s) through Chemoselective SuFEx Reaction with a Binding Site Tyr Residue. J Am Chem Soc 138, 7353–7364 (2016). 10.1021/jacs.6b02960

14 Zhang, Z., Guiley, K. Z. & Shokat, K. M. Chemical acylation of an acquired serine suppresses oncogenic signaling of K-Ras(G12S). Nat Chem Biol 18, 1177–1183 (2022). 10.1038/s41589-022-01065-9

15 Zheng, Q. et al. SuFEx-enabled, agnostic discovery of covalent inhibitors of human neutrophil elastase. Proceedings of the National Academy of Sciences 116, 18808–18814 (2019). 10.1073/pnas.1909972116

16 Zhang, Z., Morstein, J., Ecker, A. K., Guiley, K. Z. & Shokat, K. M. Chemoselective Covalent Modification of K-Ras(G12R) with a Small Molecule Electrophile. J Am Chem Soc 144, 15916–15921 (2022). 10.1021/jacs.2c05377

17 Gonzalez-Valero, A. et al. An Activity-Based Oxaziridine Platform for Identifying and Developing Covalent Ligands for Functional Allosteric Methionine Sites: Redox-Dependent Inhibition of Cyclin-Dependent Kinase 4. Journal of the American Chemical Society 144, 22890–22901 (2022). 10.1021/jacs.2c04039

18 Lowther, W. T., McMillen, D. A., Orville, A. M. & Matthews, B. W. The anti-angiogenic agent fumagillin covalently modifies a conserved active-site histidine in the <i>Escherichia coli</i> methionine aminopeptidase. Proceedings of the National Academy of Sciences 95, 12153–12157 (1998). 10.1073/pnas.95.21.12153

19 Jia, S., He, D. & Chang, C. J. Bioinspired Thiophosphorodichloridate Reagents for Chemoselective Histidine Bioconjugation. Journal of the American Chemical Society 141, 7294–7301 (2019). 10.1021/jacs.8b11912

20 Cruite, J. T. et al. Cereblon covalent modulation through structure-based design of histidine targeting chemical probes. RSC Chemical Biology 3, 1105–1110 (2022). 10.1039/D2CB00078D

21 McGregor, L. M., Jenkins, M. L., Kerwin, C., Burke, J. E. & Shokat, K. M. Expanding the Scope of Electrophiles Capable of Targeting K-Ras Oncogenes. Biochemistry 56, 3178–3183 (2017). 10.1021/acs.biochem.7b00271

22 Wang, H.-L., Cee, V. J., Parsons, A. T. & Beaver, M. Pyridopyrimidine Derivatives Useful as KRAS G12C and KRAS G12D Inhibitors in the Treatment of Cancer. WO 2021/081212 A1 (2021).

23 Yu, Z. et al. Simultaneous Covalent Modification of K-Ras(G12D) and K-Ras(G12C) with Tunable Oxirane Electrophiles. Journal of the American Chemical Society (2023). 10.1021/jacs.3c05899

24 Robinson, S. L., Christenson, J. K. & Wackett, L. P. Biosynthesis and chemical diversity of β-lactone natural products. Natural Product Reports 36, 458–475 (2019). 10.1039/C8NP00052B

25 Hassan, A. Q. et al. The novolactone natural product disrupts the allosteric regulation of Hsp70. Chem Biol 22, 87–97 (2015). 10.1016/j.chembiol.2014.11.007

26 She, Y. M., Cheng, K. D., Farnsworth, A., Li, X. G. & Cyr, T. D. Surface modifications of influenza proteins upon virus inactivation by beta-propiolactone. Proteomics 13, 3537–3547 (2013). 10.1002/pmic.201300096

27 Logrippo, G. A. Investigations of the use of beta-propiolactone in virus inactivation. Ann N Y Acad Sci 83, 578–594 (1960). 10.1111/j.1749-6632.1960.tb40931.x

28 Budowsky, E. I., Friedman, E. A., Zheleznova, N. V. & Noskov, F. S. Principles of selective inactivation of viral genome. VI. Inactivation of the infectivity of the influenza virus by the action of beta-propiolactone. Vaccine 9, 398–402 (1991). 10.1016/0264-410x(91)90125-p

29 Fan, C. et al. Beta-Propiolactone Inactivation of Coxsackievirus A16 Induces Structural Alteration and Surface Modification of Viral Capsids. J Virol 91 (2017). 10.1128/JVI.00038-17

30 Gao, Q. et al. Development of an inactivated vaccine candidate for SARS-CoV-2. Science 369, 77–81 (2020). 10.1126/science.abc1932

31 Wang, H. et al. Development of an Inactivated Vaccine Candidate, BBIBP-CorV, with Potent Protection against SARS-CoV-2. Cell 182, 713–721 e719 (2020). 10.1016/j.cell.2020.06.008

32 Böttcher, T. & Sieber, S. A. β-Lactams and β-lactones as activity-based probes in chemical biology. MedChemComm 3, 408–417 (2012). 10.1039/C2MD00275B

33 Zhang, Y., Gross, R. A. & Lenz, R. W. Stereochemistry of the ring-opening polymerization of (S)-.beta.-butyrolactone. Macromolecules 23, 3206–3212 (1990). 10.1021/ma00215a002

34 Hansen, R. et al. The reactivity-driven biochemical mechanism of covalent KRAS(G12C) inhibitors. Nat Struct Mol Biol 25, 454–462 (2018). 10.1038/s41594-018-0061-5

35 Wang, X. et al. KRAS G12D Inhibitors. WO/2021/041671 (2021).

36 Vasta, J. D. et al. KRAS is vulnerable to reversible switch-II pocket engagement in cells. Nature Chemical Biology 18, 596–+ (2022). 10.1038/s41589-022-00985-w

37 Wang, X. et al. Identification of MRTX1133, a Noncovalent, Potent, and Selective KRAS(G12D) Inhibitor. J Med Chem 65, 3123–3133 (2022). 10.1021/acs.jmedchem.1c01688

38 Miles, A. K., John, J. W. H., Yeon-Hwa, L., Chih-Shia, L. & Ji, L. A non-conserved histidine residue on KRAS drives paralog selectivity of the KRAS G12D inhibitor MRTX1133. bioRxiv, 2022.2012.2016.520846 (2022). 10.1101/2022.12.16.520846

39 Janes, M. R. et al. Targeting KRAS Mutant Cancers with a Covalent G12C-Specific Inhibitor. Cell 172, 578–589 e517 (2018). 10.1016/j.cell.2018.01.006

40 Lanman, B. A. et al. Discovery of a Covalent Inhibitor of KRAS(G12C) (AMG 510) for the Treatment of Solid Tumors. J Med Chem 63, 52–65 (2020). 10.1021/acs.jmedchem.9b01180

41 Fell, J. B. et al. Identification of the Clinical Development Candidate MRTX849, a Covalent KRAS(G12C) Inhibitor for the Treatment of Cancer. J Med Chem 63, 6679–6693 (2020). 10.1021/acs.jmedchem.9b02052

42 Lorthiois, E. et al. JDQ443, a Structurally Novel, Pyrazole-Based, Covalent Inhibitor of KRASG12C for the Treatment of Solid Tumors. Journal of Medicinal Chemistry 65, 16173–16203 (2022). 10.1021/acs.jmedchem.2c01438

43 Schulze, C. J. et al. Chemical remodeling of a cellular chaperone to target the active state of mutant KRAS. Science 381, 794–799 (2023). 10.1126/science.adg9652

44 Zhang, Z. et al. GTP-State-Selective Cyclic Peptide Ligands of K-Ras(G12D) Block Its Interaction with Raf. ACS Cent Sci 6, 1753–1761 (2020). 10.1021/acscentsci.0c00514

45 Hunter, J. C. et al. Biochemical and Structural Analysis of Common Cancer-Associated KRAS Mutations. Mol Cancer Res 13, 1325–1335 (2015). 10.1158/1541-7786.MCR-15-0203

46 Olson, A. R. & Miller, R. J. The Mechanism of the Aqueous Hydrolysis of β-Butyrolactone. Journal of the American Chemical Society 60, 2687–2692 (1938). 10.1021/ja01278a041

47 Fujioka, A. et al. Dynamics of the Ras/ERK MAPK Cascade as Monitored by Fluorescent Probes*. Journal of Biological Chemistry 281, 8917–8926 (2006). 10.1074/jbc.M509344200

48 Peacock, D. M., Kelly, M. J. S. & Shokat, K. M. Probing the KRas Switch II Groove by Fluorine NMR Spectroscopy. ACS Chem Biol 17, 2710–2715 (2022). 10.1021/acschembio.2c00566

49 Asai, A., Hasegawa, A., Ochiai, K., Yamashita, Y. & Mizukami, T. Belactosin A, a novel antitumor antibiotic acting on cyclin/CDK mediated cell cycle regulation, produced by Streptomyces sp. J Antibiot (Tokyo) 53, 81–83 (2000). 10.7164/antibiotics.53.81

50 Kawamura, S. et al. Potent proteasome inhibitors derived from the unnatural cis-cyclopropane isomer of Belactosin A: synthesis, biological activity, and mode of action. J Med Chem 56, 3689–3700 (2013). 10.1021/jm4002296

51 Kawamura, S., Unno, Y., Asai, A., Arisawa, M. & Shuto, S. Design and synthesis of the stabilized analogs of belactosin A with the unnatural cis-cyclopropane structure. Org Biomol Chem 11, 6615–6622 (2013). 10.1039/c3ob41338a

52 Battye, T. G., Kontogiannis, L., Johnson, O., Powell, H. R. & Leslie, A. G. iMOSFLM: a new graphical interface for diffraction-image processing with MOSFLM. Acta Crystallogr D Biol Crystallogr 67, 271–281 (2011). 10.1107/S0907444910048675

53 Evans, P. Scaling and assessment of data quality. Acta Crystallogr D Biol Crystallogr 62, 72–82 (2006). 10.1107/S0907444905036693

54 McCoy, A. J. et al. Phaser crystallographic software. J Appl Crystallogr 40, 658–674 (2007). 10.1107/S0021889807021206

55 Winn, M. D. et al. Overview of the CCP4 suite and current developments. Acta Crystallogr D Biol Crystallogr 67, 235–242 (2011). 10.1107/S0907444910045749

56 Emsley, P., Lohkamp, B., Scott, W. G. & Cowtan, K. Features and development of Coot. Acta Crystallogr D Biol Crystallogr 66, 486–501 (2010). 10.1107/S0907444910007493

57 Adams, P. D. et al. PHENIX: a comprehensive Python-based system for macromolecular structure solution. Acta Crystallogr D Biol Crystallogr 66, 213–221 (2010). 10.1107/S0907444909052925

