## Supplementary Information for "Strain-release alkylation of Asp12 enables mutant selective GTP-state targeting of K-Ras(G12D)"

The PDF file includes:

Supplementary Figures 1 to 12
Supplementary Tables 1 to 10
Chemical Synthesis
Supplementary Reference (58–63)
NMR Spectra

### Supplementary Figures and Tables

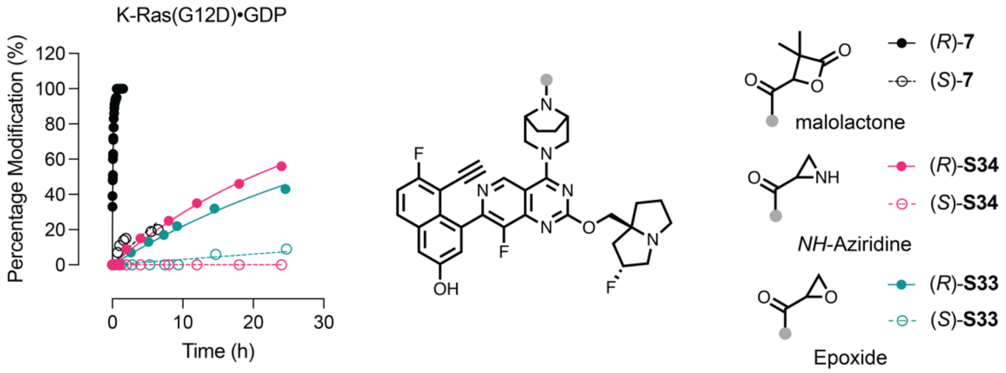


**Supplementary Figure 1**. Labeling kinetics of K-Ras(G12D) by strain-release electrophiles.

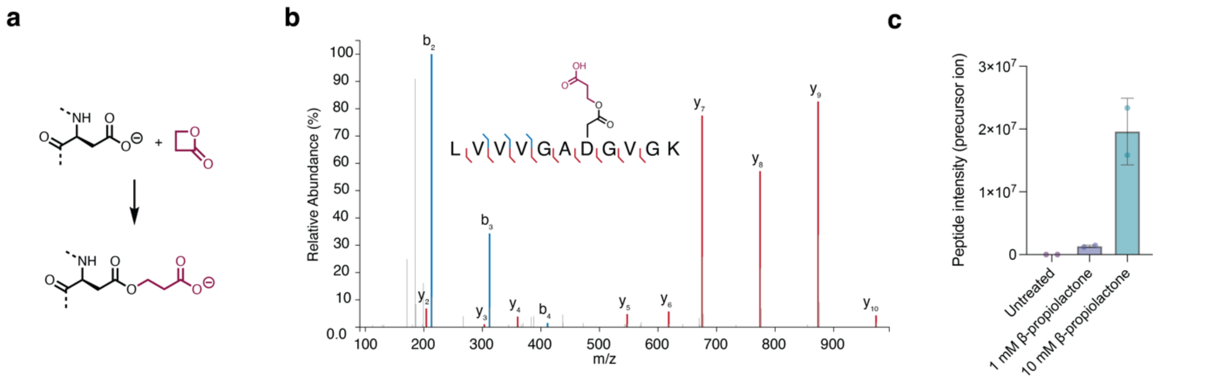


**Supplementary Figure 2**. β-Propiolactone covalently labeled Asp12 of recombinant K-Ras(G12D). **a**, Alkylation of Asp12 by β-propiolactone. **b**, Identified fragment ions of aa. 6-16 peptide including covalently modified Asp12. **c**, Extent of covalent modification is dose dependent.

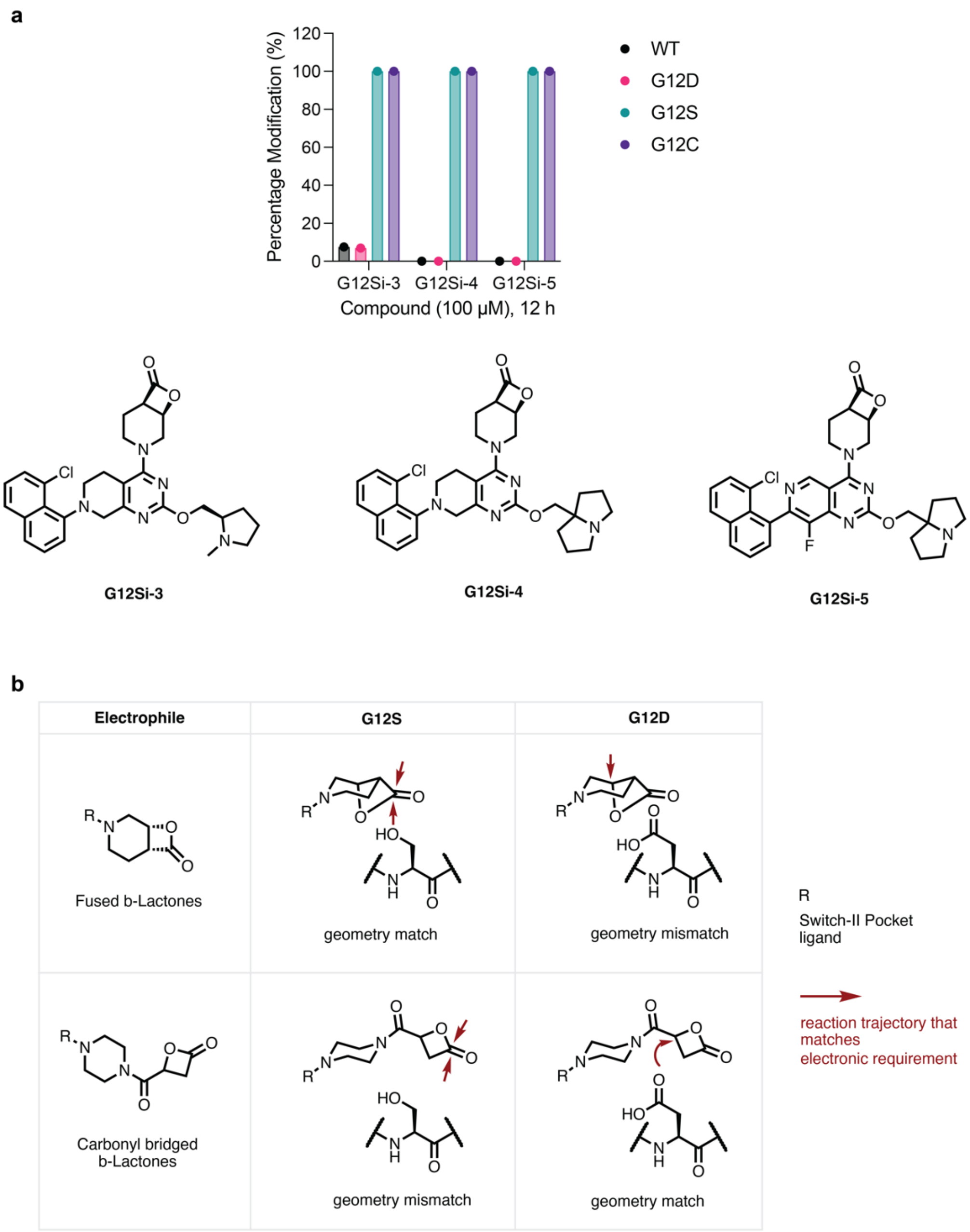


**Supplementary Figure 3**. Tuning the reactivity of β-lactone electrophile toward K-Ras(G12D) by introducing a carbonyl bridge. **a**, [4.2.0]-Fused β-lactone inhibitors covalently engage K-Ras(G12S), but not G12D. **b**, Mechanistic rationale of a carbonyl bridged β-lactone for K-Ras(G12D) covalent engagement.

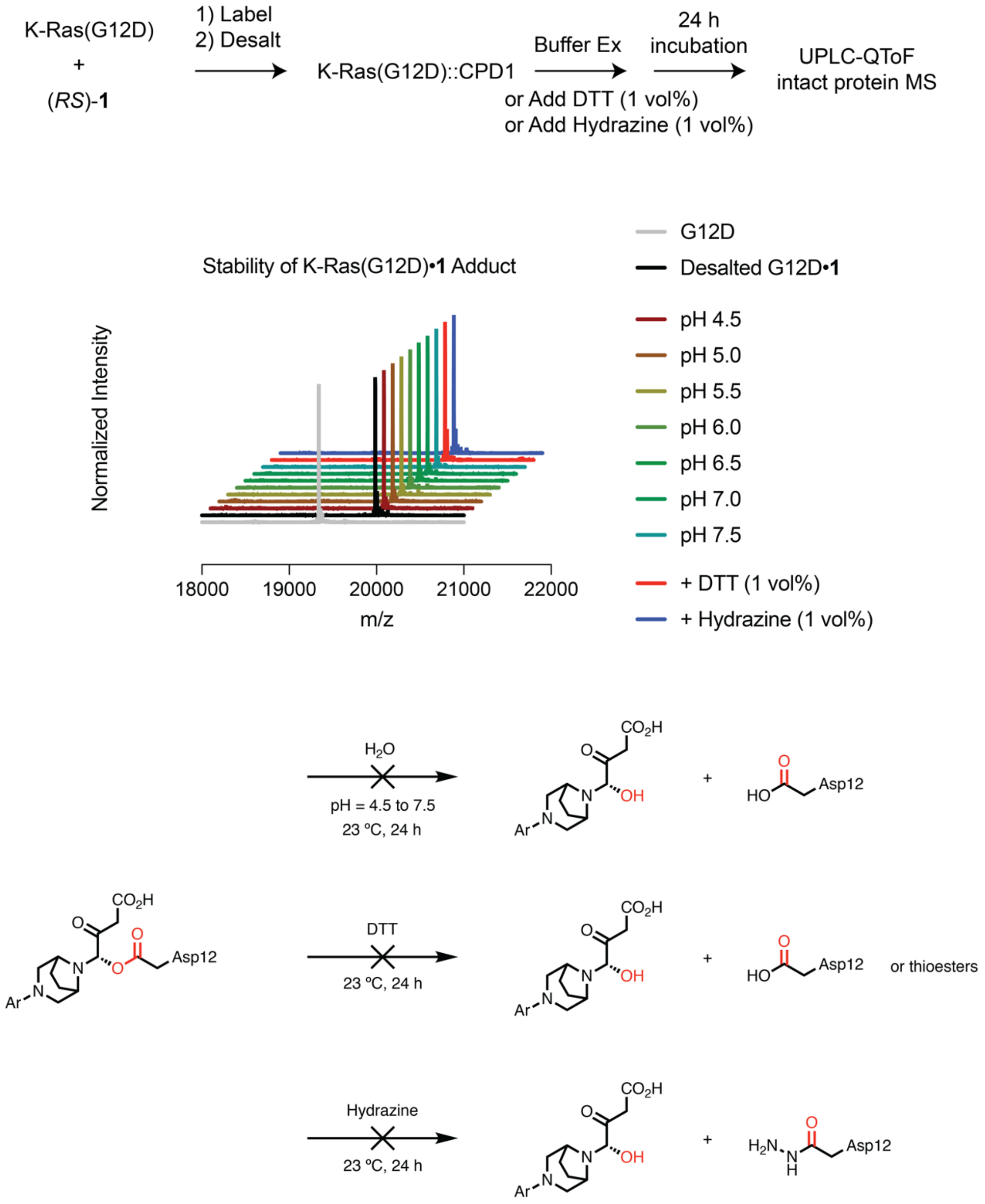


**Supplementary Figure 4**. Covalent K-Ras(G12D)•1 adduct is stable in pH 4.5–7.5 buffers and to thiol (DTT) or amine (hydrazine) electrophiles.

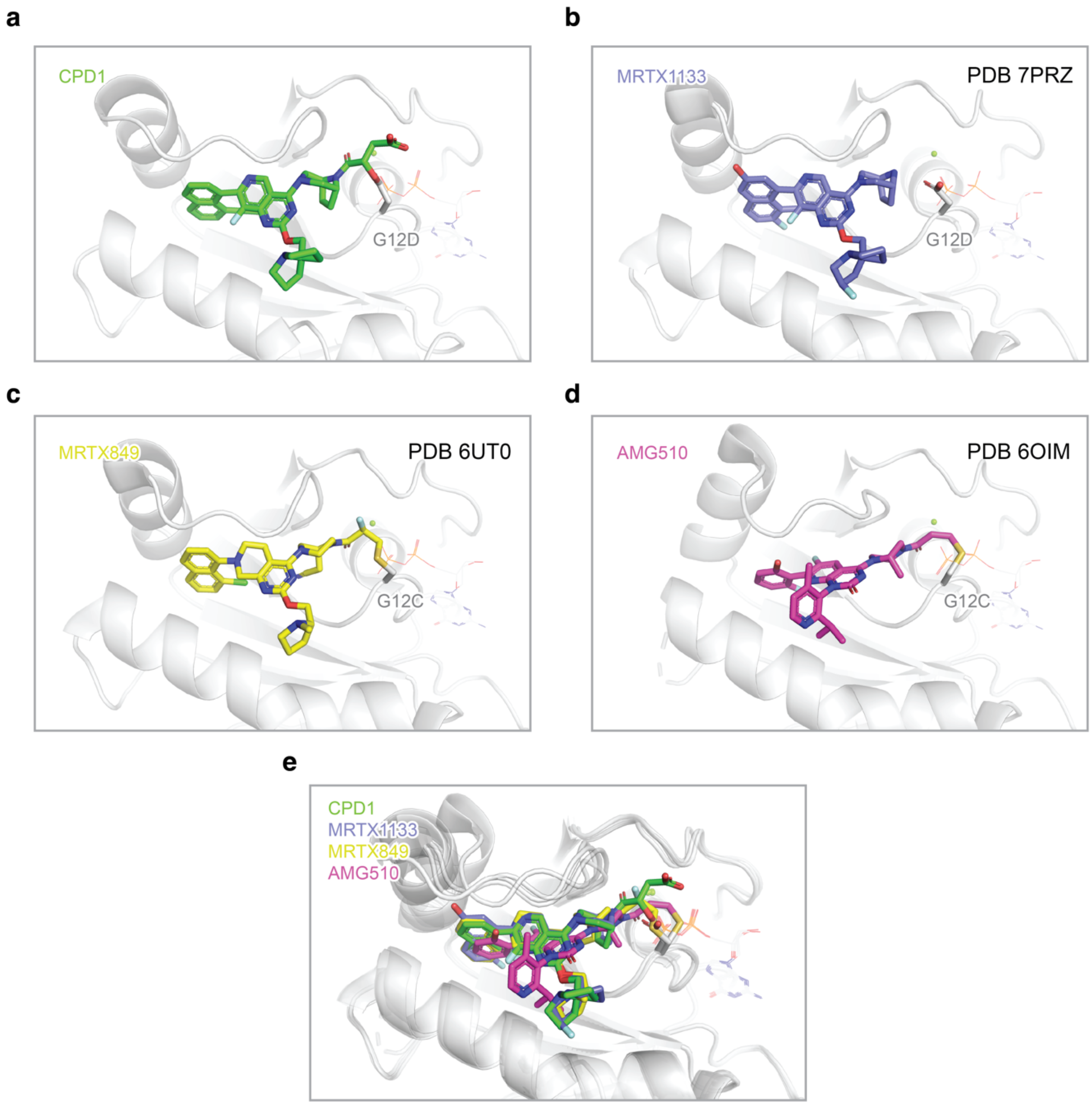


**Supplementary Figure 5**. Ligand poses in switch-II pocket of K-Ras proteins as shown in G12D•1 (**a**), G12D•MRTX1133 (**b**), G12C•MRTX849 (**c**), G12C•AMG510 (**d**), overlay (**e**).

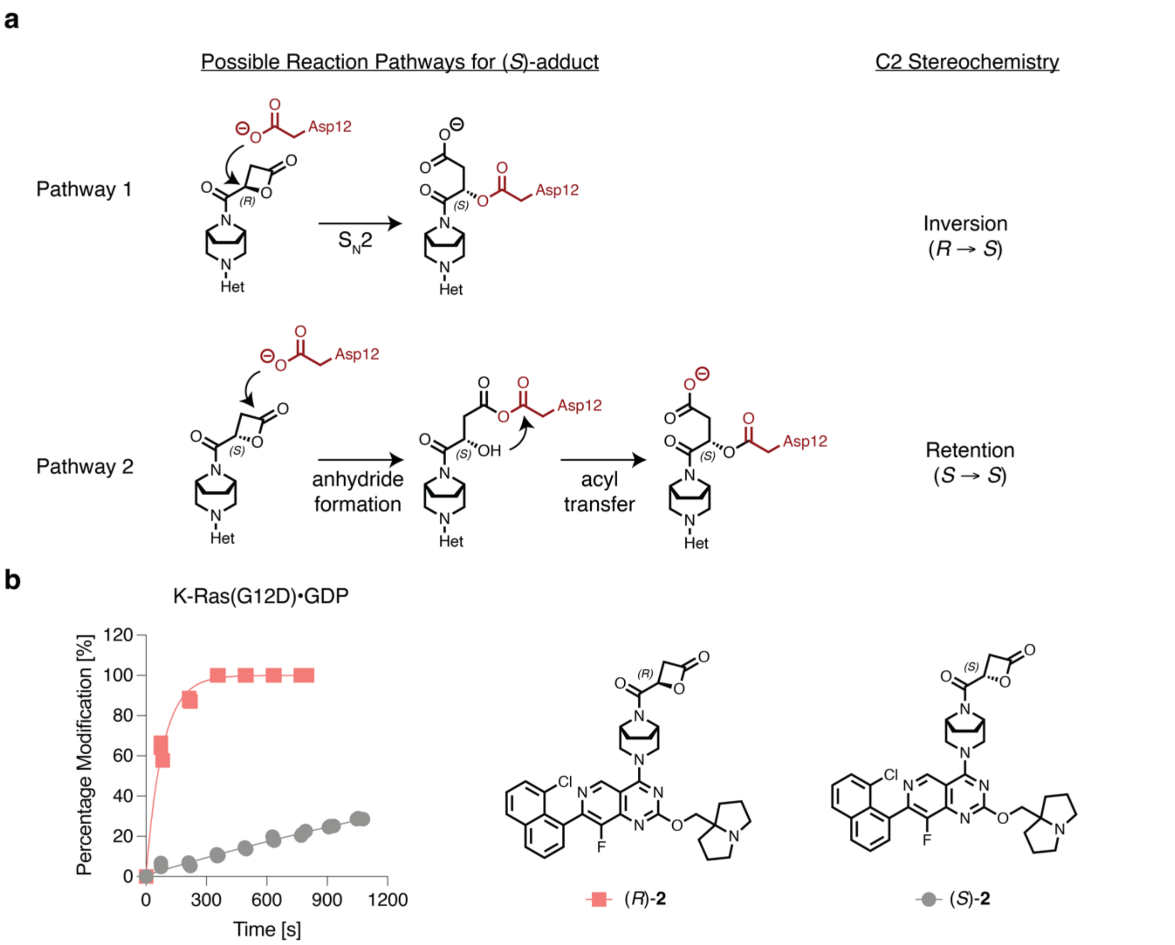


**Supplementary Figure 6**. Mechanistic rationale of the enantioselectivity of the covalent modification reaction. **a**, Two possible reaction pathways to form (*S*)-adduct with distinct stereochemical events at α-carbon. **b**, K-Ras(G12D) favors an (*R*)-enantiomer of the malolactone electrophile.

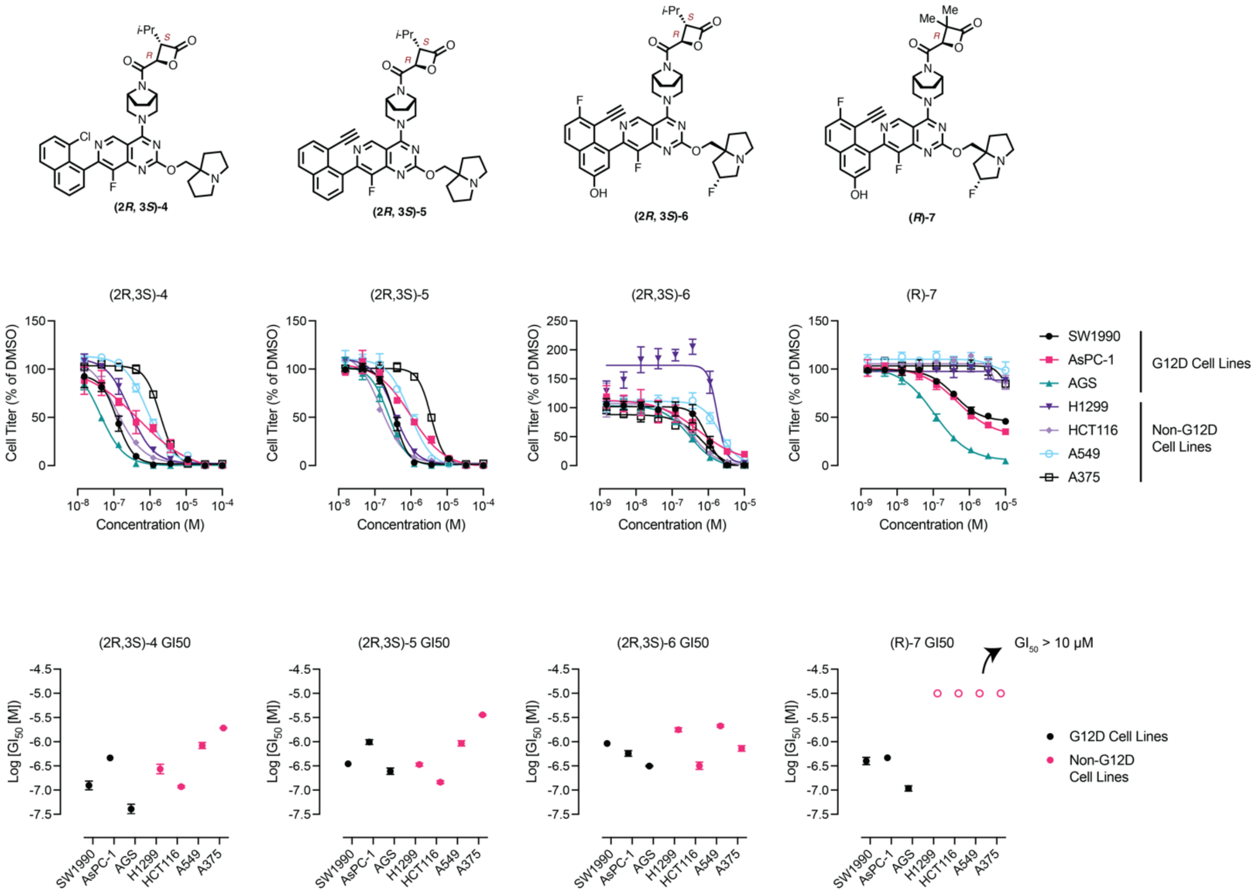


**Supplementary Figure 7**. Cell growth inhibition of advanced β-lactone G12Di against a panel of G12D or non-G12D cell lines.

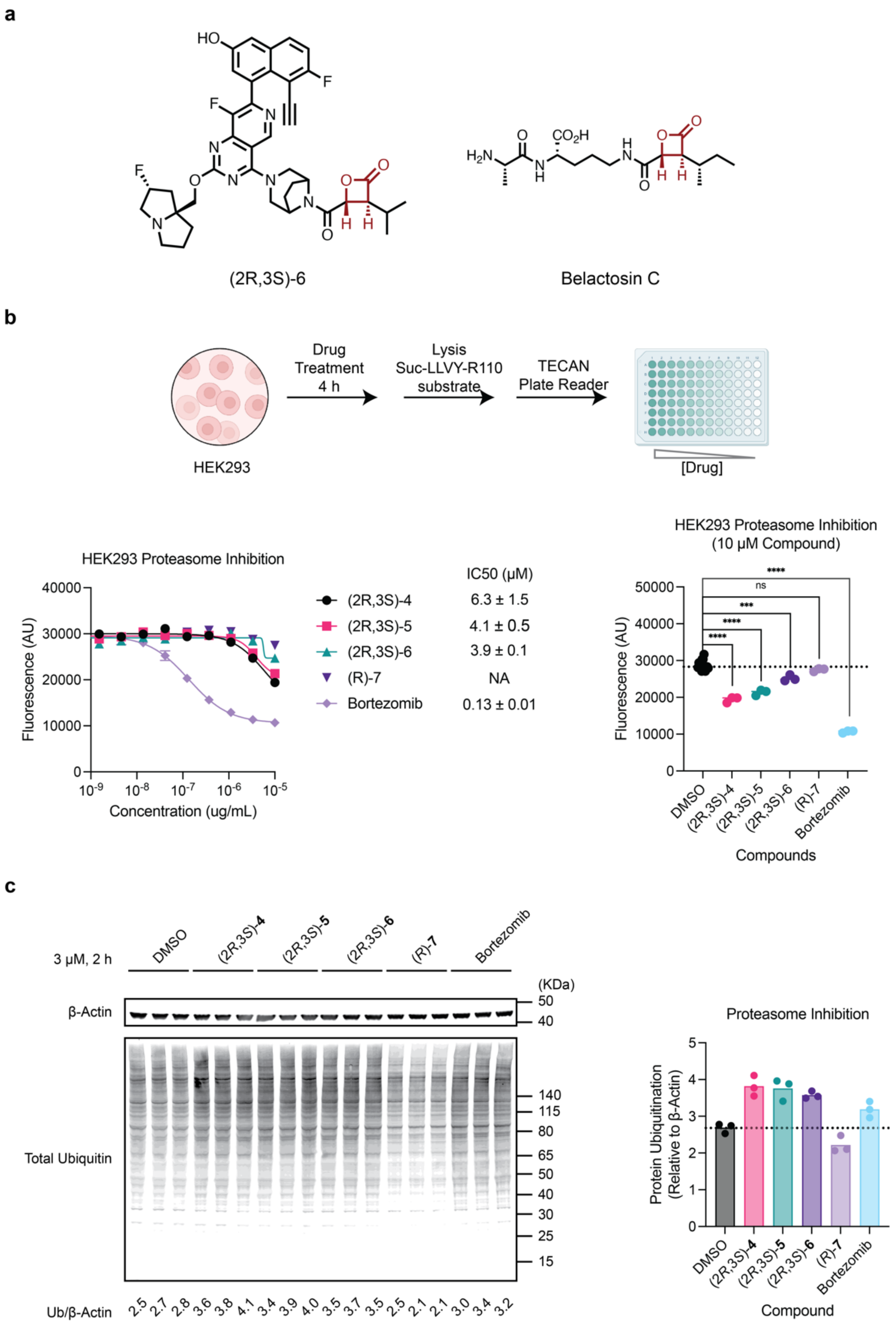


**Supplementary Figure 8**. Malolactone compounds 4–6 inhibited 20S proteasome while (*R*)-7 did not. **a**, Chemical structures of Compound 6 and Belactosin C with β-lactone warhead highlighted in red. **b**, Fluorescent-based proteasome activity assay (n = 3). **c**, Immunoblot of poly-ubiquitinated proteins in HEK293 cells (n = 3).

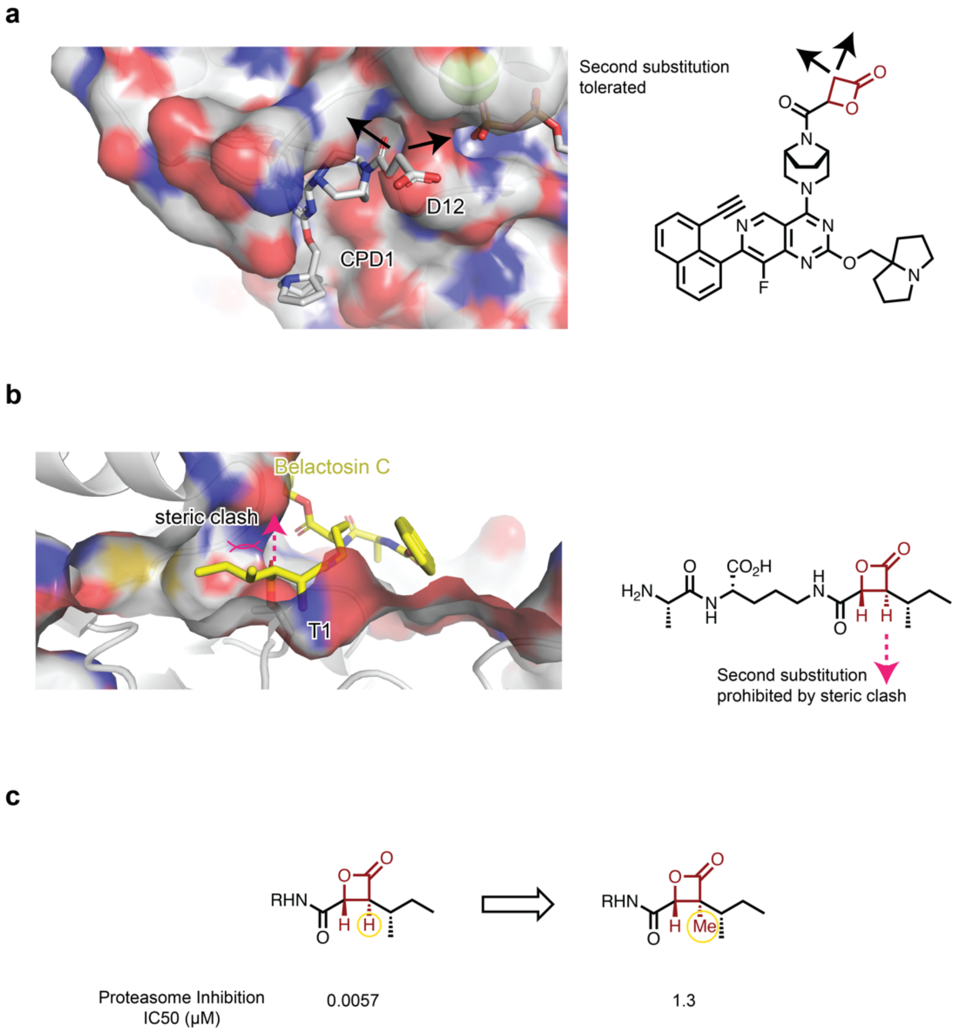


**Supplementary Figure 9**. Analysis of X-ray co-crystal structures containing covalent β-lactone warhead. **a**, α-Carbon of β-lactone G12Di could tolerate geminal disubstitution. **b**, Belactosin C could not tolerate a second α-substitution. **c**, Additional methyl group (circled in yellow) in belactosin analog caused significant loss of activity.

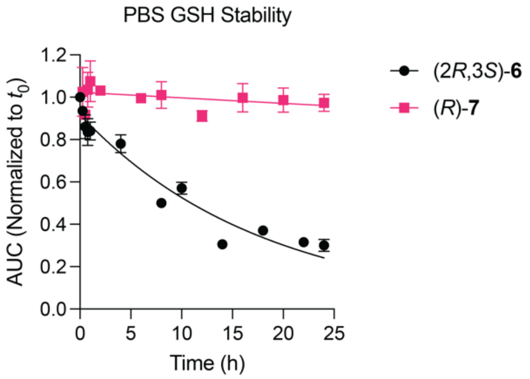


**Supplementary Figure 10**. Stability of Compounds **6** and **7** in buffered solution with reduced glutathione.

**a**

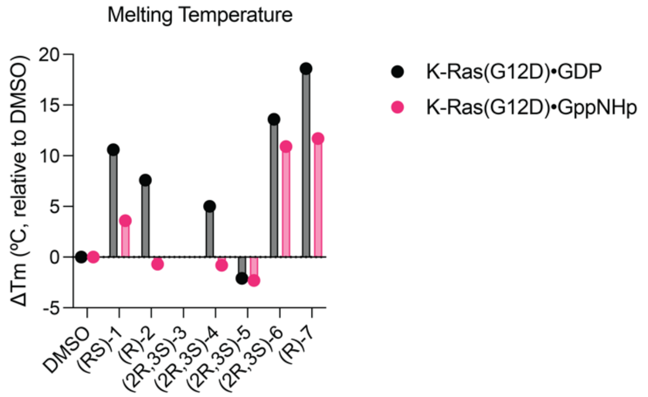


**b**

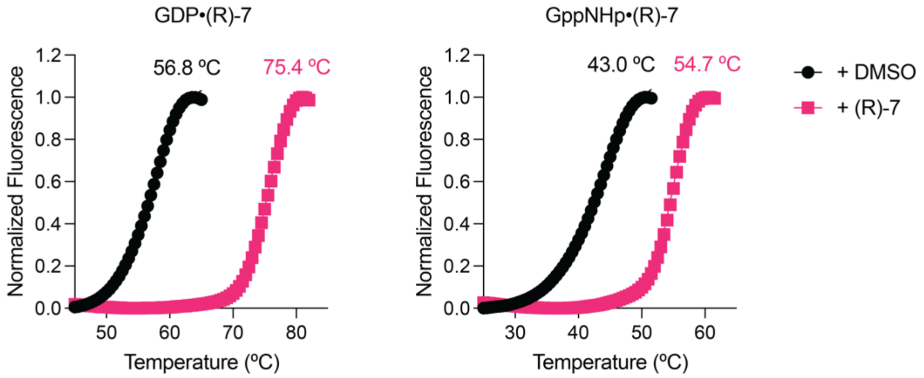


**Supplementary Figure 11**. Melting point change of covalently crosslinked K-Ras(G12D). **a**, Summary of thermal stabilization of K-Ras(G12D) by Compounds **1–7**. **b**, DSF melting curves of K-Ras(G12D)•GDP or GppNHp covalently bound to (*R*)-**7**.

**a**

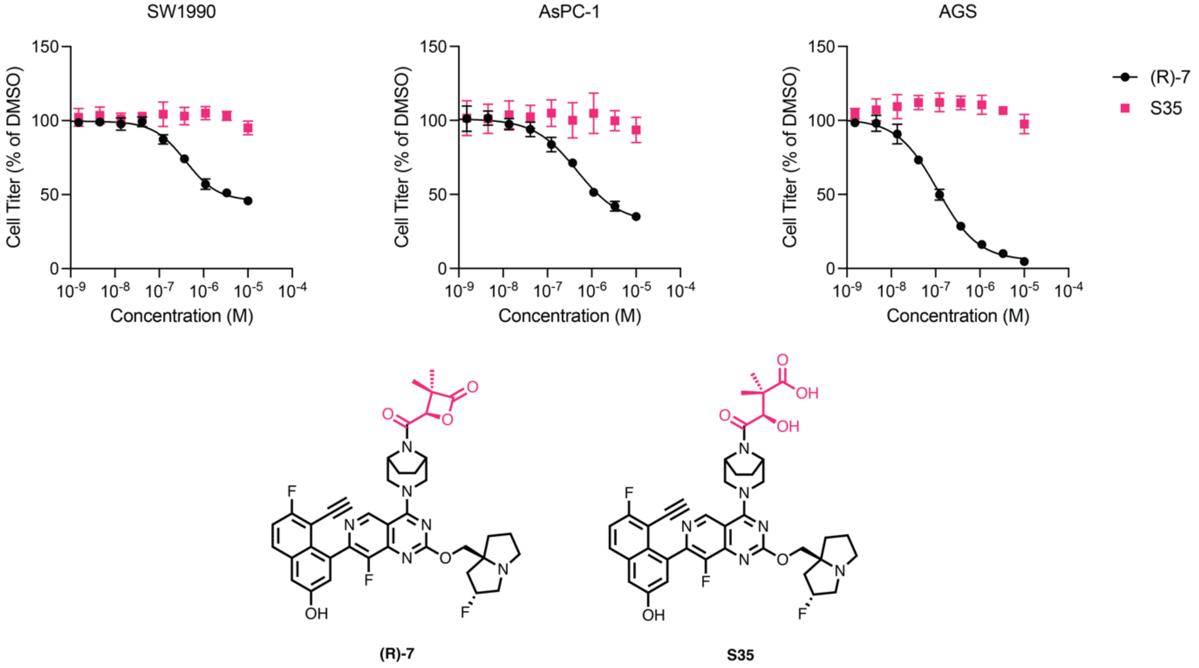


**b**

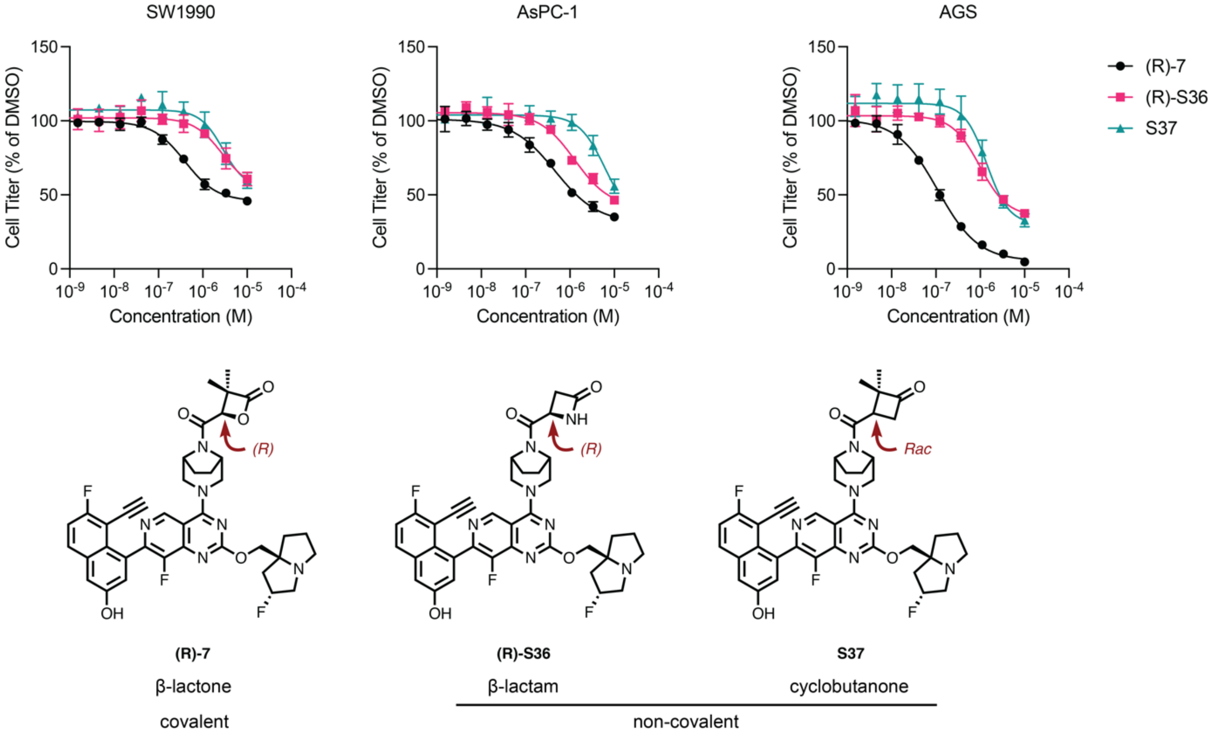


**Supplementary Figure 12**. Covalency of β-lactone compounds is important for the G12D cell growth inhibition. **a**, Hydrolyzed product did not inhibit K-Ras(G12D) cell lines. **b**, Non-covalent, structural analogs inhibited K-Ras(G12D) cell lines with significantly lower potency.

**Supplementary Table 1**. X-ray crystallography data collection and refinement statistics.

| | K-Ras(G12D)•GDP•**1** |
|---|---|
| Resolution range | 40.45 - 1.47 (1.523 - 1.47) |
| Space group | P 1 $2_1$ 1 |
| Cell dimensions | |
| *a*, *b*, *c* (Å) | 41.3198 65.8399 60.8498 |
| *α, β, γ* (º) | 90 101.75 90 |
| Total reflections | 355501 (28279) |
| Unique reflections | 53915 (5253) |
| Multiplicity | 6.6 (5.4) |
| Completeness (%) | 99.15 (96.64) |
| *I* / σ*I* | 12.52 (1.78) |
| Wilson B-factor | 18.15 |
| R-merge | 0.0779 (0.95) |
| R-meas | 0.08448 (1.054) |
| R-pim | 0.03226 (0.447) |
| CC1/2 | 0.998 (0.707) |
| CC* | 1 (0.91) |
| Reflections used in refinement | 53854 (5213) |
| Reflections used for R-free | 2728 (246) |
| R-work | 0.1892 (0.3244) |
| R-free | 0.2106 (0.3448) |
| CC(work) | 0.965 (0.818) |
| CC(free) | 0.952 (0.790) |
| Number of non-hydrogen atoms | 3212 |
| macromolecules | 2776 |
| ligands | 98 |
| solvent | 338 |
| Protein residues | 340 |
| RMS(bonds) | 0.008 |
| RMS(angles) | 1.10 |
| Ramachandran favored (%) | 98.21 |
| Ramachandran allowed (%) | 1.49 |
| Ramachandran outliers (%) | 0.30 |
| Rotamer outliers (%) | 0.00 |
| Clashscore | 8.06 |
| Average B-factor | 23.74 |
| macromolecules | 22.71 |
| ligands | 21.30 |
| solvent | 33.79 |

Statistics for the highest-resolution shell are shown in parentheses.

**Supplementary Table 2**. List of antibodies.

| Target | Species | Supplier | Identifier | Dilution |
|---|---|---|---|---|
| **p-AKT (S473)** | Rabbit | Cell Signaling Technology | 4060 | 1:1,000 |
| **AKT** | Rabbit | Cell Signaling Technology | 2920 | 1:1,000 |
| **p-ERK (T202/Y204)** | Rabbit | Cell Signaling Technology | 9101 | 1:1,000 |
| **ERK** | Rabbit | Cell Signaling Technology | 4695 | 1:1,000 |
| **Ras(G12D)** | Rabbit | Invitrogen (Thermo) | MA5-36256 | 1:1,000 |
| **Pan-Ras** | Rabbit | Abcam | 108602 | 1:5,000 |
| **GAPDH** | Mouse | Proteintech | 60004-1-Ig | 1:50,000 |
| **Ubiquitin** | Mouse | Santa Cruz Biotechnology | sc-8017 | 1:500 |
| **β-Actin** | Rabbit | Cell Signaling Technology | 8457 | 1:1,000 |

**Supplementary Table 3**. List of buffer compositions.

| Name | Composition |
|---|---|
| **RIPA Buffer** | 25 mM Tris 7.4<br>150 mM NaCl<br>0.1% SDS<br>1% NP-40<br>0.5% Sodium Deoxycholate |
| **5X SDS Loading Buffer** | 250 mM Tris 6.8<br>500 mM DTT<br>10% w/v SDS<br>0.1% w/v Bromophenol Blue<br>50% Glycerol |
| **1X TOWBIN Transfer Buffer** | 250 mM Tris<br>192 mM Glycine<br>pH 8.3 |
| **Lysis Buffer** | 20 mM Tris 8.0<br>500 mM NaCl<br>5 mM Imidazole |
| **Elution Buffer** | 20 mM Tris 8.0<br>300 mM NaCl<br>300 mM Imidazole |
| **TEV Cleavage Buffer** | 20 mM Tris 8.0<br>300 mM NaCl<br>1 mM EDTA |
| **SEC Buffer** | 20 mM HEPES 7.5<br>150 mM NaCl<br>1 mM $MgCl_2$ |
| **Nucleotide Exchange Buffer** | 20 mM HEPES 7.5<br>150 mM NaCl<br>1 mM $MgCl_2$<br>1 mM DTT |

**Supplementary Table 4**. Labeling selectivity of β-lactone K-Ras(G12D) inhibitors.

| Compound | Percentage Modification | | | | | | |
|---|---|---|---|---|---|---|---|
| | **G12D• GDP** | **G12D• GppNHp** | **WT• GDP** | **G12E• GDP** | **G13D• GDP** | **G12S• GDP** | **G12C• GDP** |
| **(*RS*)-1**[1] | 100 | 34 | 0 | 6.3 | 6 | ND | 100 |
| **(*R*)-2**[2] | 100 | ND | ND | ND | ND | ND | ND |
| **(2*R*,3*S*)-3**[2] | 100 | ND | ND | ND | ND | ND | ND |
| **(2*R*,3*S*)-4**[1] | 100 | ND | 2 | 0 | 3 | 5 | ND |
| **(2*R*,3*S*)-5**[1] | 94 | 90 | 0.5 | 0 | 0 | 0 | 96 |
| **(2*R*,3*S*)-6**[1] | 100 | 100 | 0 | 0 | 0 | 0 | 100 |
| **(*R*)-7**[1] | 100 | 100 | 0 | 0 | 0 | 0 | 100 |

[1] Conditions: 200 nM protein, 10 µM drug, 12 h room-temp incubation. [2] Conditions: 1 µM protein, 100 µM drug, 12 h room-temp incubation. ND, Not determined.

**Supplementary Table 5**. Labeling kinetics of β-lactone K-Ras(G12D) inhibitors.

| Compounds | $k_{obs}$ (min$^{-1}$) | |
|---|---|---|
| | **G12D•GDP** | **G12D•GppNHp** |
| **(*RS*)-1** | 0.372 | ND |
| **(*R*)-2** | 0.7434 | ND |
| **(2*R*,3*S*)-3** | 0.2796 | ND |
| **(2*R*,3*S*)-4** | 0.065 | 0.0031 |
| **(2*R*,3*S*)-5** | 0.143 | 0.0137 |
| **(2*R*,3*S*)-6** | 0.0704 | 0.658 |
| **(*R*)-7** | 0.133 | 0.0148 |

ND, Not determined.

**Supplementary Table 6**. Covalent β-lactone K-Ras(G12D) inhibitors induced protein stabilization.

| **Compound** | **ΔTm (ºC, relative to DMSO control)** | |
|---|---|---|
| | **G12D•GDP** | **G12D•GppNHp** |
| **(*RS*)-1** | 10.6 | 3.6 |
| **(*R*)-2** | 7.6 | -0.7 |
| **(2*R*,3*S*)-3** | ND | ND |
| **(2*R*,3*S*)-4** | 5 | -0.8 |
| **(2*R*,3*S*)-5** | -2.1 | -2.3 |
| **(2*R*,3*S*)-6** | 13.6 | 10.9 |
| **(*R*)-7** | 18.6 | 11.7 |
| **(2R,3S)-5a** | 16.0 | 17.8 |
| **(2R,3S)-5b** | 10.1 | 8.7 |
| **(2R,3S)-5c** | 14.8 | 16.3 |
| **(2R,3S)-5d** | 10.2 | 8.9 |
| **(2R,3S)-5e** | 15.1 | 16.3 |
| **(2R,3S)-5f** | 9.1 | 7.6 |

ND, Not determined.

**Supplementary Table 7**. PBS stability of β-lactone K-Ras(G12D) inhibitors.

| Compound # | half-life in PBS (h) |
|---|---|
| **(*RS*)-1** | ND |
| **(*R*)-2** | 1.718 |
| **(2*R*,3*S*)-3** | 2.547 |
| **(2*R*,3*S*)-4** | >24 |
| **(2*R*,3*S*)-5** | >24 |
| **(2*R*,3*S*)-6** | >24 |
| **(*R*)-7** | >24 |

ND, Not determined.

**Supplementary Table 8**. Growth inhibition of Ba/F3 KRAS G12D cells.

| Compound | IC50 without IL-3 (nM) | IC50 with IL-3 (nM) |
|---|---|---|
| **(*RS*)-1** | >10,000 | >10,000 |
| **(*R*)-2** | >10,000 | >10,000 |
| **(2*R*,3*S*)-3** | 6,671 | >10,000 |
| **(2*R*,3*S*)-4** | 330 | 428 |
| **(2*R*,3*S*)-5** | 356 | 976 |
| **(2*R*,3*S*)-6** | 161 | 2,738 |
| **(*R*)-7** | 72 | >10,000 |

**Supplementary Table 9**. Growth inhibition of cancer cell lines.

| **Compound** | **$GI_{50}$ (nM)** | | | | | | |
|---|---|---|---|---|---|---|---|
| | **SW1990** | **AsPC-1** | **AGS** | **H1299** | **HCT116** | **A549** | **A375** |
| **(*RS*)-1** | ND | ND | ND | ND | ND | ND | ND |
| **(*R*)-2** | ND | ND | ND | ND | ND | ND | ND |
| **(2*R*,3*S*)-3** | ND | ND | ND | ND | ND | ND | ND |
| **(2*R*,3*S*)-4** | 124 | 484 | 41 | 280 | 118 | 834 | 192 |
| **(2*R*,3*S*)-5** | 350.6 | 975.8 | 245.7 | 336.9 | 145.8 | 921.6 | 3,588 |
| **(2*R*,3*S*)-6** | 928.4 | 546 | 316.1 | 1,771 | 345.2 | 2,135 | 722.8 |
| **(*R*)-7** | 393.7 | 463.2 | 108.4 | >10,000 | >10,000 | >10,000 | >10,000 |

ND, Not determined.

**Supplementary Table 10**. Kinase profiling of Compound (*R*)-7 at 1 μM.

| Kinase | % Inhibition |
|---|---|
| AAK1 | 7 |
| ABL1 | 4 |
| ABL1 E255K | -10 |
| ABL1 F317I | 3 |
| ABL1 F317L | 5 |
| ABL1 G250E | 2 |
| ABL1 H396P | 0 |
| ABL1 M351T | 4 |
| ABL1 Q252H | -7 |
| ABL1 T315I | 1 |
| ABL1 Y253F | 4 |
| ABL2 (Arg) | 2 |
| ACVR1 (ALK2) | 1 |
| ACVR1 (ALK2) R206H | 3 |
| ACVR1B (ALK4) | 4 |
| ACVR2A | 3 |
| ACVR2B | 0 |
| ACVRL1 (ALK1) | -2 |
| ADRBK1 (GRK2) | 5 |
| ADRBK2 (GRK3) | 3 |
| AKT1 (PKB alpha) | 3 |
| AKT2 (PKB beta) | 2 |
| AKT3 (PKB gamma) | 10 |
| ALK | 3 |
| ALK C1156Y | 3 |
| ALK F1174L | 2 |
| ALK L1196M | 4 |
| ALK R1275Q | 2 |
| ALK T1151_L1152insT | 0 |
| AMPK (A1/B1/G2) | 0 |
| AMPK (A1/B1/G3) | -2 |
| AMPK (A1/B2/G1) | 2 |
| AMPK (A1/B2/G2) | 2 |
| AMPK (A1/B2/G3) | 3 |
| AMPK (A2/B1/G2) | 2 |
| AMPK (A2/B1/G3) | 4 |

| | |
|---|---|
| AMPK (A2/B2/G1) | 1 |
| AMPK (A2/B2/G2) | -2 |
| AMPK (A2/B2/G3) | 0 |
| AMPK A1/B1/G1 | -2 |
| AMPK A2/B1/G1 | 6 |
| ANKK1 | 17 |
| AURKA (Aurora A) | 4 |
| AURKB (Aurora B) | 4 |
| AURKC (Aurora C) | 11 |
| AXL | -2 |
| AXL R499C | 0 |
| BLK | 16 |
| BMPR1A (ALK3) | 0 |
| BMPR1B (ALK6) | 6 |
| BMPR2 | 17 |
| BMX | 9 |
| BRAF | -6 |
| BRAF | 4 |
| BRAF V599E | -4 |
| BRAF V599E | 3 |
| BRSK1 (SAD1) | 4 |
| BRSK2 | 0 |
| BTK | 9 |
| CAMK1 (CaMK1) | 4 |
| CAMK1D (CaMKI delta) | 1 |
| CAMK1G (CAMKI gamma) | 4 |
| CAMK2A (CaMKII alpha) | 1 |
| CAMK2B (CaMKII beta) | 14 |
| CAMK2D (CaMKII delta) | 5 |
| CAMK2G (CaMKII gamma) | -5 |
| CAMK4 (CaMKIV) | 14 |
| CAMKK1 (CAMKKA) | 2 |
| CAMKK2 (CaMKK beta) | 1 |
| CASK | 2 |
| CDC42 BPA (MRCKA) | 0 |
| CDC42 BPB (MRCKB) | 3 |
| CDC42 BPG (MRCKG) | -1 |
| CDC7/DBF4 | 3 |
| CDK1/cyclin B | 1 |

| | |
|---|---|
| CDK11 (Inactive) | 17 |
| CDK11/cyclin C | 11 |
| CDK13/cyclin K | -7 |
| CDK14 (PFTK1)/cyclin Y | 0 |
| CDK16 (PCTK1)/cyclin Y | 7 |
| CDK17/cyclin Y | 4 |
| CDK18/cyclin Y | 3 |
| CDK2/cyclin A | 7 |
| CDK2/cyclin A1 | 6 |
| CDK2/cyclin E1 | -1 |
| CDK2/cyclin O | -2 |
| CDK3/cyclin E1 | 0 |
| CDK4/cyclin D1 | -10 |
| CDK4/cyclin D3 | -2 |
| CDK5 (Inactive) | 0 |
| CDK5/p25 | 4 |
| CDK5/p35 | 6 |
| CDK6/cyclin D1 | -5 |
| CDK7/cyclin H/MNAT1 | -4 |
| CDK8/cyclin C | 7 |
| CDK9 (Inactive) | 7 |
| CDK9/cyclin K | -1 |
| CDK9/cyclin T1 | 7 |
| CDKL5 | 3 |
| CHEK1 (CHK1) | 9 |
| CHEK2 (CHK2) | 0 |
| CHUK (IKK alpha) | 3 |
| CLK1 | 1 |
| CLK2 | 8 |
| CLK3 | 2 |
| CLK4 | 4 |
| CSF1R (FMS) | 16 |
| CSK | 8 |
| CSNK1A1 (CK1 alpha 1) | -7 |
| CSNK1A1L | 2 |
| CSNK1D (CK1 delta) | 8 |
| CSNK1E (CK1 epsilon) | 6 |
| CSNK1E (CK1 epsilon) R178C | 1 |
| CSNK1G1 (CK1 gamma 1) | 1 |

| | |
|---|---|
| CSNK1G2 (CK1 gamma 2) | 1 |
| CSNK1G3 (CK1 gamma 3) | 0 |
| CSNK2A1 (CK2 alpha 1) | 4 |
| CSNK2A2 (CK2 alpha 2) | 5 |
| DAPK1 | -4 |
| DAPK2 | 2 |
| DAPK3 (ZIPK) | 3 |
| DCAMKL1 (DCLK1) | 4 |
| DCAMKL2 (DCK2) | 7 |
| DDR1 | 0 |
| DDR2 | 1 |
| DDR2 N456S | 5 |
| DDR2 T654M | 5 |
| DMPK | -7 |
| DNA-PK | 6 |
| DYRK1A | 2 |
| DYRK1B | 0 |
| DYRK2 | 1 |
| DYRK3 | 1 |
| DYRK4 | 1 |
| EEF2K | 5 |
| EGFR (ErbB1) | -6 |
| EGFR (ErbB1) C797S | -2 |
| EGFR (ErbB1) d746-750 | -5 |
| EGFR (ErbB1) d747-749 A750P | 1 |
| EGFR (ErbB1) G719C | -2 |
| EGFR (ErbB1) G719S | 3 |
| EGFR (ErbB1) L858R | -2 |
| EGFR (ErbB1) L861Q | -1 |
| EGFR (ErbB1) T790M | 4 |
| EGFR (ErbB1) T790M C797S L858R | -2 |
| EGFR (ErbB1) T790M L858R | -1 |
| EIF2AK2 (PKR) | 3 |
| EPHA1 | 4 |
| EPHA2 | 0 |
| EPHA3 | 1 |
| EPHA4 | 9 |
| EPHA5 | 4 |
| EPHA6 | -9 |

| | |
|---|---|
| EPHA7 | -1 |
| EPHA8 | 7 |
| EPHB1 | 3 |
| EPHB2 | 8 |
| EPHB3 | 5 |
| EPHB4 | 6 |
| ERBB2 (HER2) | 7 |
| ERBB4 (HER4) | 4 |
| ERN1 | -1 |
| ERN2 | 1 |
| FER | 4 |
| FES (FPS) | 2 |
| FGFR1 | 5 |
| FGFR1 V561M | 1 |
| FGFR2 | 10 |
| FGFR2 N549H | 3 |
| FGFR3 | -1 |
| FGFR3 G697C | 4 |
| FGFR3 K650E | 0 |
| FGFR3 K650M | 3 |
| FGFR3 V555M | -3 |
| FGFR4 | -4 |
| FGR | 19 |
| FLT1 (VEGFR1) | 1 |
| FLT3 | 9 |
| FLT3 D835Y | 6 |
| FLT3 ITD | 2 |
| FLT4 (VEGFR3) | 4 |
| FRAP1 (mTOR) | 1 |
| FRK (PTK5) | 9 |
| FYN | -4 |
| FYN A | 2 |
| GAK | -5 |
| GRK1 | 1 |
| GRK4 | 8 |
| GRK5 | 2 |
| GRK6 | -2 |
| GRK7 | 6 |
| GSG2 (Haspin) | 2 |

| | |
|---|---|
| GSK3A (GSK3 alpha) | 4 |
| GSK3B (GSK3 beta) | 2 |
| HCK | 7 |
| HIPK1 (Myak) | 2 |
| HIPK2 | 3 |
| HIPK3 (YAK1) | 3 |
| HIPK4 | 7 |
| HUNK | 2 |
| ICK | 1 |
| IGF1R | 13 |
| IKBKB (IKK beta) | 8 |
| IKBKE (IKK epsilon) | 11 |
| INSR | 4 |
| INSRR (IRR) | 2 |
| IRAK1 | 1 |
| IRAK3 | 5 |
| IRAK4 | 7 |
| ITK | 9 |
| JAK1 | 0 |
| JAK2 | 5 |
| JAK2 JH1 JH2 | 2 |
| JAK2 JH1 JH2 V617F | 4 |
| JAK3 | 3 |
| KDR (VEGFR2) | 4 |
| KIT | 9 |
| KIT A829P | -2 |
| KIT D816H | 6 |
| KIT D816V | -5 |
| KIT D820E | 0 |
| KIT N822K | 4 |
| KIT T670E | 3 |
| KIT T670I | 3 |
| KIT V559D | 3 |
| KIT V559D T670I | -2 |
| KIT V559D V654A | -1 |
| KIT V560G | -1 |
| KIT V654A | 1 |
| KIT Y823D | 2 |
| KSR2 | 4 |

| | |
|---|---|
| LATS2 | 13 |
| LCK | 13 |
| LIMK1 | 4 |
| LIMK2 | 2 |
| LRRK2 | 6 |
| LRRK2 FL | -3 |
| LRRK2 G2019S | 0 |
| LRRK2 G2019S FL | -9 |
| LRRK2 I2020T | 1 |
| LRRK2 R1441C | -1 |
| LTK (TYK1) | 7 |
| LYN A | 1 |
| LYN B | 4 |
| MAP2K1 (MEK1) | -2 |
| MAP2K1 (MEK1) | 4 |
| MAP2K1 (MEK1) S218D S222D | 1 |
| MAP2K2 (MEK2) | -1 |
| MAP2K2 (MEK2) | 6 |
| MAP2K4 (MEK4) | 0 |
| MAP2K5 (MEK5) | -3 |
| MAP2K6 (MKK6) | 0 |
| MAP2K6 (MKK6) | -4 |
| MAP2K6 (MKK6) S207E T211E | -2 |
| MAP3K10 (MLK2) | -1 |
| MAP3K11 (MLK3) | 3 |
| MAP3K14 (NIK) | -4 |
| MAP3K19 (YSK4) | 0 |
| MAP3K2 (MEKK2) | 1 |
| MAP3K3 (MEKK3) | 0 |
| MAP3K5 (ASK1) | 4 |
| MAP3K7/MAP3K7IP1 (TAK1-TAB1) | 4 |
| MAP3K8 (COT) | 2 |
| MAP3K9 (MLK1) | 6 |
| MAP4K1 (HPK1) | 0 |
| MAP4K2 (GCK) | 4 |
| MAP4K3 (GLK) | 5 |
| MAP4K4 (HGK) | 17 |
| MAP4K5 (KHS1) | 8 |
| MAPK1 (ERK2) | 3 |

| | |
|---|---|
| MAPK10 (JNK3) | -4 |
| MAPK10 (JNK3) | 5 |
| MAPK11 (p38 beta) | 7 |
| MAPK12 (p38 gamma) | 7 |
| MAPK13 (p38 delta) | 4 |
| MAPK14 (p38 alpha) | 2 |
| MAPK14 (p38 alpha) Direct | 1 |
| MAPK15 (ERK7) | -14 |
| MAPK3 (ERK1) | 2 |
| MAPK7 (ERK5) | 0 |
| MAPK8 (JNK1) | -9 |
| MAPK8 (JNK1) | -7 |
| MAPK9 (JNK2) | 4 |
| MAPK9 (JNK2) | 0 |
| MAPKAPK2 | 1 |
| MAPKAPK3 | -1 |
| MAPKAPK5 (PRAK) | 10 |
| MARK1 (MARK) | 2 |
| MARK2 | 5 |
| MARK3 | 10 |
| MARK4 | 5 |
| MASTL | -4 |
| MATK (HYL) | 8 |
| MELK | 16 |
| MERTK (cMER) | 7 |
| MERTK (cMER) A708S | -9 |
| MET (cMet) | 16 |
| MET (cMet) Y1235D | -1 |
| MET D1228H | 2 |
| MET M1250T | 0 |
| MINK1 | 11 |
| MKNK1 (MNK1) | 3 |
| MKNK2 (MNK2) | 0 |
| MLCK (MLCK2) | -11 |
| MLK4 | -2 |
| MST1R (RON) | 8 |
| MST4 | 0 |
| MUSK | 7 |
| MYLK (MLCK) | -7 |

| | |
|---|---|
| MYLK2 (skMLCK) | 4 |
| MYLK4 | 4 |
| MYO3A (MYO3 alpha) | 4 |
| MYO3B (MYO3 beta) | 3 |
| NEK1 | 3 |
| NEK2 | 8 |
| NEK4 | 9 |
| NEK6 | 6 |
| NEK8 | -3 |
| NEK9 | -1 |
| NIM1K | 1 |
| NLK | 7 |
| NTRK1 (TRKA) | 8 |
| NTRK2 (TRKB) | 9 |
| NTRK3 (TRKC) | 11 |
| NUAK1 (ARK5) | 3 |
| NUAK2 | 14 |
| PAK1 | 9 |
| PAK2 (PAK65) | 2 |
| PAK3 | 7 |
| PAK4 | 10 |
| PAK6 | 10 |
| PAK7 (KIAA1264) | 3 |
| PASK | 12 |
| PDGFRA (PDGFR alpha) | 0 |
| PDGFRA D842V | 0 |
| PDGFRA T674I | 0 |
| PDGFRA V561D | -2 |
| PDGFRB (PDGFR beta) | 4 |
| PDK1 | 15 |
| PDK1 Direct | 2 |
| PHKG1 | 7 |
| PHKG2 | 15 |
| PI4K2A (PI4K2 alpha) | -6 |
| PI4K2B (PI4K2 beta) | -5 |
| PI4KA (PI4K alpha) | 2 |
| PI4KB (PI4K beta) | -2 |
| PIK3C2A (PI3K-C2 alpha) | 1 |
| PIK3C2B (PI3K-C2 beta) | 2 |

| | |
|---|---|
| PIK3C2G (PI3K-C2 gamma) | 0 |
| PIK3C3 (hVPS34) | 2 |
| PIK3CA E542K/PIK3R1 (p110 alpha E542K/p85 alpha) | 0 |
| PIK3CA E545K/PIK3R1 (p110 alpha E545K/p85 alpha) | -9 |
| PIK3CA/PIK3R1 (p110 alpha/p85 alpha) | -7 |
| PIK3CA/PIK3R3 (p110 alpha/p55 gamma) | -3 |
| PIK3CB/PIK3R1 (p110 beta/p85 alpha) | 0 |
| PIK3CB/PIK3R2 (p110 beta/p85 beta) | -17 |
| PIK3CD/PIK3R1 (p110 delta/p85 alpha) | -4 |
| PIK3CG (p110 gamma) | -6 |
| PIM1 | 1 |
| PIM2 | 5 |
| PIM3 | 4 |
| PIP4K2A | -3 |
| PIP5K1A | -7 |
| PIP5K1B | -11 |
| PIP5K1C | 3 |
| PKMYT1 | -3 |
| PKN1 (PRK1) | 5 |
| PKN2 (PRK2) | 3 |
| PLK1 | 6 |
| PLK2 | 3 |
| PLK3 | 12 |
| PLK4 | -1 |
| PRKACA (PKA) | 1 |
| PRKACB (PRKAC beta) | -2 |
| PRKACG (PRKAC gamma) | 16 |
| PRKCA (PKC alpha) | 14 |
| PRKCB1 (PKC beta I) | 11 |
| PRKCB2 (PKC beta II) | 5 |
| PRKCD (PKC delta) | 18 |
| PRKCE (PKC epsilon) | 7 |
| PRKCG (PKC gamma) | 5 |
| PRKCH (PKC eta) | 15 |
| PRKCI (PKC iota) | 6 |
| PRKCN (PKD3) | 7 |
| PRKCZ (PKC zeta) | 10 |

| | |
|---|---|
| PRKD1 (PKC mu) | 10 |
| PRKD2 (PKD2) | 8 |
| PRKG1 | 0 |
| PRKG2 (PKG2) | 5 |
| PRKX | 10 |
| PTK2 (FAK) | 7 |
| PTK2B (FAK2) | 4 |
| PTK6 (Brk) | 5 |
| RAF1 (cRAF) Y340D Y341D | -1 |
| RAF1 (cRAF) Y340D Y341D | -1 |
| RET | 4 |
| RET A883F | -3 |
| RET G691S | 1 |
| RET M918T | 2 |
| RET S891A | -6 |
| RET V804E | -3 |
| RET V804L | 3 |
| RET V804M | 2 |
| RET Y791F | 3 |
| RIPK2 | 2 |
| RIPK3 | 0 |
| ROCK1 | 3 |
| ROCK2 | 3 |
| ROS1 | 5 |
| RPS6KA1 (RSK1) | 0 |
| RPS6KA2 (RSK3) | 5 |
| RPS6KA3 (RSK2) | 5 |
| RPS6KA4 (MSK2) | 10 |
| RPS6KA5 (MSK1) | 2 |
| RPS6KA6 (RSK4) | 1 |
| RPS6KB1 (p70S6K) | 21 |
| RPS6KB2 (p70S6Kb) | 8 |
| SBK1 | 0 |
| SGK (SGK1) | 0 |
| SGK2 | 2 |
| SGKL (SGK3) | 6 |
| SIK1 | -2 |
| SIK3 | 7 |
| SLK | -2 |

| | |
|---|---|
| SNF1LK2 | 5 |
| SPHK1 | 2 |
| SPHK2 | -4 |
| SRC | 7 |
| SRC N1 | 4 |
| SRMS (Srm) | 8 |
| SRPK1 | 3 |
| SRPK2 | 1 |
| STK16 (PKL12) | 3 |
| STK17A (DRAK1) | -13 |
| STK17B (DRAK2) | 1 |
| STK22B (TSSK2) | 5 |
| STK22D (TSSK1) | 9 |
| STK23 (MSSK1) | 1 |
| STK24 (MST3) | 1 |
| STK25 (YSK1) | 3 |
| STK3 (MST2) | 0 |
| STK32B (YANK2) | 1 |
| STK32C (YANK3) | -1 |
| STK33 | 5 |
| STK38 (NDR) | 1 |
| STK38L (NDR2) | 1 |
| STK39 (STLK3) | 6 |
| STK4 (MST1) | 6 |
| SYK | 5 |
| TAOK1 | 1 |
| TAOK2 (TAO1) | 6 |
| TAOK3 (JIK) | 0 |
| TBK1 | 3 |
| TEC | 3 |
| TEK (Tie2) | 5 |
| TEK (TIE2) R849W | -5 |
| TEK (TIE2) Y1108F | 1 |
| TEK (TIE2) Y897S | 2 |
| TESK1 | 0 |
| TESK2 | 1 |
| TGFBR1 (ALK5) | 1 |
| TGFBR2 | 2 |
| TLK1 | -1 |

| | |
|---|---|
| TLK2 | -3 |
| TNIK | -3 |
| TNK1 | 1 |
| TNK2 (ACK) | -11 |
| TTK | -15 |
| TXK | 9 |
| TYK2 | 5 |
| TYRO3 (RSE) | 3 |
| ULK1 | -10 |
| ULK2 | 1 |
| ULK3 | -1 |
| VRK2 | 6 |
| WEE1 | 4 |
| WNK1 | 5 |
| WNK2 | -3 |
| WNK3 | 5 |
| YES1 | 0 |
| ZAK | 0 |
| ZAP70 | 10 |

**Chemical Synthesis**

General Experiment Procedure

All reactions were performed in oven-dried glassware fitted with rubber septa under a positive pressure of argon, unless otherwise noted. Air- and moisture-sensitive liquids were transferred via syringe. Solutions were concentrated by rotary evaporation at or below 40 °C. Analytical thin-layer chromatography (TLC) was performed using glass plates pre-coated with silica gel (0.25-mm, 60-Å pore size, 230−400 mesh, Merck KGA) impregnated with a fluorescent indicator (254 nm). TLC plates were visualized by exposure to ultraviolet light (UV), then were stained by submersion in a 10% solution of phosphomolybdic acid (PMA) in ethanol or an 2% aqueous solution of potassium permanganate followed by brief heating on a hot plate. Flash column chromatography was performed with Teledyne ISCO CombiFlash EZ Prep chromatography system, employing pre-packed silica gel cartridges (Teledyne ISCO RediSep).

Solvents and Reagents

Anhydrous solvents were purchased from Acros Organics. Unless specified below, all chemical reagents were purchased from Sigma-Aldrich, AK Scientific, AstaTech, Combi-Blocks Chemscene, and Advanced ChemBlock.

Instrumentation

Proton nuclear magnetic resonance ($^1$H NMR) spectra, carbon nuclear magnetic resonance ($^{13}$C NMR) spectra, and fluorine nuclear magnetic resonance ($^{19}$F NMR) spectra were recorded on Bruker AvanceIII HD instrument (400 MHz/100 MHz/376 MHz) or Varian Inova 600 instrument (600 MHz/126 MHz/564 MHz) at 23 °C. Proton chemical shifts are expressed in parts per million (ppm, δ scale) and are referenced to residual protium in the NMR solvent ($CHCl_3$: δ 7.26, $D_2HCOD$: δ 3.31, $CD_2HCOCD_3$: δ 2.05, $CD_2HSOCD_3$: δ 2.50). Carbon chemical shifts are expressed in parts per million (ppm, δ scale) and are referenced to the carbon resonance of the NMR solvent ($CDCl_3$: δ 77.0, $CD_3OD$: δ 49.0, $CD_3\underline{C}OCD_3$: δ 206.26, $CD_3SOCD_3$: δ 39.52). Fluorine chemical shifts are expressed in parts per million (ppm, δ scale) and are referenced to an internal standard of trifluoroacetic acid (–76.55 ppm). Data are represented as follows: chemical shift, multiplicity (s = singlet, d = doublet, t = triplet, q = quartet, dd = doublet of doublets, dt = doublet of triplets, m = multiplet, br = broad, app = apparent), integration, and coupling constant (*J*) in Hertz (Hz). Accurate (high-resolution) mass spectra were obtained using a Waters Xevo G2-XS time-of-flight mass spectrometer.

Mini-workup

When a mini-workup (A/B) is indicated in the procedure, it was performed as follows: an aliquot (5 µL) of the reaction mixture was retrieved with a glass pipet and added to a plastic vial containing 0.2 mL organic solvent A and 0.2 mL aqueous solution B. The vial was shaken vigorously and allowed to stand until the two layers partitioned. The organic layer was then used for TLC or LC-MS analysis as specified in the procedure.

Monitoring Reaction Progress by LC-MS

When LC-MS analysis of the reaction mixture is indicated in the procedure, it was performed as follows. An aliquot (1 µL) of the reaction mixture (or the organic phase of a mini-workup mixture) was diluted with 100 µL 1:1 acetonitrile:water. 1 µL of the diluted solution was injected onto a Waters Acquity UPLC BEH C18 1.7 µm column and eluted with a linear gradient of 5–95% acetonitrile/water (+0.1% formic acid) over 3.0 min. Chromatograms were recorded with a UV detector set at 254 nm and a time-of-flight mass spectrometer (Waters Xevo G2-XS).

Synthesis of (*RS*)-1

Note: The synthesis of the core structure was adapted from known methods[58,59] with modifications.

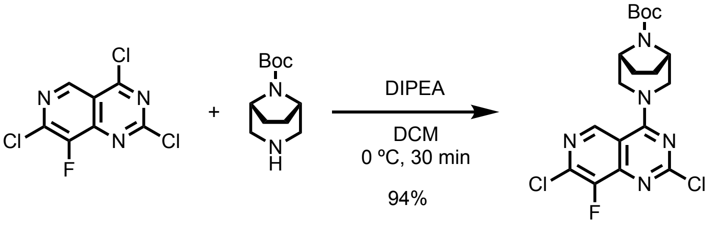


**S1** **S2** **S3**

***tert*-Butyl 3-(2,7-dichloro-8-fluoropyrido[4,3-d]pyrimidin-4-yl)-3,8-diazabicyclo[3.2.1]octane-8-carboxylate (S3).** A 100-mL round-bottom flask equipped with a stir bar was charged with 2,4,7-trichloro-8-fluoro-pyrido[4,3-d]pyrimidine (1262.3 mg, 5.0000 mmol), *tert*-butyl 3,8-diazabicyclo[3.2.1]octane-8-carboxylate (1061.45 mg, 5.0000 mmol, 1.0 equiv). DCM (25 mL) and *N*,*N*-diisopropylethylamine (1.74 mL, 10 mmol) were added via syringe. The mixture was stirred at 0 ºC for 30 min, at which point the full consumption of the starting material was judged by LC-MS. The reaction mixture was concentrated. The residue was purified by flash column chromatography (0–100% EA-hexanes, 40-g RediSep(R) Rf column, Teledyne ISCO, Lincoln, NE). The desired product was obtained as a white crystalline tert-butyl 3-(2,7-dichloro-8-fluoro-pyrido[4,3-d]pyrimidin-4-yl)-3,8-diazabicyclo[3.2.1]octane-8-carboxylate (2019.0 mg, 4.7151 mmol, 94% yield). TLC $R_f$ = 0.56 (50% EA-Hexanes, $UV_{254}$). $^1$H NMR (400 MHz, $CDCl_3$) δ 8.84 (d, *J* = 0.6 Hz, 1H), 4.70 – 4.22 (m, 4H), 3.72 (s, 2H), 1.98 (dd, *J* = 8.7, 4.3 Hz, 2H), 1.67 (d, *J* = 7.7 Hz, 2H), 1.51 (s, 9H). $^{13}$C NMR (100 MHz, $CDCl_3$) δ 163.72, 163.70, 161.33, 153.36, 149.80, 148.95, 148.83, 147.13, 144.20, 144.12, 139.11, 138.94, 111.15, 80.81, 77.48, 76.84, 54.34, 28.51. $^{19}$F NMR (376 MHz, $CDCl_3$) δ -131.93. Accurate MS (ESI-TOF) calculated for $C_{18}H_{21}Cl_2FN_5O_2$ $[M + H]^+$ 428.1056, found 428.1121.

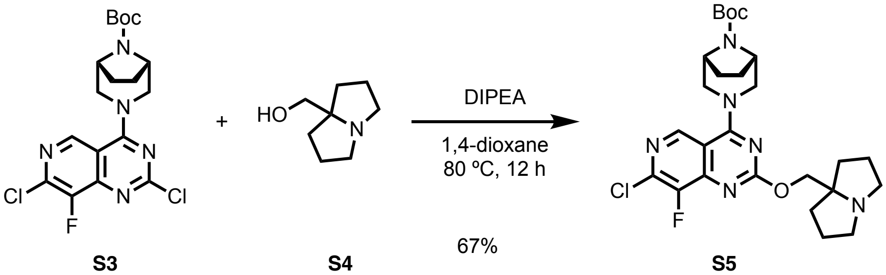


**S3** **S4** **S5**

***tert*-Butyl 3-(7-chloro-8-fluoro-2-((tetrahydro-1H-pyrrolizin-7a(5H)-yl)methoxy)pyrido[4,3-d]pyrimidin-4-yl)-3,8-diazabicyclo[3.2.1]octane-8-carboxylate (S5).** A 100-mL round-bottom flask equipped with a stir bar was charged with *tert*-butyl 3-(2,7-dichloro-8-fluoro-pyrido[4,3-d]pyrimidin-4-yl)-3,8-diazabicyclo[3.2.1]octane-8-carboxylate (2019.0 mg, 4.7151 mmol). Sealed with a rubber septum, the flask was repeatedly deaerated and backfilled with argon three times. 1,2,3,5,6,7-Hexahydropyrrolizin-8-ylmethanol (1997.0 mg, 14.141 mmol, 3.0 equiv), 1,4-dioxane (20 mL) and *N*,*N*-diisopropylethylamine (2.46 mL, 14.1 mmol, 3.0 equiv) were added sequentially. The mixture was stirred at 80 ºC for 12 h, at which point the full consumption of the starting material was judged by LC-MS. The reaction mixture was concentrated *in vacuo*. The residue was partitioned between EA (100 mL) and water (100 mL). The aqueous phase was further extracted with EA (50 mL x 3). The combined organic solution was sequentially washed with water, brine, dried over anhydrous sodium sulfate and concentrated. The residue was purified by flash column chromatography (0–30% MeOH*-DCM, *MeOH contains 2% ammonium hydroxide, 24-g RediSep(R) Rf column, Teledyne ISCO, Lincoln, NE). The desired product was obtained as a white crystalline (1695.0 mg, 3.1791 mmol, 67% yield). TLC $R_f$ = 0.51 (20% MeOH-DCM, $UV_{254}$). $^1$H NMR (400 MHz, $CDCl_3$) δ 8.67 (s, 1H), 4.45 (d, *J* = 12.6 Hz, 2H), 4.32 (s, 2H), 4.13 (s, 2H), 3.61 (s, 2H), 3.07 (dt, *J* = 10.7, 5.7 Hz, 2H), 2.60 (ddd, *J* = 10.2, 7.5, 6.2 Hz, 2H), 2.02 (ddd, *J* = 12.3, 6.6, 5.2 Hz, 2H), 1.90 (dd, *J* = 8.5, 4.2 Hz, 2H), 1.83 (ddd, *J* = 13.1, 6.6, 5.3 Hz, 4H), 1.70 – 1.56 (m, 3H), 1.47 (s, 9H). $^{13}$C NMR (100 MHz, $CDCl_3$) δ 165.38, 165.36, 164.54, 153.39, 150.60, 150.49, 149.87, 147.22, 143.93, 143.86, 138.02, 137.85, 110.58, 110.57, 80.53, 73.46, 72.49, 65.87, 55.69, 54.13, 36.04, 28.49, 25.63, 15.32. $^{19}$F NMR (376 MHz, $CDCl_3$) δ -134.29. Accurate MS (ESI-TOF) calculated for $C_{26}H_{35}ClFN_6O_3$ $[M + H]^+$ 533.2443, found 533.2467.

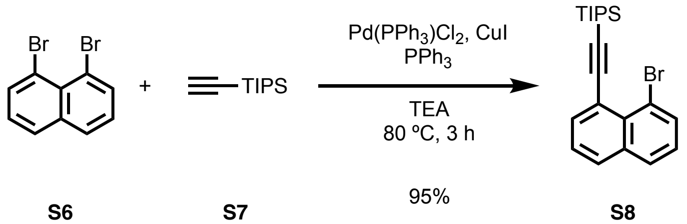


**S6** **S7** **S8**

**((8-Bromonaphthalen-1-yl)ethynyl)triisopropylsilane (S8).** A 100 mL round-bottom flask equipped with a stir bar was charged with 1,8-dibromonaphthalene (1429.8 mg, 5.0000 mmol), (triisopropylsilyl)acetylene (1094.3 mg, 6.0000 mmol, 1.2 equiv), bis(triphenylphosphine)palladium(II) dichloride (175.5 mg, 0.2500 mmol, 0.05 equiv), copper(I) iodide (95.2 mg, 0.500 mmol, 0.1 equiv), and triphenylphosphine (131.2 mg, 0.5000 mmol, 0.1 equiv). Sealed with a rubber septum, the flask was repeatedly deaerated and backfilled with argon for three times. Triethylamine (25 mL) was added via syringe. The mixture was stirred at 80 ºC for 150 min, at which point the full consumption of 1,8-dibromonaphthalene was judged by TLC. The reaction mixture was cooled to room temperature, and partitioned between EA (100 mL) and water (100 mL). The organic phase was sequentially washed with water (100 mL) and brine (100 mL), dried over anhydrous sodium sulfate, and concentrated. The residue was purified by flash column chromatography (100% Hexanes, 80-g RediSep(R) Rf column, Teledyne ISCO, Lincoln, NE). The desired product was obtained as a pale yellow solid (1841.0 mg, 4.7520 mmol, 95% yield). TLC $R_f$ = 0.74 (10% EA-Hexanes, $UV_{254}$). $^1$H NMR (400 MHz, $CDCl_3$) δ 7.89 (dd, *J* = 7.3, 1.4 Hz, 1H), 7.82 (dd, *J* = 7.4, 1.3 Hz, 1H), 7.78 (td, *J* = 8.2, 1.3 Hz, 2H), 7.40 (dd, *J* = 8.2, 7.2 Hz, 1H), 7.29 – 7.23 (m, 1H), 1.26 – 1.16 (m, 21H). $^{13}$C NMR (100 MHz, $CDCl_3$) δ 137.75, 135.71, 130.74, 130.05, 129.14, 126.50, 125.67, 121.50, 121.03, 107.15, 101.52, 18.96, 11.71.

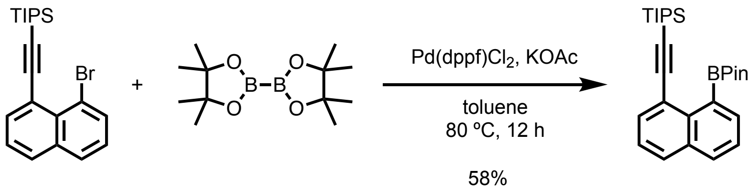


S8 B$_2$Pin$_2$ S9

**Triisopropyl((8-(4,4,5,5-tetramethyl-1,3,2-dioxaborolan-2-yl)naphthalen-1-yl)ethynyl)silane (S9).** A 100-mL round-bottom flask equipped with a stir bar was charged with ((8-Bromonaphthalen-1-yl)ethynyl)triisopropylsilane (1840.9 mg, 4.7520 mmol), Bis(pinacolato)diboron (2413.2 mg, 9.5030 mmol, 2.0 equiv), [1,1-bis(diphenylphosphino)ferrocene]dichloropalladium(II) (347.7 mg, 0.4750 mmol, 0.1 equiv), and Potassium acetate (1632.1 mg, 16.6308 mmol, 3.5 equiv). Sealed with a rubber septum, the flask was repeatedly deaerated and backfilled with argon three times. Toluene (20 mL) was added. the mixture was stirred at 80 ºC for 8 h, at which point the full consumption of the starting material was judged by TLC. The reaction mixture was cooled to room temperature, diluted with EA (100 mL), and filtered through a tightly packed Celite column. The filtrate was concentrated *in vacuo*. The residue was purified by flash column chromatography (0–20% EA-hexanes, 80-g RediSep(R) Rf column, Teledyne ISCO, Lincoln, NE). The desired product was obtained as a pale-yellow solid (1200.1 mg,2.7620 mmol, 58% yield). TLC $R_f$ = 0.43 (10% EA-hexanes, $UV_{254}$). $^1$H NMR (400 MHz, $CDCl_3$) δ 7.88 (d, $J$ = 1.3 Hz, 1H), 7.86 (t, $J$ = 1.3 Hz, 1H), 7.82 (dd, $J$ = 4.1, 1.3 Hz, 1H), 7.80 (dd, $J$ = 2.7, 1.3 Hz, 1H), 7.48 (dd, $J$ = 8.2, 6.8 Hz, 1H), 7.43 (dd, $J$ = 8.2, 7.2 Hz, 1H), 1.47 (s, 12H), 1.21 (d, $J$ = 2.2 Hz, 21H). $^{13}$C NMR (100 MHz, $CDCl_3$) δ 135.46, 134.78, 133.93, 133.62, 130.65, 129.61, 125.39, 125.31, 122.73, 110.09, 99.37, 84.28, 25.28, 19.07, 12.08.

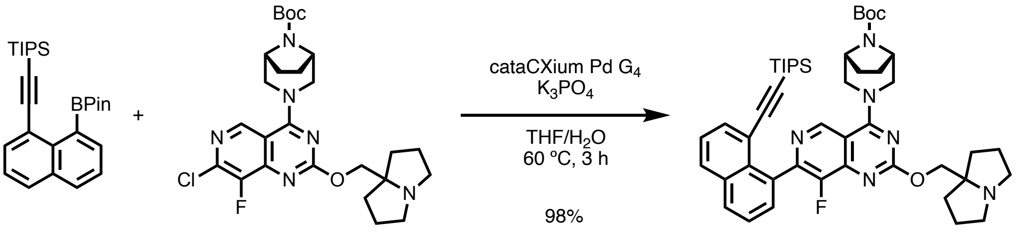


S9 S5 S10

***tert*-Butyl 3-(8-fluoro-2-((tetrahydro-1*H*-pyrrolizin-7a(5*H*)-yl)methoxy)-7-(8-((triisopropylsilyl)ethynyl)naphthalen-1-yl)pyrido[4,3-*d*]pyrimidin-4-yl)-3,8-diazabicyclo[3.2.1]octane-8-carboxylate (S10).**

A 100-mL round-bottom flask equipped with a stir bar was charged with *tert*-butyl 3-[7-chloro-8-fluoro-2-(1,2,3,5,6,7-hexahydropyrrolizin-8-ylmethoxy)pyrido[4,3-d]pyrimidin-4-yl]-3,8-diazabicyclo[3.2.1]octane-8-carboxylate (1599.1 mg, 3.0000 mmol), triisopropyl-[2-[8-(4,4,5,5-tetramethyl-1,3,2-dioxaborolan-2-yl)-1-naphthyl]ethynyl]silane (1694.5 mg, 3.9000 mmol, 1.3 equiv), and cataCXium Pd G4 pre-catalyst (222.7 mg, 0.3000, 0.1 equiv mmol). THF (25 mL) was added. The mixture was degassed by bubbling with a stream of Argon for 30 min. Potassium phosphate (1.5 M, 6.6 mL, 9.9 mmol, 3.3 equiv) was added. The mixture was stirred at 60 ºC for 3 h, at which point the consumption of the limiting reagent was judged by LC-MS. The reaction mixture was cooled to ambient temperature, and concentrated *in vacuo*. The residue was partitioned between EA (100 mL) and saturated sodium bicarbonate solution (100 mL). The aqueous phase was further extracted with EA (50 mL x 3). The combined organic solution was washed sequentially with water, brine, and dried over anhydrous sodium sulfate. Upon concentration, the residue was purified by flash column chromatography (0–30% MeOH*-DCM, *MeOH contains 2% ammonium hydroxide, 40-g RediSep(R) Rf column, Teledyne ISCO, Lincoln, NE). The desired product was obtained as a brown crystalline (2374.0 mg,2.9490 mmol, 98 % yield). $^1$H NMR (400 MHz, $CDCl_3$) δ 9.03 (s, 1H), 7.90 (ddd, *J* = 15.6, 7.8, 1.8 Hz, 2H), 7.77 (dd, *J* = 7.2, 1.3 Hz, 1H), 7.59 – 7.47 (m, 2H), 7.42 (dd, *J* = 8.2, 7.2 Hz, 1H), 4.82 (d, *J* = 12.9 Hz, 1H), 4.34 (s, 2H), 4.20 – 4.08 (m, 3H), 3.78 (s, 1H), 3.39 (s, 1H), 3.08 (dt, *J* = 10.6, 5.5 Hz, 2H), 2.61 (ddd, *J* = 10.1, 7.6, 6.2 Hz, 2H), 2.12 – 2.02 (m, 2H), 2.01 – 1.91 (m, 4H), 1.89 – 1.79 (m, 4H), 1.74 – 1.57 (m, 4H), 1.49 (s, 10H), 1.19 (s, 2H), 0.85 (dd, *J* = 7.4, 3.5 Hz, 18H), 0.50 (p, *J* = 7.5 Hz, 3H). $^{13}$C NMR (100 MHz, $CDCl_3$) δ 164.24, 152.17, 149.61, 149.56, 149.49, 147.00, 146.86, 143.53, 143.46, 136.72, 134.45, 132.97, 132.94, 130.60, 130.41, 130.20, 129.79, 125.48, 125.35, 120.37, 111.09, 111.07, 107.30, 98.02, 80.43, 74.98, 73.33, 72.41, 55.72, 36.12, 36.05, 28.51, 25.77, 25.75, 24.91, 18.66, 18.53, 11.55. $^{19}$F NMR (376 MHz, $CDCl_3$) δ -137.81. Accurate MS (ESI-TOF) calculated for $C_{47}H_{62}FN_6O_3Si$ $[M + H]^+$ 805.4637, found 805.4393.

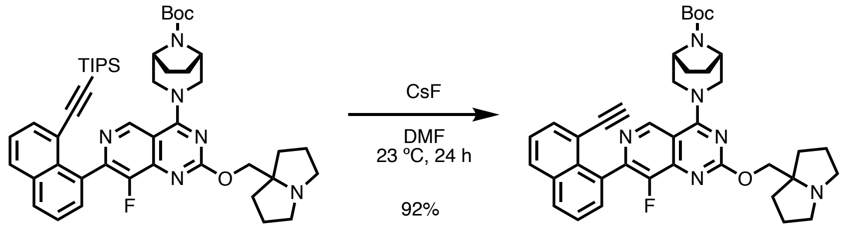


**S10** **S11**

***tert*-butyl (1*R*,5*S*)-3-(7-(8-ethynylnaphthalen-1-yl)-8-fluoro-2-((tetrahydro-1*H*-pyrrolizin-7a(5*H*)-yl)methoxy)pyrido[4,3-*d*]pyrimidin-4-yl)-3,8-diazabicyclo[3.2.1]octane-8-carboxylate (S11)**. A 100-mL round-bottom flask equipped with a stir bar was charged with *tert*-butyl 3-[8-fluoro-2-(1,2,3,5,6,7-hexahydropyrrolizin-8-ylmethoxy)-7-[8-(2-triisopropylsilylethynyl)-1-naphthyl]pyrido[4,3-d]pyrimidin-4-yl]-3,8-diazabicyclo[3.2.1]octane-8-carboxylate (1207.7 mg, 1.5000 mmol) and cesium fluoride (6835.5 mg, 45.000 mmol, 30 equiv) in DMF (15 mL) was stirred at ambient temperature for 24 h, at which point the full consumption of the starting material was judged by LC-MS. The reaction mixture was partitioned between EA (100 mL) and water (100 mL). The aqueous phase was further extracted with EA (50 mL x 3). The combined organic solution was sequentially washed with water (100 mL x 2) and brine (100 mL), and dried over anhydrous sodium sulfate. The solution was filtered and concentrated *in vacuo*. The residue was purified by flash column chromatography (0–30% MeOH*-DCM, *MeOH contains 2% ammonium hydroxide, 24-g RediSep(R) Rf column, Teledyne ISCO, Lincoln, NE). The desired product was obtained as a dark crystalline (896.4 mg,1.382 mmol, 92% yield). $^1$H NMR (400 MHz, $CDCl_3$) δ 8.96 (s, 1H), 7.93 (ddd, *J* = 11.9, 7.4, 2.2 Hz, 2H), 7.71 (dd, *J* = 7.2, 1.3 Hz, 1H), 7.58 – 7.54 (m, 2H), 7.42 (dd, *J* = 8.2, 7.2 Hz, 1H), 4.64 – 4.46 (m, 2H), 4.36 (s, 2H), 4.19 (d, *J* = 1.9 Hz, 2H), 3.65 (s, 2H), 3.09 (dt, *J* = 10.8, 5.6 Hz, 2H), 2.62 (dt, *J* = 10.2, 6.8 Hz, 2H), 2.51 (s, 1H), 2.12 – 2.01 (m, 2H), 2.01 – 1.55 (m, 15H), 1.50 (s, 9H). $^{13}$C NMR (100 MHz, $CDCl_3$) δ 165.76, 164.30,

153.47, 152.66, 150.06, 149.39, 149.27, 146.93, 146.79, 143.30, 143.24, 135.47, 134.31, 132.78, 132.75, 131.16, 130.45, 130.36, 130.03, 130.01, 125.59, 125.36, 118.57, 110.59, 110.57, 83.07, 81.55, 80.46, 73.37, 72.53, 55.73, 55.72, 54.19, 53.52, 36.76, 36.66, 36.64, 36.13, 36.11, 28.53, 27.85, 27.76, 25.65. $^{19}$F NMR (376 MHz, $CDCl_3$) δ -137.81. Accurate MS (ESI-TOF) calculated for $C_{38}H_{42}FN_6O_3$ $[M + H]^+$ 649.3302, found 649.3500.

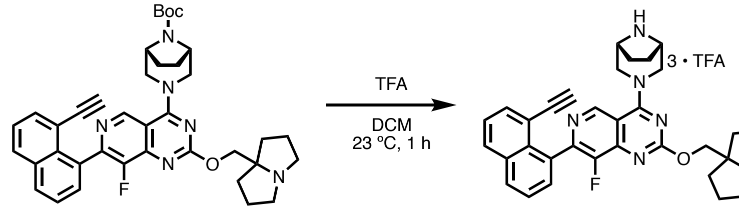


**S11** **S12**

**4-((1*R*,5*S*)-3,8-diazabicyclo[3.2.1]octan-3-yl)-7-(8-ethynylnaphthalen-1-yl)-8-fluoro-2-((tetrahydro-1*H*-pyrrolizin-7a(5*H*)-yl)methoxy)pyrido[4,3-*d*]pyrimidine • TFA salt (S12)**. A 20-mL scintillation vial equipped with a stir bar was charged with **S11** (300.0 mg, 0.462 mmol) and DCM (1.8 mL). The solution was cooled to 0 ºC. TFA (1.8 mL, 50 equiv) was added dropwise. The resulting mixture was stirred for 10 min at room temperature, when the starting material was found fully consumed by LC-MS. Volatiles were removed by rotary evaporation. The viscous residue was dropwise added into vigorously stirred ether pre-cooled to 0 ºC. Red solid precipitated from the reaction. The precipitate was aged in the etherous solution at 0 ºC for 1 h before centrifuged at 500 X g for 2 min to pellet. The supernatant was decanted. The pellet was resuspended and washed with 10-mL ether, and again centrifuged to separate. The solid was dried under high vacuum to a red crystalline (351.0 mg, 0.394 mmol, 85% yield). $^1$H NMR (400 MHz, DMSO) δ 10.41 (s, 1H), 9.51 (d, *J* = 10.2 Hz, 1H), 9.28 (d, *J* = 16.9 Hz, 1H), 9.17 (s, 1H), 8.16 (ddd, *J* = 11.0, 8.3, 1.4 Hz, 2H), 7.76 – 7.68 (m, 2H), 7.62 – 7.55 (m, 2H), 4.71 (d, *J* = 13.8 Hz, 2H), 4.62 – 4.51 (m, 3H), 4.23 (s, 2H), 3.89 (d, *J* = 13.7 Hz, 2H), 3.63 (s, 1H), 3.52 (dd, *J* = 11.9, 5.9 Hz, 2H), 3.29 – 3.17 (m, 2H), 2.23 – 1.90 (m, 14H). $^{19}$F NMR (376 MHz, DMSO) δ -74.07 (s, 9F), -139.34 (s, 1F). Accurate MS (ESI-TOF) calculated for $C_{33}H_{34}N_6FO$ $[M + H]^+$ 549.2778, found 549.2809. The solid material was directly used in the following steps without further purification.

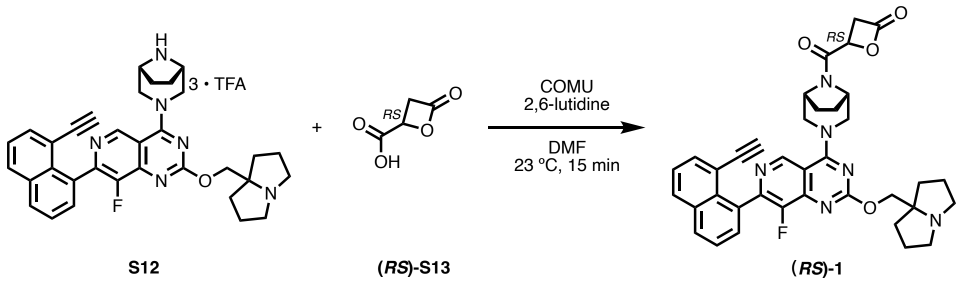


**S12** **(*RS*)-S13** **(*RS*)-1**

**(*RS*)-1 • 2TFA.** Racemic acid **(*RS*)-S13** was synthesized from (±)-malic acid via a known route[60]. A 4-mL dram vial equipped with a stir bar was charged with **(*RS*)-S13** (23.2 mg, 11.6 mmol, 3 equiv), COMU (42.8 mg, 0.1 mmol, 3 equiv). Anhydrous DMF (167 µL) and 2,6-lutidine (23 µL,

0.200 mmol, 6 equiv) were added. The resulting solution was stirred at room temperature for 15 min, before transferring to a new vial containing **S12** (25.9 mg, 0.033 mmol). The mixture was further stirred for 15 min, at which point the limiting agent **S12** was judged fully consumed. The reaction mixture was directly loaded onto a silica cartridge and purified by flash column chromatography (0–20% MeOH-DCM, 12-g RediSep Gold silica column, Teledyne ISCO, Lincoln, NE) to give a red solid (13.2 mg, 0.017 mmol, 50% yield). $^1$H NMR (600 MHz, Acetone) δ 9.16 – 9.06 (m, 1H), 8.12 (ddd, *J* = 13.9, 8.3, 1.4 Hz, 2H), 7.75 (dt, *J* = 7.1, 1.3 Hz, 1H), 7.73 – 7.67 (m, 1H), 7.63 (dt, *J* = 7.1, 1.7 Hz, 1H), 7.55 (ddd, *J* = 8.5, 7.1, 1.3 Hz, 1H), 5.54 – 5.41 (m, 1H), 4.91 – 4.57 (m, 7H), 3.91 – 3.71 (m, 5H), 3.25 (dq, *J* = 7.7, 3.3 Hz, 4H), 2.44 – 2.32 (m, 3H), 2.23 (h, *J* = 6.6 Hz, 5H), 2.19 – 2.11 (m, 3H), 2.00 – 1.84 (m, 3H). $^{13}$C NMR (151 MHz, Acetone) δ 168.07, 164.31, 164.29, 163.87, 161.89, 161.67, 161.45, 161.23, 135.75, 135.19, 135.18, 134.31, 131.93, 131.15, 131.09, 130.80, 126.58, 126.37, 121.16, 119.97, 119.20, 117.25, 115.29, 112.33, 84.48, 83.20, 79.88, 69.91, 66.42, 66.19, 56.40, 55.99, 55.96, 55.94, 55.90, 55.78, 55.70, 55.48, 55.32, 53.31, 53.26, 53.19, 43.76, 42.69, 42.32, 35.59, 35.54, 34.68, 28.30, 28.19, 28.07, 26.25, 26.19, 26.13, 24.98, 24.94, 24.90, 24.89. $^{19}$F NMR (564 MHz, Acetone) δ -75.59 (6F, 2TFA), -139.73 – -139.95 (m, 1F). Accurate MS (ESI-TOF) calculated for $C_{37}H_{36}FN_6O_4$ [M + H]+ 647.2782, found 647.2768.

Synthesis of (*RS*)-2

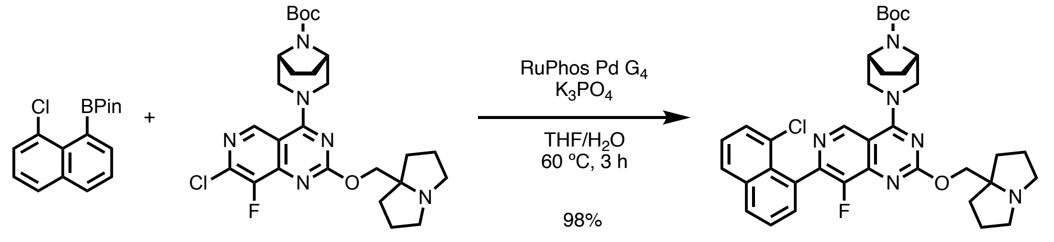


**S14** **S5** **S15**

***tert*-butyl (1*R*,5*S*)-3-(7-(8-chloronaphthalen-1-yl)-8-fluoro-2-((tetrahydro-1*H*-pyrrolizin-7a(5*H*)-yl)methoxy)pyrido[4,3-*d*]pyrimidin-4-yl)-3,8-diazabicyclo[3.2.1]octane-8-carboxylate (S15).** To a toluene solution (26.7 mL) of *tert*-butyl 3-[7-chloro-8-fluoro-2-(1,2,3,5,6,7-hexahydropyrrolizin-8-ylmethoxy)pyrido[4,3-d]pyrimidin-4-yl]-3,8-diazabicyclo[3.2.1]octane-8-carboxylate (**S5**, 1.4 g, 2.6 mmol) and 2-(8-chloronaphthalen-1-yl)-4,4,5,5-tetramethyl-1,3,2-dioxaborolane (**S14**, 1.14 g, 3.94 mmol, 1.5 equiv) was added 1.5 M aqueous solution of potassium phosphate (5.6 mL, 7.9 mmol). A stream of Argon was bubbled through this biphasic mixture for 5 min. RuPhos Pd G4 (241.8 mg, 0.263 mmol, 10 mol%) was added to this mixture. Sealed with a PTFE cap, the reaction mixture was stirred at 80 ºC for 5 h, at which point LC-MS analysis indicated full consumption of the starting material. The reaction mixture was cooled to 23 ºC. The layers were separated, and the aqueous layer was extracted with ethyl acetate (2 x 20 mL). The combined organic layers were dried over sodium sulfate. The dried solution was filtered, and the filtrate was concentrated. The residue was resuspended in 10 mL 50% MeCN-$H_2O$, and the pH was adjusted to 5 with 80% formic acid. This solution was loaded onto a Celite cartridge and purified by reverse phase flash column chromatography (10–50% MeCN-$H_2O$ with 0.1% formic acid, 50-g RediSep C18 Gold Column, Teledyne ISCO, Lincoln, NE). The product-containing fractions were pooled, and volatiles were removed under reduced pressure. The pH of the residual aqueous solution was adjusted to approximately 9 with saturated aqueous disodium hydrogen phosphate solution. The mixture was extracted with DCM (100 mL X 3). The combined organic layers were dried over anhydrous sodium sulfate. The dried solution was filtered and concentrated under reduced pressure to afford the product in di-TFA salt form as a yellow solid (977.0 mg, 1.482 mmol, 56% yield). $^{1}$H NMR (600 MHz, $CDCl_3$) δ 8.98 (s, 1H), 7.97 (dd, *J* = 8.0, 1.6 Hz, 1H), 7.85 (dd, *J* = 8.2, 1.3 Hz, 1H), 7.60 – 7.54 (m, 2H), 7.52 (dd, *J* = 7.5, 1.3 Hz, 1H), 7.39 (dd, *J* = 8.2, 7.4 Hz, 1H), 4.69 – 4.47 (m, 2H), 4.38 (s, 2H), 4.21 (s, 2H), 3.67 (s, 2H), 3.12 (dt, *J* = 11.0, 5.8 Hz, 2H), 2.64 (dt, *J* = 10.2, 6.9 Hz, 2H), 2.14 – 2.06 (m, 2H), 2.01 – 1.92 (m, 2H), 1.86 (dtd, *J* = 12.0, 6.4, 2.1 Hz, 4H), 1.81 – 1.73 (m, 2H), 1.67 (dt, *J* = 12.6, 7.6 Hz, 2H), 1.51 (s, 9H). $^{13}$C NMR (151 MHz, $CDCl_3$) δ 165.83, 165.82, 164.39, 151.25, 149.56, 149.53, 149.48, 147.65, 147.55, 143.46, 143.42, 136.05, 131.83, 131.81, 131.04, 131.03, 130.68, 130.62, 129.48, 128.78, 128.45, 126.08, 125.68, 110.58, 73.25, 72.73, 55.76, 55.75, 54.49, 54.07, 53.55, 36.13, 28.58, 28.56, 28.56, 25.67, 25.66. $^{19}$F NMR (564 MHz, $CDCl_3$) δ -137.22 – -137.68 (m, 1F).

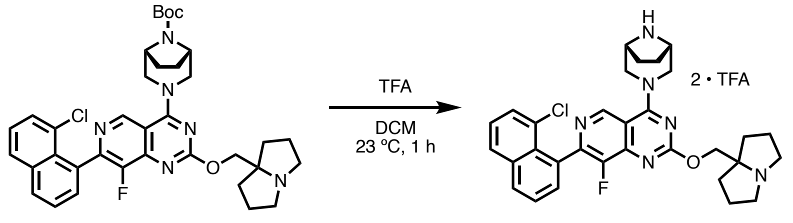


S15 S16

**4-(3,8-diazabicyclo[3.2.1]octan-3-yl)-7-(8-chloronaphthalen-1-yl)-8-fluoro-2-((tetrahydro-1*H*-pyrrolizin-7a(5*H*)-yl)methoxy)pyrido[4,3-*d*]pyrimidine • TFA salt (S16)**. A 20-mL scintillation vial was charged with *tert*-butyl 3-[7-(8-chloro-1-naphthyl)-8-fluoro-2-(1,2,3,5,6,7-hexahydropyrrolizin-8-ylmethoxy)pyrido[4,3-d]pyrimidin-4-yl]-3,8-diazabicyclo[3.2.1]octane-8-carboxylate (100.0 mg, 0.1521 mmol) and DCM (1 mL). The resulting solution was cooled to 0 ºC before TFA (1.0 mL, 13 mmol) was added. The mixture was stirred at autogenous temperature (0 to 23 ºC) for 15 min, at which point full consumption of the starting material was observed. Volatiles were removed by rotary evaporation. The viscous residue was dropwise added into vigorously stirred ether pre-cooled to 0 ºC. Red solid precipitated from the reaction. The precipitate was aged in the etherous solution at 0 ºC for 1 h before centrifuged at 500 X g for 2 min to pellet. The supernatant was decanted. The pellet was resuspended and washed with 10-mL ether, and again centrifuged to separate. The solid was dried under high vacuum to an off-white crystalline 7-(8-chloro-1-naphthyl)-4-(3,8-diazabicyclo[3.2.1]octan-3-yl)-8-fluoro-2-(1,2,3,5,6,7-hexahydropyrrolizin-8-ylmethoxy)pyrido[4,3-d]pyrimidine di-TFA salt (108.2 mg,0. 137 mmol, 91% yield). $^{1}$H NMR (400 MHz, DMSO) δ 10.49 (s, 1H), 9.57 (d, *J* = 9.4 Hz, 1H), 9.36 (s, 1H), 9.20 (s, 1H), 8.22 (dd, *J* = 8.3, 1.4 Hz, 1H), 8.11 (dd, *J* = 8.3, 1.4 Hz, 1H), 7.74 (dd, *J* = 8.2, 7.1 Hz, 1H), 7.66 (dd, *J* = 7.4, 1.3 Hz, 1H), 7.63 – 7.55 (m, 2H), 4.78 – 4.62 (m, 2H), 4.58 (d, *J* = 1.6 Hz, 2H), 4.22 (d, *J* = 11.2 Hz, 2H), 3.97 – 3.80 (m, 2H), 3.52 (dq, *J* = 12.1, 6.1 Hz, 2H), 3.28 – 3.16 (m, 2H), 2.23 – 1.87 (m, 13H). $^{19}$F NMR (376 MHz, Acetone) δ -74.41 (6F, 2•TFA), -139.68 (s, 1F). Accurate MS (ESI-TOF) calculated for $C_{31}H_{33}ClFN_6O$ $[M + H]^+$ 559.2388, found 559.2377. The material was used for the next steps without further purification.

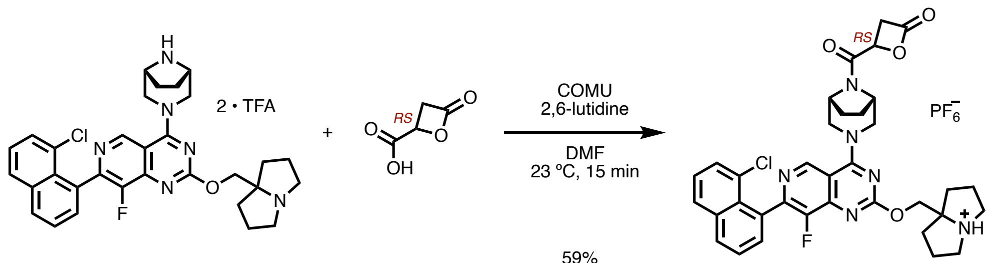

**S16** **(*RS*)-S13** **(*RS*)-2**

**(*RS*)-2 • hexafluorophosphate.** A 4-mL dram vial equipped with a stir bar was charged with **(*RS*)-S13** (5.9 mg, 0.051 mmol, 3 equiv), COMU (21.9 mg, 0.051 mmol, 3 equiv). DMF (150 µL) and 2,6-lutidine (11.8 µL, 0.100 mmol, 6 equiv) were added. The mixture was stirred at room temperature for 15 min before **S16** (15.3 mg, 0.017 mmol, 1 equiv) was added as solid. The resulting solution was further stirred at room temperature for 15 min, at which point the consumption of limiting agent **S16** was complete. The reaction mixture was directly loaded onto a silica gel cartridge, and purified by flash column chromatography (0–20% MeOH-DCM, 12-g Gold RediSep(R) Rf column, Teledyne ISCO, Lincoln, NE) to give the title compound as a white solid (8.9 mg, 0.010 mmol, 59% yield). $^{1}$H NMR (600 MHz, Acetone) δ 9.26 – 9.18 (m, 1H), 8.21 (dd, *J* = 8.3, 1.3 Hz, 1H), 8.10 (dd, *J* = 8.2, 1.3 Hz, 1H), 7.74 (dd, *J* = 8.2, 7.1 Hz, 1H), 7.65 (ddd, *J* = 7.1, 2.8, 1.3 Hz, 2H), 7.58 (dd, *J* = 8.2, 7.4 Hz, 1H), 5.62 (s, 1H), 5.52 – 5.42 (m, 1H), 4.91 – 4.64 (m, 6H), 3.95 – 3.77 (m, 6H), 3.46 (dd, *J* = 12.1, 6.4 Hz, 2H), 2.45 – 2.37 (m, 2H), 2.30 (dt, *J* = 12.8, 6.7 Hz, 2H), 2.25 – 2.08 (m, 5H), 2.03 – 1.84 (m, 3H). $^{13}$C NMR (151 MHz, Acetone) δ 170.91, 168.00, 167.93, 166.81, 166.75, 164.72, 163.92, 163.64, 151.58, 149.88, 148.84, 148.75, 145.30, 145.25, 145.19, 145.15, 136.92, 132.56, 132.05, 131.66, 130.95, 130.34, 129.69, 129.26, 127.29, 126.74, 112.03, 82.03, 71.11, 66.42, 66.19, 60.54, 56.40, 56.35, 56.32, 55.61, 55.37, 55.04, 53.12, 53.07, 42.61, 42.23, 35.00, 34.90, 28.19, 28.12, 28.08, 26.13, 26.07, 25.26, 25.23, 25.21, 20.82, 14.50. $^{19}$F NMR (564 MHz, Acetone) δ -72.72 (d, J = 705 Hz, 6F, P$\underline{F_6}$), -141.23 – -141.36 (m, 1F). Accurate MS (ESI-TOF) calculated for $C_{35}H_{35}ClFN_6O_4$ $[M + H]^+$ 657.2392, found 657.2383.

Synthesis of (R)-2

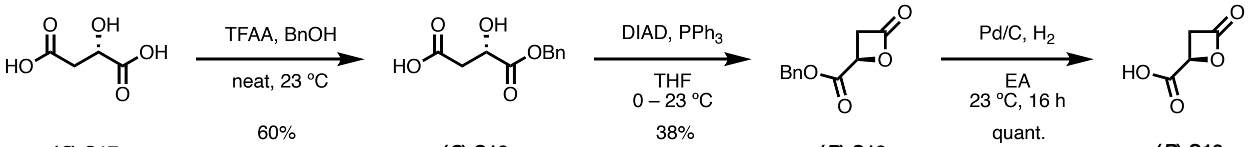

**(*S*)-S17** **(*S*)-S18** **(*R*)-S19** > 20:1 er (NMR) **(*R*)-S13**

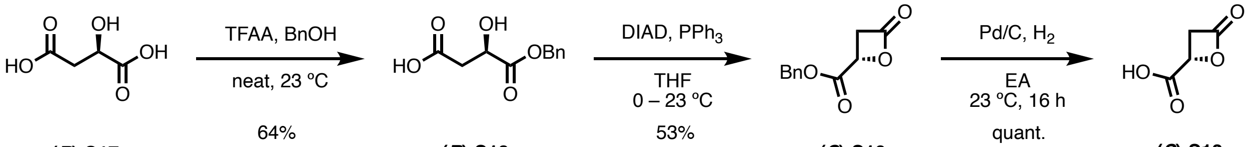

**(*R*)-S17** **(*R*)-S18** **(*S*)-S19** > 20:1 er (NMR) **(*S*)-S13**

**(*R*)-4-(benzyloxy)-3-hydroxy-4-oxobutanoic acid [(*R*)-S13].** Synthesis of **(*R*)-S13** and its enantiomer **(*S*)-S13** was adapted from known procedures in the literature[60]. Enantiomeric ratio (er) of both synthetic samples were determined by $^1$H NMR of respective benzyl ester precursor **(*R*/*S*)-S19** in the presence of Eu(hfc)$_3$ NMR shift reagent and were referenced to synthetic **(*RS*)-S19** from (±)-malic acid. With 1 equiv of Eu(hfc)$_3$ reagent in $CDCl_3$ (20 mM), the racemate benzylic protons were separated by 0.05 ppm with no overlap of each enantiomer's major doublet peaks. Both synthetic samples from above scheme showed single enantiomer peaks [5.27 ppm without Eu(hfc)$_3$, 5.72 ppm for *R*-isomer with Eu(hfc)$_3$, 5.67 ppm for *S*-isomer with Eu(hfc)$_3$] on $^1$H NMR with Eu-shift reagents. The er of both samples were estimated >20:1.

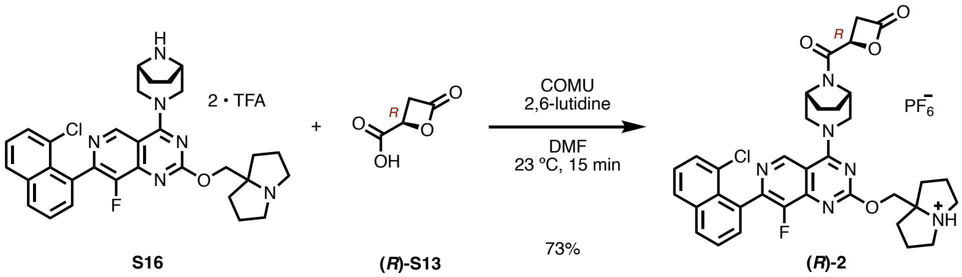


**S16** **(*R*)-S13** **(*R*)-2**

**(*R*)-2 • hexafluorophosphate.** Like the racemate synthesis, **S16** (26.2 mg, 0.033 mmol, 1 equiv), **(*R*)-S13** (11.6 mg, 0.100 mmol, 3 equiv), COMU (42.8 mg, 0.100 mmol, 3 equiv), 2,6-lutidine (23.1 µL, 0.200 mmol, 6 equiv) afforded the title product in DMF as white solid (21.4 mg, 0.024 mmol, 73% yield). $^1$H NMR was identical to that of the racemate. Accurate MS (ESI-TOF) calculated for $C_{35}H_{35}ClFN_6O_4$ $[M + H]^+$ 657.2392, found 657.2383.

Synthesis of (S)-2

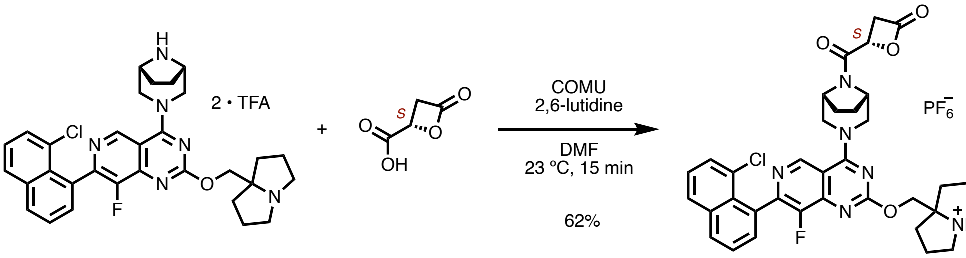


**S16** **(*S*)-S13** **(*S*)-2**

**(*S*)-2 • hexafluorophosphate.** Like the racemate synthesis, **S16** (26.2 mg, 0.033 mmol, 1 equiv), **(*S*)-S13** (11.6 mg, 0.100 mmol, 3 equiv), COMU (42.8 mg, 0.100 mmol, 3 equiv), 2,6-lutidine (23.1 µL, 0.200 mmol, 6 equiv) afforded the title product in DMF as white solid (18.2 mg, 0.021 mmol, 62% yield). $^1$H NMR was identical to that of the racemate. Accurate MS (ESI-TOF) calculated for $C_{35}H_{35}ClFN_6O_4$ $[M + H]^+$ 657.2392, found 657.2383.

Synthesis of (2R, 3S)-3

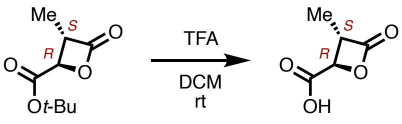


**(2*R*, 3*S*)-S20** **(2*R*, 3*S*)-S21**

**(2*R*,3*S*)-3-methyl-4-oxooxetane-2-carboxylic acid [(2*R*, 3*S*)-S21].** Precursor **(2*R*, 3*S*)-S20** was synthesized using known procedure in the literature[61] using enantiopure commercial material. A 20-mL scintillation vial equipped with a stir bar was charged with **(2*R*, 3*S*)-S20** (20.0 mg, 0.107 mmol). DCM (164 μL) and TFA (164 μL) were added. The resulting solution was stirred at room temperature for 30 min, at which point the consumption of the starting material was judged complete by TLC. The reaction mixture was concentrated <30 ºC under reduced pressure. The residue, **(2*R*,3*S*)-S21**, was used in the next step without further purification.

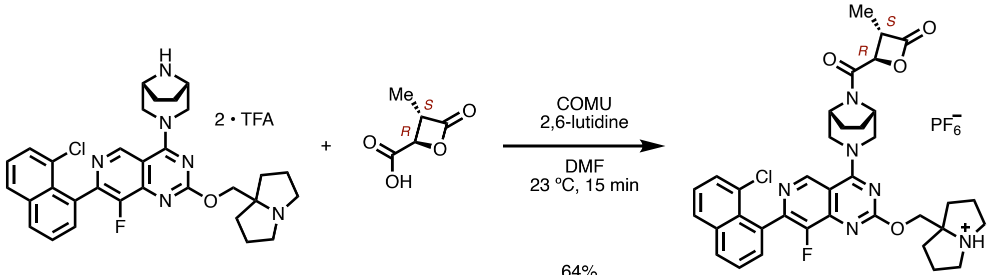


**S16** **(2*R*, 3*S*)-S21** **(2*R*, 3*S*)-3**

**(2*R*,3*S*)-3 • hexafluorophosphate.** A 4-mL dram vial equipped with a stir bar was charged with **(2*R*,3*S*)-S21** (11.7 mg, 0.090 mmol, 3 equiv), COMU (38.5 mg, 0.090 mmol, 3 equiv). DMF (149 μL) and 2,6-lutidine (20.9 μL, 0.189 mmol, 6 equiv) were added. The mixture was stirred at room temperature for 15 min before **S16** (27.0 mg, 0.030 mmol, 1 equiv) was added as solid. The resulting solution was further stirred at room temperature for 15 min, at which point the consumption of limiting agent **S16** was complete. The reaction mixture was directly loaded onto a silica gel cartridge, and purified by flash column chromatography (0–20% MeOH-DCM, 12-g Gold RediSep(R) Rf column, Teledyne ISCO, Lincoln, NE) to give the title compound as a white solid (15.3 mg, 0.019 mmol, 62% yield). $^{1}$H NMR (600 MHz, Acetone) δ 9.27 – 9.16 (m, 1H), 8.21 (dd, *J* = 8.2, 1.3 Hz, 1H), 8.10 (dd, *J* = 8.3, 1.3 Hz, 1H), 7.74 (dd, *J* = 8.2, 7.1 Hz, 1H), 7.65 (dd, *J* = 7.0, 1.3 Hz, 2H), 7.58 (dd, *J* = 8.2, 7.4 Hz, 1H), 5.30 – 5.15 (m, 1H), 4.91 – 4.62 (m, 6H), 4.10 (td, *J* = 7.2, 3.6 Hz, 1H), 4.00 – 3.76 (m, 5H), 3.53 – 3.42 (m, 2H), 2.46 – 2.37 (m, 2H), 2.35 – 2.27 (m, 2H), 2.27 – 2.16 (m, 4H), 2.09 (s, 3H), 1.55 – 1.44 (m, 3H). $^{13}$C NMR (151 MHz, Acetone) δ 171.55, 171.50, 166.74, 164.67, 163.79, 163.57, 151.55, 149.85, 148.80, 148.72, 148.27, 148.19, 145.26, 145.22, 145.17, 145.13, 136.88, 132.52, 132.06, 131.64, 130.92, 130.33, 129.67, 129.23, 127.28, 126.74, 112.00, 82.27, 73.71, 73.49, 70.99, 56.38, 56.31, 56.28, 56.22, 55.57, 55.21, 53.09, 53.00, 50.97, 50.53, 50.51, 34.92, 34.89, 34.82, 28.21, 28.16, 28.10, 28.05, 26.12, 26.06, 25.95, 25.90, 25.19, 25.15, 25.13, 12.43, 12.34. $^{19}$F NMR (564 MHz, Acetone) δ -73.25 (d, P<u>F6</u>, 6F), -141.27 – -141.37 (m, 1F). Accurate MS calculated for $C_{36}H_{37}ClFN_6O_4$ $[M + H]^+$ 671.2549, found 671.2585.

Synthesis of (2R, 3S)-4

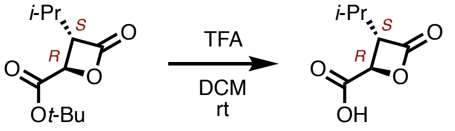


**(2*R*, 3*S*)-S22** **(2*R*, 3*S*)-S23**

**(2*R*,3*S*)-3-isopropyl-4-oxooxetane-2-carboxylic acid [(2*R*, 3*S*)-S23].** Precursor **(2*R*, 3*S*)-S22** was synthesized using known procedure in the literature[62] using enantiopure commercial material. A 20-mL scintillation vial equipped with a stir bar was charged with **(2*R*, 3*S*)-S22** (214.3 mg, 0.350 mmol, 1 equiv). DCM (0.54 mL) and TFA (0.54 mL) were added. The resulting solution was stirred at room temperature for 30 min, at which point the consumption of the starting material was judged complete by TLC. The reaction mixture was concentrated <30 ºC under reduced pressue. The residue, **(2*R*, 3*S*)-S23**, was used in the next step without further purification.

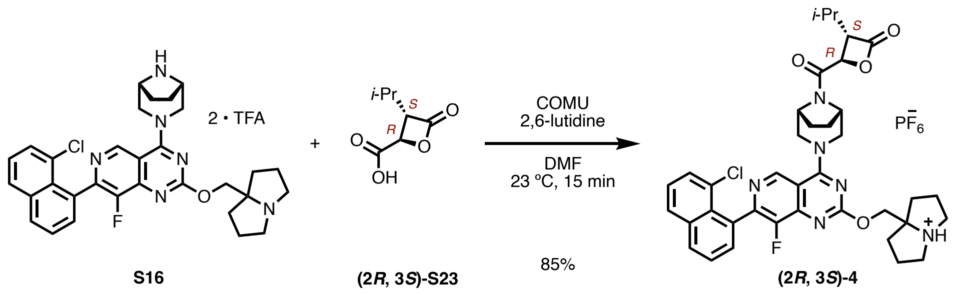


**(2*R*,3*S*)-4 • hexafluorophosphate.** A 4-mL dram vial equipped with a stir bar was charged with **(2*R*,3*S*)-S23** (14.2 mg, 0.090 mmol, 3 equiv), COMU (38.5 mg, 0.090 mmol, 3 equiv). DMF (149 μL) and 2,6-lutidine (20.9 μL, 0.189 mmol, 6 equiv) were added. The mixture was stirred at room temperature for 15 min before **S16** (27.0 mg, 0.030 mmol, 1 equiv) was added as solid. The resulting solution was further stirred at room temperature for 15 min, at which point the consumption of limiting agent **S16** was complete. The reaction mixture was directly loaded onto a silica gel cartridge, and purified by flash column chromatography (0–20% MeOH-DCM, 12-g Gold RediSep(R) Rf column, Teledyne ISCO, Lincoln, NE) to give the title compound as a white solid (21.3 mg, 0.025 mmol, 84% yield). $^{1}$H NMR (600 MHz, Acetone) δ 9.27 – 9.15 (m, 1H), 8.20 (dd, *J* = 8.3, 1.3 Hz, 1H), 8.09 (dd, *J* = 8.3, 1.3 Hz, 1H), 7.74 (dd, *J* = 8.2, 7.1 Hz, 1H), 7.69 (t, *J* = 7.7 Hz, 1H), 7.67 – 7.63 (m, 2H), 7.57 (dd, *J* = 8.2, 7.4 Hz, 1H), 7.16 (dp, *J* = 7.8, 0.6 Hz, 2H), 5.43 – 5.19 (m, 2H), 4.93 – 4.70 (m, 7H), 3.95 (ddd, *J* = 8.1, 4.1, 1.1 Hz, 1H), 3.91 – 3.78 (m, 3H), 3.47 (dd, *J* = 11.9, 6.1 Hz, 2H), 2.50 (d, *J* = 0.6 Hz, 7H), 2.46 – 2.37 (m, 2H), 2.35 – 2.27 (m, 2H), 2.25 – 2.15 (m, 5H), 2.09 (s, 8H), 1.15 – 1.03 (m, 8H). $^{13}$C NMR (151 MHz, Acetone) δ 170.94, 170.39, 170.30, 166.71, 164.62, 163.67, 157.51, 157.51, 151.56, 149.86, 148.75, 148.66, 148.34, 148.26, 145.24, 145.20, 139.20, 136.87, 132.50, 132.07, 132.06, 131.62, 130.92, 130.31, 129.65, 129.22, 127.26, 126.73, 121.74, 121.71, 111.98, 82.18, 70.95, 70.88, 70.80, 70.68, 64.34, 62.42, 62.36, 62.34, 56.35, 56.29, 56.26, 55.68, 55.43, 53.19, 53.09, 53.08, 34.92, 34.84, 30.59, 28.55, 28.09, 28.04, 27.92, 26.07, 26.02, 25.16, 25.12, 25.10, 23.45, 23.44, 21.07, 20.62, 20.45, 20.28, 19.71, 19.57. $^{19}$F NMR (564 MHz, Acetone) δ -72.54 (d, P<u>F6</u>, 6F), -141.14 – -141.23 (m, 1F). Accurate MS calculated for $C_{38}H_{41}ClFN_6O_4$ [M + H]$^+$ 699.2862, found 699.2878.

Synthesis of (2R, 3S)-5

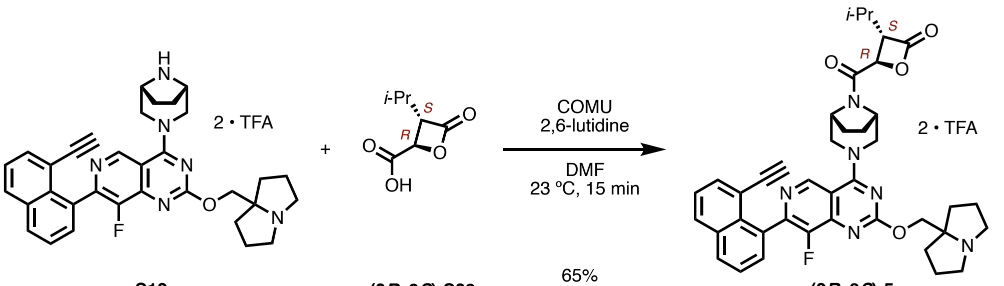


**S12** **(2*R*, 3*S*)-S23** **(2*R*, 3*S*)-5**

**[(2*R*, 3*S*)-5] • 2TFA.** An oven-dried one-dram vial was charged with **(2*R*, 3*S*)-S23** (6.4 mg, 0.040 mmol, 3 equiv), COMU (17.3 mg, 0.040 mmol, 3 equiv) and a magnetic stir bar. DMF (135 μL) and 2,6-lutidine (9.4 μL, 0.081 mmol, 6 equiv) were added sequentially. The reaction mixture was stirred for 15 min at room temperature. **S12** (12.0 mg, 0.013 mmol) was added as a solid, and the resulting dark solution was stirred at room temperature for 15 min, at which point full consumption of **S12** was judged by LC-MS. The reaction mixture was directly loaded onto purified by column chromatography (0–30% methanol–dichloromethane, 12-g RediSep(R) Rf Gold column, Teledyne ISCO, Lincoln, NE). The title compound was isolated as a red solid (29.7 mg, 0.032 mmol, 65% yield). $^{1}$H NMR (600 MHz, Acetone) δ 12.16 – 11.71 (m, 2H), 9.28 – 9.13 (m, 1H), 8.15 (ddd, *J* = 17.9, 8.3, 1.3 Hz, 2H), 7.78 (dd, *J* = 7.1, 1.3 Hz, 1H), 7.72 (dd, *J* = 8.2, 7.1 Hz, 1H), 7.65 (dd, *J* = 7.1, 1.4 Hz, 1H), 7.58 (dd, *J* = 8.3, 7.1 Hz, 1H), 5.39 – 5.26 (m, 2H, overlapping with residual MeOH), 4.91 – 4.72 (m, 2H, overlapping with residual MeOH), 4.01 – 3.90 (m, 2H), 3.91 – 3.74 (m, 4H), 3.38 (s, 2H), 3.26 (d, *J* = 3.8 Hz, 1H), 2.39 (t, *J* = 6.2 Hz, 3H), 2.34 – 2.15 (m, 8H), 2.00 (s, 3H), 1.33 – 1.22 (m, 1H), 1.22 – 0.98 (m, 7H). $^{13}$C NMR (151 MHz, Acetone) δ 170.59, 166.95, 164.71, 164.15, 163.89, 163.85, 160.25, 160.00, 159.76, 151.32, 147.76, 145.06, 144.93, 136.10, 135.37, 133.73, 131.99, 131.67, 131.64, 131.45, 131.11, 129.13, 128.62, 126.80, 126.70, 125.15, 119.90, 118.24, 116.32, 112.41, 84.62, 83.50, 81.37, 71.02, 70.91, 70.75, 64.72, 56.70, 56.48, 56.27, 55.98, 55.91, 55.73, 55.47, 55.30, 55.03, 53.49, 53.38, 53.32, 53.17, 35.45, 35.34, 28.45, 28.32, 28.16, 26.39, 26.29, 25.26, 25.23, 25.14, 20.84, 20.67, 19.93, 19.86, 19.79. $^{19}$F NMR (564 MHz, Acetone) δ -76.35 (6F, 2TFA), -140.26 – -140.53 (m, 1F). Accurate MS calculated for $C_{40}H_{42}FN_6O_4$ [M + H]+ 689.3252, found 689.3256.

Synthesis of (2R, 3S)-6

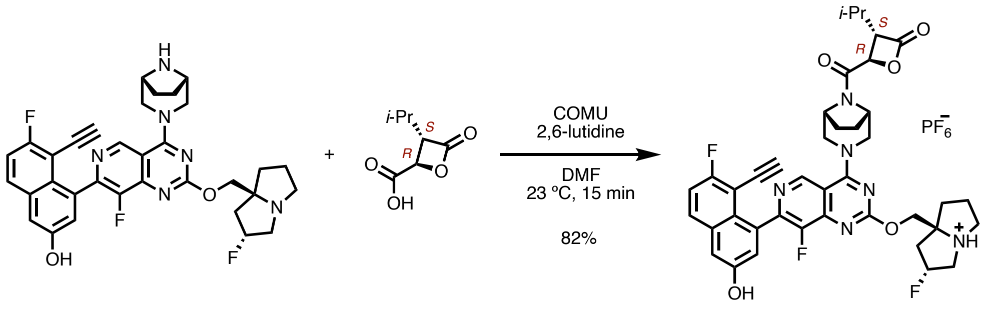


**S24** **(2*R*, 3*S*)-S23** **(2*R*, 3*S*)-6**

**(2*R*, 3*S*)-6 • hexafluorophosphate. S24** was synthesized using a known procedure[59]. A 4-mL dram vial equipped with a stir bar was charged with **(2*R*, 3*S*)-S23** (15.8 mg, 0.100 mmol), COMU (42.8 mg, 0.100 mmol, 3 equiv) and a magnetic stir bar. DMF (166 µL) and 2,6-lutidine (23 µL, 0.200 mmol, 6 equiv) were added sequentially via pipette. The reaction mixture was stirred for 15 min at 23 ºC, during which the color of the solution turned dark brown. **S24** (20.0 mg, 0.033 mmol) was added as a solid, and the resulting dark solution was stirred at room temperature for 15 min, at which point full consumption of **S12** was judged by LC-MS. The reaction mixture was directly loaded onto purified by column chromatography (0–30% methanol–dichloromethane, 12-g RediSep(R) Rf Gold column, Teledyne ISCO, Lincoln, NE). The title compound was isolated as an off-white solid (24.4 mg, 0.028 mmol, 83% yield).

$^{1}$H NMR (600 MHz, Acetone) δ 9.22 – 9.11 (m, 1H), 7.97 (dd, *J* = 9.1, 5.8 Hz, 1H), 7.48 (dd, *J* = 2.7, 1.2 Hz, 1H), 7.41 (t, *J* = 9.0 Hz, 1H), 7.31 (t, *J* = 2.4 Hz, 1H), 5.62 (s, 2H), 5.37 – 5.26 (m, 1H), 4.94 – 4.65 (m, 6H), 4.05 – 3.75 (m, 6H), 3.54 – 3.41 (m, 2H), 2.75 – 2.54 (m, 2H), 2.46 (d, *J* = 11.0 Hz, 1H), 2.30 (s, 2H), 2.21 (dd, *J* = 13.9, 7.1 Hz, 1H), 2.17 – 2.07 (m, 2H), 2.03 – 1.82 (m, 3H), 1.16 – 1.02 (m, 6H). $^{13}$C NMR (151 MHz, Acetone) δ 170.36, 170.32, 166.65, 164.69, 164.43, 164.41, 163.94, 163.68, 163.65, 162.78, 162.76, 155.07, 155.06, 152.49, 150.79, 147.94, 147.02, 146.93, 145.07, 135.03, 135.00, 133.72, 131.53, 131.46, 126.75, 123.90, 123.85, 116.92, 116.75, 112.80, 112.29, 105.15, 105.13, 105.04, 105.02, 96.65, 96.58, 95.48, 95.41, 90.48, 90.44, 90.40, 90.35, 75.93, 75.90, 71.46, 70.74, 70.64, 62.39, 62.36, 62.33, 60.57, 60.48, 60.44, 60.35, 58.31, 58.28, 56.49, 56.06, 55.69, 55.60, 55.42, 55.35, 54.93, 53.18, 53.09, 53.03, 41.42, 41.27, 41.08, 35.81, 35.65, 28.18, 28.06, 28.00, 27.88, 26.13, 26.00, 25.79, 25.78, 25.71, 25.70, 20.59, 20.42, 19.67, 19.53, 1.07. $^{19}$F NMR (376 MHz, Acetone) δ -71.64 (d, 6F, $PF_6$), -111.14 – -111.19 (m, 1F), -141.15 – -141.16 (m, 1F), -174.12 – -174.46 (m, 1F). Accurate MS calculated for $C_{40}H_{40}F_3N_6O_5$ [M + H]$^+$ 741.3012, found 741.3011.

Synthesis of (R)-7

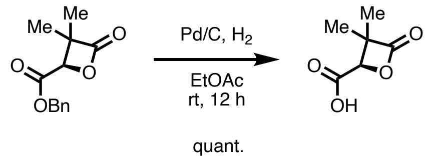


(*R*)-S25 (*R*)-S26

**(*R*)-3,3-dimethyl-4-oxooxetane-2-carboxylic acid [(*R*)-S26].** Precursor **(*R*)-S25** was synthesized using a known method form optically pure diethyl (*S*)-malate[63]. The enantiomeric excess (e.e.) of precursor **(*R*)-S25** was measured by chiral HPLC 97.5%, $t_R$(*R*) 1.840 min [$t_R$(*S*) 1.704 min, CHIRALPAK IC-3 column, Gradient (B%, MeOH containing 20 mM $NH_3$) 5 to 20% in 4 min, hold at 20% for 2 min, temperature 25 ºC, Flow Rate 2 mL/min]. The absolute configuration was determined by comparing optical rotation, $[\alpha]^{20}$ = -7.8 (*c* = 0.1 g/100 mL MeOH), with reported value in the literature[63].

A 4-mL dram vial equipped with a stir bar was charged with Pd/C (2.7 mg, 10 mol%) and **(*R*)-S25** (11.7 mg, 0.050 mmol). Ethyl acetate (1 mL) was added. The solution was bubbled with a stream of hydrogen gas for 10 min, and stirred under hydrogen atmosphere for 16 h. The reaction mixture was filtered through a plug of Celite® before concentrated to give crude **(*R*)-S26** as a colorless oil. The crude was directly used in the next amide coupling.

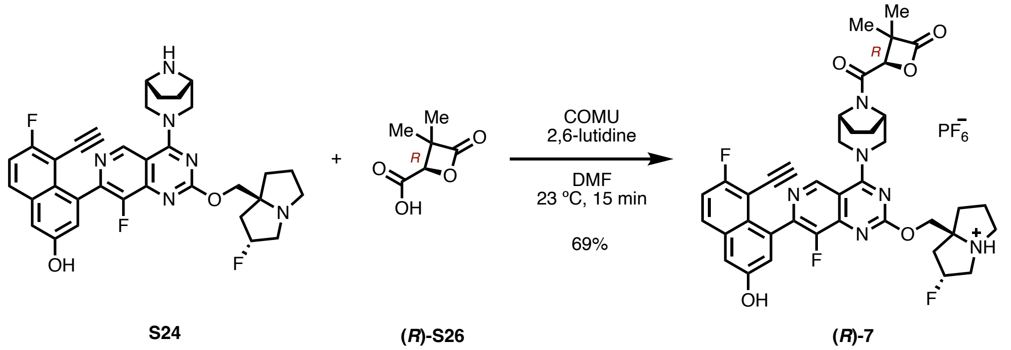


**(*R*)-7 • hexafluorophosphate.** A 4-mL dram-vial was charged with crude **(*R*)-S26**, COMU (21.2 mg, 0.0500 mmol, 3 equiv). Anhydrous DMF (165 μL), and 2,6-lutidine (11.5 μL, 0.100 mmol, 6.0 equiv) were added via pipette. The reaction mixture was stirred for 15 min at 23 ºC, during which the color of the solution turned dark brown. Amine **S24** (MRTX1133) (19.8 mg, 0.0333 mmol) was added as a solid. The mixture was further stirred at room temperature for 15 min before purified by flash column chromatography (0–30% methanol–dichloromethane, 12-g RediSep(R) Rf Gold column, Teledyne ISCO, Lincoln, NE) to give the title compound as an off-white solid (19.8 mg, 0.023 mmol, 69% yield). $^{1}$H NMR (600 MHz, Acetone) δ 9.17 – 9.08 (m, 1H), 7.93 (ddd, *J* = 9.1, 5.8, 1.9 Hz, 1H), 7.44 (t, *J* = 2.9 Hz, 1H), 7.37 (t, *J* = 8.9 Hz, 1H), 7.29 – 7.24 (m, 1H), 5.62 – 5.43 (m, 1H), 5.40 – 5.23 (m, 1H), 4.99 – 4.43 (m, 6H), 4.11 – 3.60 (m, 6H), 3.49 – 3.34 (m, 2H), 2.66 – 2.46 (m, 2H), 2.40 (t, *J* = 8.7 Hz, 1H), 2.24 (dh, *J* = 13.2, 6.0 Hz, 2H), 2.13 – 2.05 (m, 2H), 1.95 – 1.81 (m, 3H), 1.58 (d, *J* = 9.7 Hz, 3H), 1.38 (d, *J* = 2.4 Hz, 1H), 1.29 – 1.14 (m, 3H). $^{13}$C NMR (151 MHz, Acetone) δ 174.42, 174.25, 170.86, 166.61, 166.56, 164.64, 164.62, 164.35, 164.33, 163.08, 162.82, 162.71, 162.69, 155.02, 155.01, 152.49, 150.78, 150.73, 148.11, 148.05, 147.98, 146.92, 146.87, 146.84, 146.78, 145.12, 145.08, 144.95, 144.91, 135.01, 134.96, 133.66, 131.45, 131.39, 126.74, 126.69, 123.84, 123.80, 123.78, 116.84, 116.67, 112.71, 112.23, 112.16, 105.09, 104.98, 96.78, 96.71, 95.61, 95.54, 90.39, 90.35, 90.32, 90.28, 80.31, 78.03, 78.00, 77.15, 77.13, 75.88, 75.86, 75.83, 71.58, 60.57, 60.53, 60.50, 60.47, 60.45, 60.43, 60.40, 60.36, 60.33, 58.21, 58.18, 57.51, 57.39, 57.37, 57.00, 56.50, 56.18, 55.59, 55.34, 55.23, 55.07, 54.98, 54.89, 54.81, 52.97, 52.89, 52.86, 52.80, 41.52, 41.38, 41.33, 41.21, 35.81, 35.66, 28.54, 28.41, 28.37, 28.25, 26.42, 26.27, 25.74, 25.73, 25.66, 25.53, 25.41, 21.82, 21.76, 21.75, 20.73, 18.13, 17.62, 14.38. $^{19}$F NMR (564 MHz, Acetone) δ -72.65 (d, 6F, $PF_6$), -111.16 – -111.24 (m, 1F), -141.02 – -141.20 (m, 1F), -174.15 – -174.35 (m, 1F). Accurate MS (ESI-TOF) calculated for $C_{39}H_{38}F_3N_6O_5$ [M + H]$^+$ 727.2856, found 727.2836.

**Gram-scale synthesis of free-base (*R*)-7.** An oven-dried 100-mL round bottom flask was charged with COMU (2137.9 mg, 4.992 mmol) and DMF (20.8 mL). **(*R*)-S26** (719.5 mg, 4.992 mmol) and 2,6-lutidine (1.45 mL, 12.5 mmol) were added sequentially via syringe. The reaction mixture was stirred at 23 ºC for 10 min. MRTX1133 **S34** (2498.6 mg, 4.160 mmol) was added as a solid in one portion. The mixture was further stirred at room temperature for 15 min before quenched with water. The reaction mixture was partitioned between 5% sodium bicarbonate (200 mL) and EA (200 mL). The organic phase was saved and the aqueous phase was further extracted with EA (200 mL X 2). The combined organic solution was washed with half-saturated sodium chloride solution twice,  saturated sodium chloride solution, and dried over anhydrous sodium sulfate. The dried solution was filtered and concentrated. The crude was purified by flash column chromatography (0–30% MeOH-DCM, 120-g Gold RediSep(R) column, Teledyne ISCO, Lincoln, NE) to give the title free-base form of **(*R*)-7** as a yellow solid (2.1620 g, 2.975 mmol, 72% yield). NMR spectra were consistent with the PF6 salt obtained in smaller-scale. Melting point (ether): 144 ºC (decomposed).

Syntheses of Miscellaneous Compounds

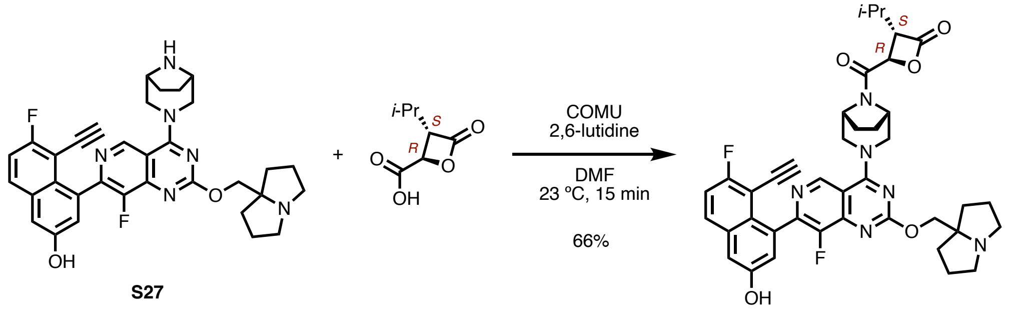


**S27** **(2*R*, 3*S*)-S23** **(2*R*, 3*S*)-5a**

**(2*R*,3*S*)-5a.** Synthesis of precursor **S27** was adapted from a known procedure [58]. Quality control was performed for the synthetic material by comparing the proton NMR to reported characterizations and the observed m/z to theoretical mass. $^1$H NMR (400 MHz, DMSO-d6) δ 10.13 (s, 1H), 9.02 (s, 1H), 7.97 (dd, *J* = 9.2, 5.6 Hz, 1H), 7.48 - 7.46 (m, 1H), 7.38 (d, *J* = 2.4 Hz, 1H), 7.17 (d, *J* = 2.4 Hz, 1H), 4.48 (d, *J* = 11.6 Hz, 1H), 4.29 (d, *J* = 12.0 Hz, 1H), 4.02 (s, 2H), 3.93 (d, *J* = 1.2 Hz, 1H), 3.64 (d, *J* = 12.0 Hz, 1H), 3.61 - 3.54 (m, 3H), 2.94 - 2.91 (m, 2H), 2.55 (d, J = 7.2 Hz, 2H), 1.90 - 1.86 (m, 2H), 1.83 - 1.77 (m, 4H), 1.75 - 1.65 (m, 4H), 1.60 - 1.57 (m, 2H). LC-MS (ESI-Q) calculated for $C_{33}H_{32}F_2N_6O_2$ [M + H]$^+$ 583.26, found 583.15. A 4-mL dram-vial equipped with a stir bar was charged with **(2*R*,3*S*)-S23**, and COMU (8.8 mg, 0.021 mmol). DMF (200 µL) and 2,6-lutidine (5.5 mg, 0.051 mmol) were added. The mixture was stirred at room temperature for 10 min until it turned yellow. Amine **S27** (10.0 mg, 0.017 mmol) was added as a solid. The mixture was further stirred at room temperature for 15 min when the conversion of the starting material was apparent in LC-MS. The reaction mixture without work-up was directly loaded onto a tightly packed silica cartridge and purified by flash column chromatography (0–30% MeOH-DCM, 12-g Gold RediSep(R) Rf column, Teledyne ISCO, Lincoln, NE). The title compound **(2*R*,3*S*)-5a** was isolated as a yellow solid (8.2 mg,0.011 mmol, 66% yield). $^1$H NMR (600 MHz, DMSO) δ 10.19 (s, 2H), 9.12 (dd, *J* = 8.2, 5.1 Hz, 1H), 7.99 (dd, *J* = 9.2, 5.9 Hz, 1H), 7.48 (t, *J* = 9.0 Hz, 1H), 7.41 (d, *J* = 2.5 Hz, 1H), 7.18 (d, *J* = 2.7 Hz, 1H), 5.44 (q, *J* = 5.1 Hz, 1H), 4.83 – 4.44 (m, 6H), 3.97 – 3.81 (m, 2H), 3.81 – 3.63 (m, 2H), 3.49 (s, 2H), 3.22 (s, 2H), 2.23 – 1.72 (m, 12H), 1.23 (s, 1H), 1.13 – 0.98 (m, 5H).Accurate MS (ESI-QTOF) calculated for $C_{40}H_{41}F_2N_6O_5$ [M + H]$^+$ 723.3107, found 723.3104.

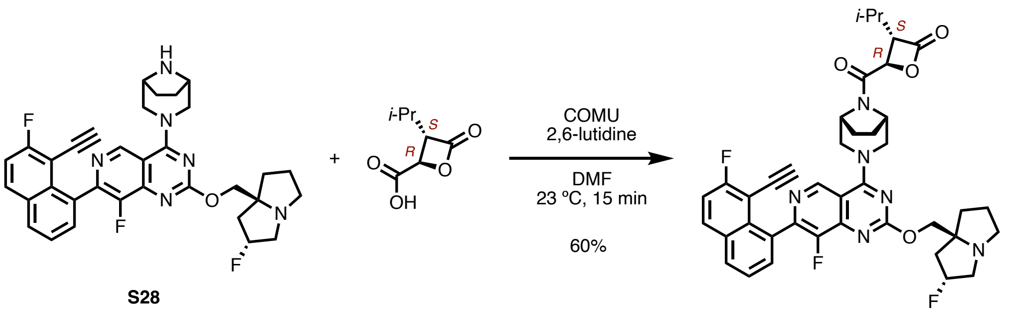


**S28** **(2*R*, 3*S*)-S23** **(2*R*, 3*S*)-5b**

**(2*R*,3*S*)-5b.** Synthesis of precursor **S28** was adapted from a known procedure [58]. Quality control was performed for the synthetic material by comparing the proton NMR to reported characterizations and the observed m/z to theoretical mass. $^1$H NMR (400 MHz, DMSO-d$_6$) δ 9.04 (s, 1H), 8.24 - 8.18 (m, 2H), 7.70 - 7.58 (m, 3H), 5.34 - 5.21 (m, 1H), 4.47 (d, *J* = 12.0 Hz, 1H), 4.33 (d, *J* = 12.0 Hz, 1H), 4.11 (dd, *J* = 10.4, 4.0 Hz, 1H), 4.09 - 3.98 (m, 2H), 3.63 - 3.58 (m, 1H), 3.54 (s, 3H), 3.09 - 3.07 (m, 2H), 3.01 (d, *J* = 11.2 Hz, 1H), 2.83 - 2.67 (m, 1H), 2.13 (d, *J* = 4.4 Hz, 1H), 2.05 (d, *J* = 3.2 Hz, 2H), 1.85 - 1.76 (m, 3H), 1.65 (s, 4H). LC-MS (ESI-Q) calculated for $C_{33}H_{31}F_3N_6O$ $[M + H]^+$ 584.25, found 585.25. A 4-mL dram-vial equipped with a stir bar was charged with **(2*R*,3*S*)-S23**, and COMU (8.8 mg, 0.021 mmol). DMF (200 µL) and 2,6-lutidine (5.5 mg, 0.051 mmol) were added. The mixture was stirred at room temperature for 10 min until it turned yellow. Amine **S28** (10.0 mg, 0.017 mmol) was added as a solid. The mixture was further stirred at room temperature for 15 min when the conversion of the starting material was apparent in LC-MS. The reaction mixture without work-up was directly loaded onto a tightly packed silica cartridge and purified by flash column chromatography (0–30% MeOH-DCM, 12-g Gold RediSep(R) Rf column, Teledyne ISCO, Lincoln, NE). The title compound **(2*R*,3*S*)-5b** was isolated as a yellow solid (7.5 mg,0.011 mmol, 60% yield). $^1$H NMR (600 MHz, DMSO) δ 9.16 – 9.02 (m, 1H), 8.31 – 8.15 (m, 2H), 7.78 – 7.56 (m, 3H), 5.56 – 5.19 (m, 3H), 4.81 – 4.60 (m, 3H), 4.50 (q, *J* = 17.5 Hz, 1H), 4.27 (s, 2H), 4.01 (d, *J* = 5.9 Hz, 1H), 3.95 – 3.83 (m, 1H), 3.71 (h, *J* = 12.8 Hz, 2H), 2.98 (s, 1H), 2.18 (tt, *J* = 14.2, 6.9 Hz, 4H), 2.03 – 1.74 (m, 7H), 1.23 (s, 1H), 1.14 – 0.96 (m, 5H). Accurate MS (ESI-QTOF) calculated for $C_{40}H_{40}F_3N_6O_4$ $[M + H]^+$ 725.3063, found 725.3051.

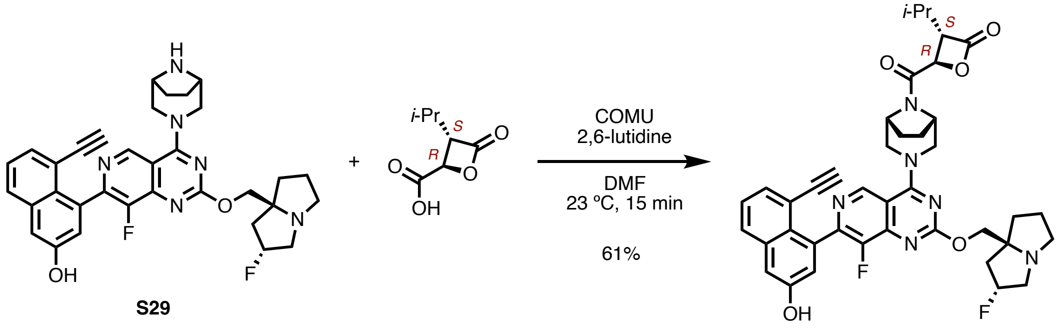


**S29** **(2*R*, 3*S*)-S23** **(2*R*, 3*S*)-5c**

**(2*R*,3*S*)-5c.** Synthesis of precursor **S29** was adapted from a known procedure [58]. Quality control was performed for the synthetic material by comparing the proton NMR to reported characterizations and the observed m/z to theoretical mass. $^1$H NMR (300 MHz, DMSO-$d_6$) δ 10.17 (s, 1H), 9.03 (s, 1H), 7.87 - 7.90 (m, 1H), 7.49 - 7.39 (m, 2H), 7.34 (d, *J* = 2.7 Hz, 1H), 7.12 (d, *J* = 2.4 Hz, 1H), 5.28 (d, *J* = 54.3 Hz, 1H), 4.48 (d, *J* = 12.3 Hz, 1H), 4.30 (d, *J* = 12.0 Hz, 1H), 4.08 - 4.12 (m, 1H), 3.98 - 4.02 (m, 1H), 3.68 - 3.49 (m, 5H), 3.11 (d, *J* = 7.2 Hz, 3H), 3.00 (s, 1H), 2.79 - 7.39 (m, 2H), 2.15 (d, *J* = 3.9 Hz, 1H), 2.08 - 1.95 (m, 2H), 1.89 - 1.74 (m, 3H), 1.66 (s, 3H). LC-MS (ESI-Q) calculated for $C_{33}H_{32}F_2N_6O_2$ [M + H]$^+$ 582.26, found 583.26. A 4-mL dram-vial equipped with a stir bar was charged with **(2*R*,3*S*)-S23**, and COMU (8.8 mg, 0.021 mmol). DMF (200 µL) and 2,6-lutidine (5.5 mg, 0.051 mmol) were added. The mixture was stirred at room temperature for 10 min until it turned yellow. Amine **S29** (10.0 mg, 0.017 mmol) was added as a solid. The mixture was further stirred at room temperature for 15 min when the conversion of the starting material was apparent in LC-MS. The reaction mixture without work-up was directly loaded onto a tightly packed silica cartridge and purified by flash column chromatography (0–30% MeOH-DCM, 12-g Gold RediSep(R) Rf column, Teledyne ISCO, Lincoln, NE). The title compound **(2*R*,3*S*)-5c** was isolated as a yellow solid (7.6 mg, 0.011 mmol, 61% yield). $^1$H NMR (600 MHz, DMSO) δ 10.70 (s, 1H), 10.17 (s, 1H), 9.09 (t, *J* = 6.4 Hz, 1H), 7.90 (d, *J* = 8.1 Hz, 1H), 7.50 – 7.40 (m, 2H), 7.36 (d, *J* = 2.5 Hz, 1H), 7.13 (d, *J* = 2.6 Hz, 1H), 5.44 (d, *J* = 4.2 Hz, 2H), 4.84 – 4.61 (m, 3H), 4.60 – 4.18 (m, 3H), 3.96 – 3.83 (m, 1H), 3.80 – 3.63 (m, 2H), 3.57 (d, *J* = 4.9 Hz, 1H), 2.45 – 1.73 (m, 11H), 1.24 (s, 1H), 1.12 – 0.97 (m, 5H). Accurate MS (ESI-QTOF) calculated for $C_{40}H_{41}F_2N_6O_5$ [M + H]$^+$ 723.3107, found 723.3104.

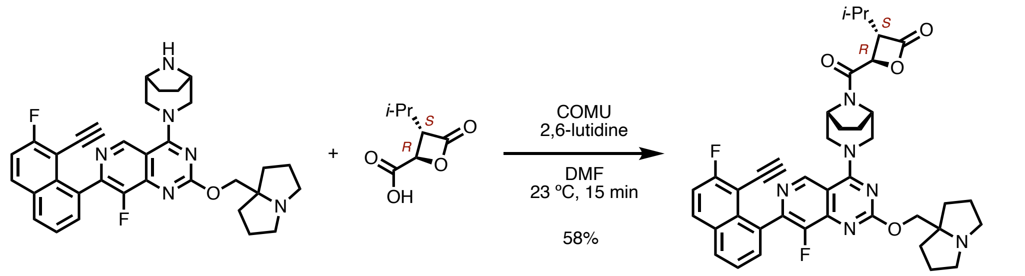


**S30**

**(2*R*, 3*S*)-S23**

**(2*R*, 3*S*)-5d**

**(2*R*,3*S*)-5d.** Synthesis of precursor **S30** was adapted from a known procedure [58]. Quality control was performed for the synthetic material by comparing the proton NMR to reported characterizations and the observed m/z to theoretical mass. $^1$H NMR (300 MHz, DMSO-$d_6$) δ 9.05 (s, 1H), 8.28 - 8.15 (m, 2H), 7.74 - 7.57 (m, 3H), 4.30 - 4.48 (m, 2H), 4.07 - 4.00 (m, 3H), 3.68 - 3.51 (m, 4H), 2.89 - 2.96 (m, 2H), 2.56 (d, *J* = 6.0 Hz, 1H), 2.55 (d, *J* = 2.4 Hz, 2H), 1.94 - 1.72 (m, 6H), 1.69 - 1.52 (m, 6H). LC-MS (ESI-Q) calculated for $C_{33}H_{32}F_2N_6O$ [M + H]$^+$ 566.26, found 567.26. A 4-mL dram-vial equipped with a stir bar was charged with **(2*R*,3*S*)-S23**, and COMU (8.8 mg, 0.021 mmol). DMF (200 µL) and 2,6-lutidine (5.5 mg, 0.051 mmol) were added. The mixture was stirred at room temperature for 10 min until it turned yellow. Amine **S30** (10.0 mg, 0.017 mmol) was added as a solid. The mixture was further stirred at room temperature for 15 min when the conversion of the starting material was apparent in LC-MS. The reaction mixture without work-up was directly loaded onto a tightly packed silica cartridge and purified by flash column chromatography (0–30% MeOH-DCM, 12-g Gold RediSep(R) Rf column, Teledyne ISCO, Lincoln, NE). The title compound **(2*R*,3*S*)-5d** was isolated as a yellow solid (7.1 mg, 0.011 mmol, 58% yield). $^{1}$H NMR (600 MHz, DMSO) δ 10.16 (s, 1H), 9.13 (t, *J* = 6.9 Hz, 1H), 8.32 – 8.17 (m, 2H), 7.78 – 7.56 (m, 3H), 5.44 (t, *J* = 4.1 Hz, 1H), 4.77 (t, *J* = 7.6 Hz, 1H), 4.73 – 4.62 (m, 2H), 4.55 (q, *J* = 12.7 Hz, 3H), 3.98 (d, *J* = 6.2 Hz, 1H), 3.95 – 3.84 (m, 1H), 3.81 – 3.65 (m, 2H), 3.49 (s, 2H), 3.22 (s, 2H), 2.26 – 1.70 (m, 12H), 1.23 (s, 1H), 1.16 – 0.95 (m, 5H). Accurate MS (ESI-QTOF) calculated for $C_{40}H_{41}F_2N_6O_4$ [M + H]$^+$ 707.3157, found 707.3165.

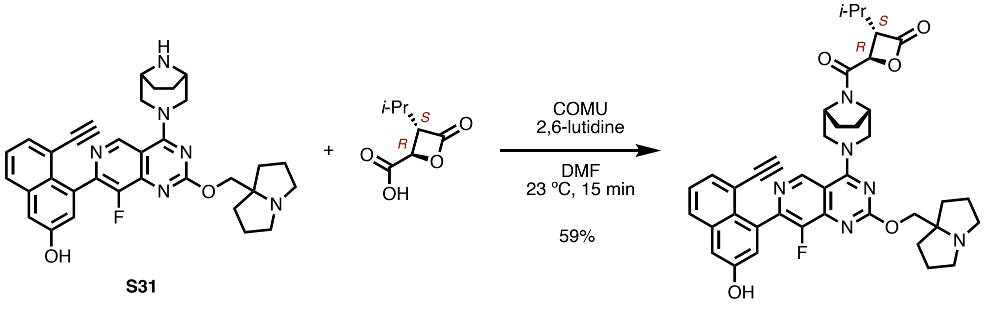


**S31** **(2*R*, 3*S*)-S23** **(2*R*, 3*S*)-5e**

**(2*R*,3*S*)-5e.** Synthesis of precursor **S31** was adapted from a known procedure [58]. Quality control was performed for the synthetic material by comparing the proton NMR to reported characterizations and the observed m/z to theoretical mass. $^{1}$H NMR (400 MHz, DMSO-d$_6$) δ 10.17 (s, 1H), 9.02 (s, 1H), 7.88 (dd, *J*=8.0, 1.7 Hz, 1H), 7.46 (dd, *J*=7.1, 1.7 Hz, 1H), 7.44 - 7.38 (m, 1H), 7.34 (d, *J*=2.6 Hz, 1H), 7.12 (d, *J*=2.6 Hz, 1H), 4.46 (d, *J*=12.0 Hz, 1H), 4.30 (d, *J*=12.0 Hz, 1H), 4.02 (s, 2H), 3.63 (d, *J*=12.1 Hz, 1H), 3.57 (d, *J*=17.1 Hz, 4H), 2.95 - 2.89 (m, 2H), 2.56 - 2.49 (m,2H), 1.90 - 1.85 (m, 2H), 1.81 - 1.72 (m, 4H), 1.65 - 1.55 (m, 6H). LC-MS (ESI-Q) calculated for $C_{33}H_{33}FN_6O_2$ [M + H]$^+$ 565.27, found 565.20. A 4-mL dram-vial equipped with a stir bar was charged with **(2*R*,3*S*)-S23**, and COMU (8.8 mg, 0.021 mmol). DMF (200 µL) and 2,6-lutidine (5.5 mg, 0.051 mmol) were added. The mixture was stirred at room temperature for 10 min until it turned yellow. Amine **S31** (10.0 mg, 0.017 mmol) was added as a solid. The mixture was further stirred at room temperature for 15 min when the conversion of the starting material was apparent in LC-MS. The reaction mixture without work-up was directly loaded onto a tightly packed silica cartridge and purified by flash column chromatography (0–30% MeOH-DCM, 12-g Gold RediSep(R) Rf column, Teledyne ISCO, Lincoln, NE). The title compound **(2*R*,3*S*)-5e** was isolated as a yellow solid (7.1 mg, 0.010 mmol, 59% yield). $^{1}$H NMR (600 MHz, DMSO) δ 10.17 (d, *J* = 1.4 Hz, 2H), 9.11 (t, *J* = 6.7 Hz, 1H), 7.90 (dd, *J* = 8.2, 1.6 Hz, 1H), 7.50 – 7.40 (m, 2H), 7.36 (t, *J* = 2.0 Hz, 1H), 7.13 (d, *J* = 2.8 Hz, 1H), 5.44 (d, *J* = 4.1 Hz, 1H), 4.77 (s, 1H), 4.73 – 4.61 (m, 2H), 4.54 (t, *J* = 9.7 Hz, 3H), 3.96 – 3.83 (m, 1H), 3.81 – 3.63 (m, 2H), 3.57 – 3.44 (m, 3H), 3.21 (s, 2H), 2.26 – 1.71 (m, 14H), 1.23 (s, 1H), 1.14 – 0.97 (m, 6H). Accurate MS (ESI-QTOF) calculated for $C_{40}H_{42}FN_6O_5$ [M + H]$^+$ 705.3201, found 705.3196.

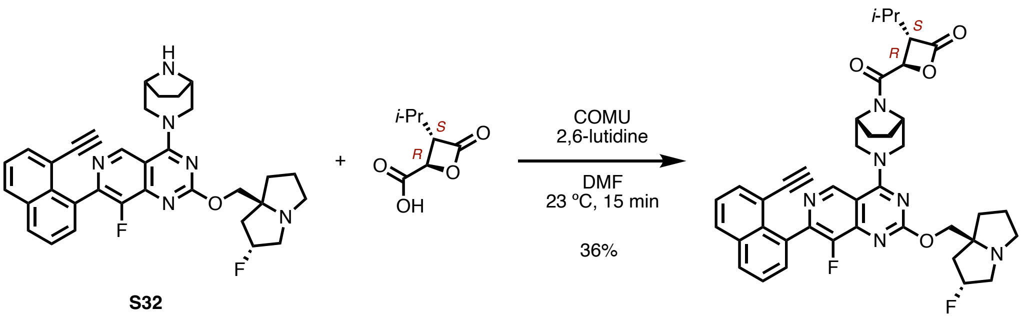


**S32** **(2*R*, 3*S*)-S23** **(2*R*, 3*S*)-5f**

**(2*R*,3*S*)-5f.** Synthesis of precursor **S32** was adapted from a known procedure [58]. Quality control was performed for the synthetic material by comparing the proton NMR to reported characterizations and the observed m/z to theoretical mass. $^1$H NMR (300 MHz, DMSO-$d_6$) δ 9.05 (s, 1H), 8.16 - 8.12 (m, 2H), 7.73 - 7.67 (m, 2H), 7.59 - 7.54 (m, 2H), 5.37 - 5.19 (m, 1H), 4.48 (d, *J* = 12.2 Hz, 1H), 4.32 (d, *J* = 12.2 Hz, 1H), 4.11 (dd, *J* = 10.3, 2.4 Hz, 1H), 4.01 (dd, *J* = 10.3, 2.5 Hz, 1H), 3.68 - 3.56 (m, 5H), 3.10 (d, *J* = 6.6 Hz, 2H), 3.00 (s, 1H), 2.86 - 2.82 (m, 1H), 2.14 (s, 1H), 2.04 - 1.99 (m, 2H), 1.85 - 1.76 (m, 3H), 1.66 (s, 4H). LC-MS (ESI-Q) calculated for $C_{33}H_{32}F_2N_6O$ [M + H]$^+$ 567.27, found 565.20. A 4-mL dram-vial equipped with a stir bar was charged with **(2*R*,3*S*)-S23**, and COMU (8.8 mg, 0.021 mmol). DMF (200 µL) and 2,6-lutidine (5.5 mg, 0.051 mmol) were added. The mixture was stirred at room temperature for 10 min until it turned yellow. Amine **S32** (10.0 mg, 0.017 mmol) was added as a solid. The mixture was further stirred at room temperature for 15 min when the conversion of the starting material was apparent in LC-MS. The reaction mixture without work-up was directly loaded onto a tightly packed silica cartridge and purified by flash column chromatography (0–30% MeOH-DCM, 12-g Gold RediSep(R) Rf column, Teledyne ISCO, Lincoln, NE). The title compound **(2*R*,3*S*)-5f** was isolated as a yellow solid (4.4 mg, 0.004 mmol, 36% yield). $^1$H NMR (600 MHz, DMSO) δ 10.69 (s, 2H), 9.09 (t, *J* = 6.3 Hz, 1H), 8.15 (dd, *J* = 14.3, 8.2 Hz, 2H), 7.76 – 7.67 (m, 2H), 7.58 (q, *J* = 7.3 Hz, 2H), 5.44 (d, *J* = 4.1 Hz, 3H), 4.77 (d, *J* = 7.3 Hz, 1H), 4.66 (q, *J* = 8.1 Hz, 2H), 4.50 (d, *J* = 12.8 Hz, 2H), 4.33 (s, 3H), 3.97 – 3.83 (m, 2H), 3.83 – 3.59 (m, 4H), 2.40 – 1.70 (m, 12H), 1.23 (s, 1H), 1.12 – 0.96 (m, 6H). Accurate MS (ESI-QTOF) calculated for $C_{40}H_{41}F_2N_6O_4$ [M + H]$^+$ 707.3157, found 707.3165.

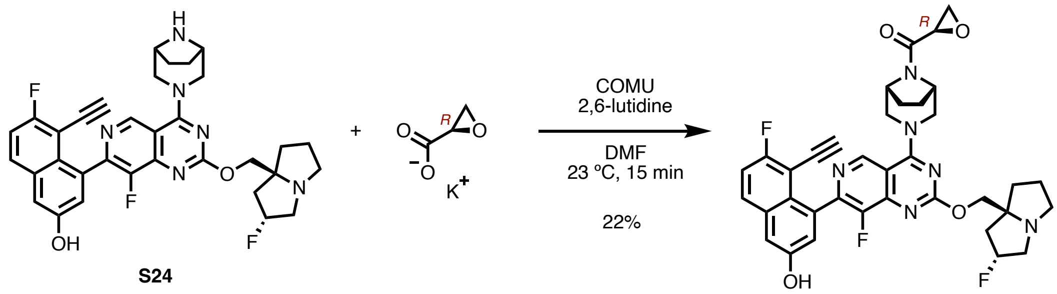


**S24** **(*R*)-S33**

**(*R*)-S33.** A 4-mL dram vial equipped with a stir bar was charged with potassium (2R)-oxirane-2-carboxylate (5.0 mg, 0.040 mmol, 1.2 equiv), COMU (17.1 mg, 0.040 mmol, 1.2 equiv). DMF (333 μL, 0.1 M), and 2,6-lutidine (12 μL, 3 equiv) were added via pipette. The mixture was stirred at room temperature for 10 min before **S24** (20.0 mg, 0.033 mmol) was added as a solid. The reaction mixture was stirred at room temperature for 10 min. The mixture was directly loaded onto a packed silica cartridge and was purified by flash column chromatography (0–30% MeOH-DCM, 12-g Gold RediSep(R) Rf column, Teledyne ISCO, Lincoln, NE) to give the title compound as a yellow solid (8.5 mg, 0.013 mmol, 22% yield). $^1$H NMR (600 MHz, DMSO) δ 10.64 (s, 2H), 10.15 (s, 1H), 9.08 (dd, *J* = 6.6, 3.9 Hz, 1H), 7.95 (dd, *J* = 9.2, 5.9 Hz, 1H), 7.43 (t, *J* = 9.0 Hz, 1H), 7.37 (d, *J* = 2.6 Hz, 1H), 7.14 (d, *J* = 3.2 Hz, 1H), 5.67 – 5.36 (m, 2H), 4.94 – 4.36 (m, 8H), 4.06 – 3.50 (m, 10H), 3.15 – 2.82 (m, 7H), 2.32 – 1.65 (m, 11H), 1.19 (s, 2H). Accurate MS (ESI-QTOF) calculated for $C_{36}H_{34}F_3N_6O_4$ $[M + H]^+$ 671.2594, found 671.2692.

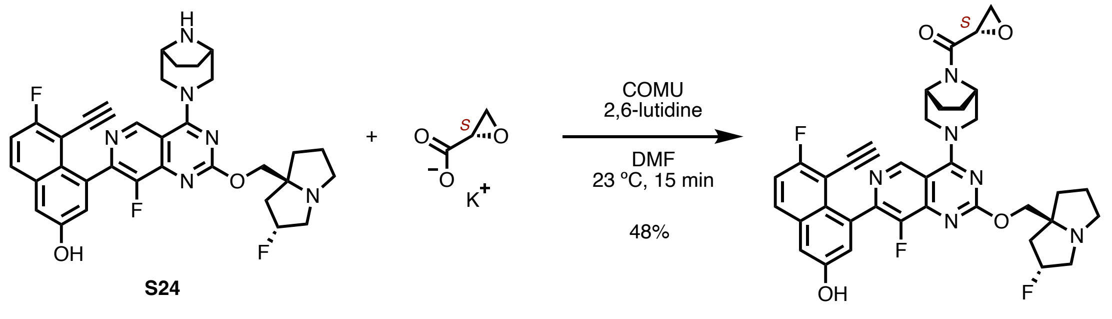


**S24** (***S***)**-S33**

(***S***)**-S33.** A 4-mL dram vial equipped with a stir bar was charged with potassium (2S)-oxirane-2-carboxylate (5.0 mg, 0.040 mmol, 1.2 equiv), COMU (17.1 mg, 0.040 mmol, 1.2 equiv). DMF (333 μL, 0.1 M), and 2,6-lutidine (12 μL, 3 equiv) were added via pipette. The mixture was stirred at room temperature for 10 min before **S24** (20.0 mg, 0.033 mmol) was added as a solid. The reaction mixture was stirred at room temperature for 10 min. The mixture was directly loaded onto a packed silica cartridge and was purified by flash column chromatography (0–30% MeOH-DCM, 12-g Gold RediSep(R) Rf column, Teledyne ISCO, Lincoln, NE) to give the title compound as a yellow solid (10.8 mg, 0.016 mmol, 48% yield). $^1$H NMR (400 MHz, DMSO) δ 10.64 (s, 1H), 10.16 (s, 1H), 9.08 (s, 1H), 7.95 (dd, *J* = 9.2, 6.0 Hz, 1H), 7.51 – 7.30 (m, 2H), 7.14 (s, 1H), 5.69 – 5.34 (m, 1H), 4.96 – 4.31 (m, 7H), 4.11 – 3.46 (m, 11H), 2.41 – 1.62 (m, 14H), 1.19 (s, 1H).Accurate MS (ESI-QTOF) calculated for $C_{36}H_{34}F_3N_6O_4$ $[M + H]^+$ 671.2594, found 671.2692.

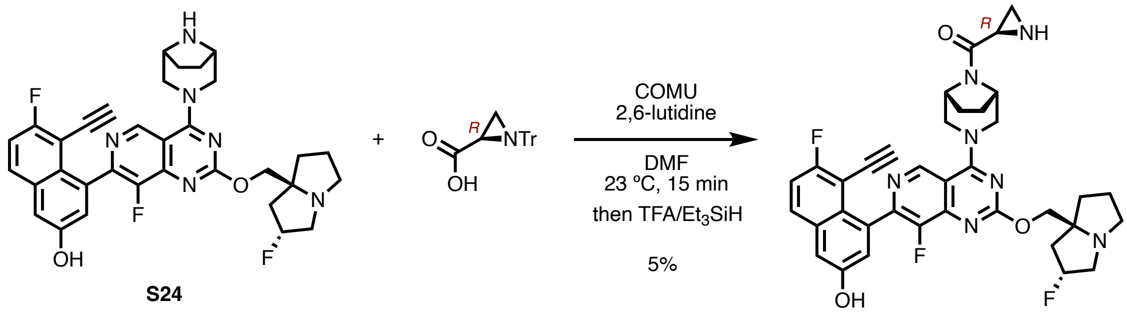


**S24** (***R***)**-S34**

**(*R*)-S34.** A dram-vial equipped with a stir bar was charged with (2R)-1-tritylaziridine-2-carboxylic acid (19.8 mg, 0.060 mmol), COMU (32.1 mg, 0.075 mmol). DMF (0.500 mL) and 2,6-lutidine (0.02 mL, 0.150 mmol) were added. The mixture was stirred at room temperature for 15 min before **S24** (30.0 mg, 0.050 mmol) was added in one batch. The resulting mixture was stirred at room temperature for another 30 min before a flash column chromatography to remove COMU-oxime byproduct. The crude was dissolved in DCM (0.5 mL) and cooled to 0 ºC. TFA (31 µL, 8 equiv), triethylsilane (32 µL, 4 equiv), and DIPEA (87 µL, 10 equiv) were added. The mixture was stirred 0 ºC for 30 min, and then room temperature for 1 h before concentrated. The crude was purified by reverse-phase HPLC (5–95% MeCN-H2O) to give the title compound as a pale yellow solid (1.6 mg, 0.002 mmol, 5% yield). The compound was not stable by its own. Bronsted acid was found to accelerate the hydrolysis. Accurate MS (ESI-QTOF) calculated for $C_{36}H_{35}F_3N_7O_3$ [M + H]$^+$ 670.2853, found 670.2831. **Note:** The compound was not stable by its own. Bronsted acid was found to accelerate the hydrolysis. Obtaining NMRs for pure material was not possible.

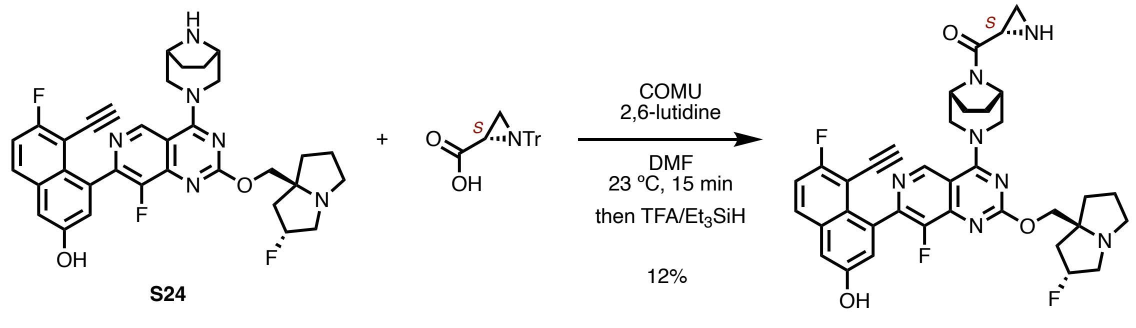


**(*S*)-S34.** A dram-vial equipped with a stir bar was charged with (2S)-1-tritylaziridine-2-carboxylic acid (19.8 mg, 0.060 mmol), COMU (32.1 mg, 0.075 mmol). DMF (0.500 mL) and 2,6-lutidine (0.02 mL, 0.150 mmol) were added. The mixture was stirred at room temperature for 15 min before **S24** (30.0 mg, 0.050 mmol) was added in one batch. The resulting mixture was stirred at room temperature for another 30 min before a flash column chromatography to remove COMU-oxime byproduct. The crude was dissolved in DCM (0.5 mL) and cooled to 0 ºC. TFA (31 µL, 8 equiv), triethylsilane (32 µL, 4 equiv), and DIPEA (87 µL, 10 equiv) were added. The mixture was stirred 0 ºC for 30 min, and then room temperature for 1 h before concentrated. The crude was purified by reverse-phase HPLC (5–95% MeCN-H2O) to give the title compound as a pale yellow solid (4.0 mg, 0.006 mmol, 12% yield). Accurate MS (ESI-QTOF) calculated for $C_{36}H_{35}F_3N_7O_3$ [M + H]$^+$ 670.2853, found 670.2831. **Note:** The compound was not stable by its own. Bronsted acid was found to accelerate the hydrolysis. Obtaining NMRs for pure material was not possible.

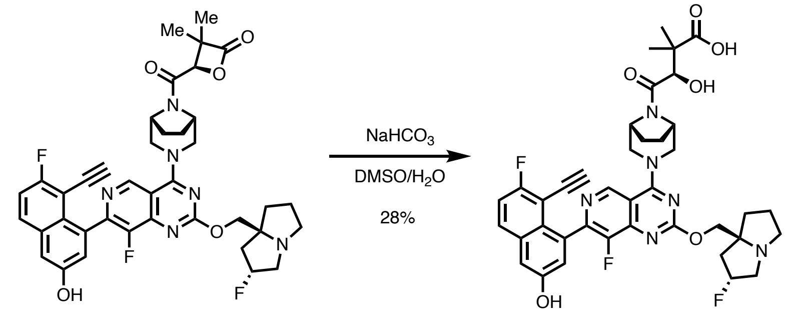


(***R***)**-7** **S35**

**S35.** A 4-mL dram vial equipped with a stir bar was charged with (***R***)**-7** (20 mg, 0.028 mmol) in DMSO (275 µL). Saturated sodium bicarbonate solution (275 µL) was added. The mixture was stirred at room temperature for 3 days before purified by reverse phase HPLC (5–95% MeCN-H2O without TFA) to give the title compound as a pale yellow solid (5.7 mg, 0.008 mmol, 28% yield). **Note:** Compound **S35** is sensitive to Bronsted acid (formic acid, TFA, etc.), which protonates the carboxylic acid and catalyzes the amide bond cleavage intramolecularly. Purification and concentration should be run with care in an acid-free environment. $^{1}$H NMR (600 MHz, DMSO) δ 9.06 (dd, *J* = 7.8, 3.2 Hz, 1H), 7.96 (dd, *J* = 9.2, 5.9 Hz, 1H), 7.45 (t, *J* = 9.0 Hz, 1H), 7.40 (d, *J* = 2.6 Hz, 1H), 7.19 (d, *J* = 2.7 Hz, 1H), 5.36 – 5.20 (m, 1H), 4.94 – 4.34 (m, 6H), 4.17 – 3.99 (m, 2H), 3.94 (d, *J* = 4.3 Hz, 1H), 3.85 – 3.49 (m, 4H), 3.16 – 2.99 (m, 3H), 2.83 (td, *J* = 8.8, 5.8 Hz, 1H), 2.19 – 1.70 (m, 10H), 1.27 – 1.09 (m, 6H). $^{13}$C NMR (151 MHz, DMSO) δ 178.15, 178.02, 177.99, 167.84, 167.82, 167.54, 165.24, 165.23, 165.19, 165.14, 165.09, 163.63, 162.82, 161.18, 154.25, 154.23, 151.41, 151.38, 149.72, 149.70, 149.67, 147.95, 147.92, 147.87, 147.84, 145.04, 145.01, 144.96, 144.94, 144.92, 144.01, 143.97, 133.94, 133.90, 132.51, 130.55, 130.49, 125.11, 125.07, 122.97, 118.09, 116.03, 115.86, 111.60, 111.22, 111.17, 103.90, 103.80, 98.49, 98.44, 97.34, 97.28, 91.56, 91.52, 91.49, 91.45, 74.85, 74.39, 74.32, 73.21, 73.14, 71.90, 59.86, 59.82, 59.73, 59.70, 56.37, 55.31, 55.02, 54.63, 54.57, 54.27, 54.20, 54.10, 53.86, 51.37, 51.33, 51.30, 51.24, 45.01, 44.98, 44.92, 42.49, 42.45, 42.36, 42.32, 35.63, 35.62, 27.40, 27.32, 27.16, 27.08, 25.31, 25.22, 25.18, 25.12, 22.98, 22.94, 22.86, 22.80, 21.00, 20.63, 1.17. $^{19}$F NMR (564 MHz, DMSO) δ -73.47 (TFA), -110.77 – -110.80 (m, 1F), -139.89 – -140.10 (m, 1F), -171.99 – -172.20 (m, 1F). Accurate MS (ESI-QTOF) calculated for $C_{39}H_{40}F_3N_6O_6$ $[M + H]^+$ 745.2961, found 745.3125.

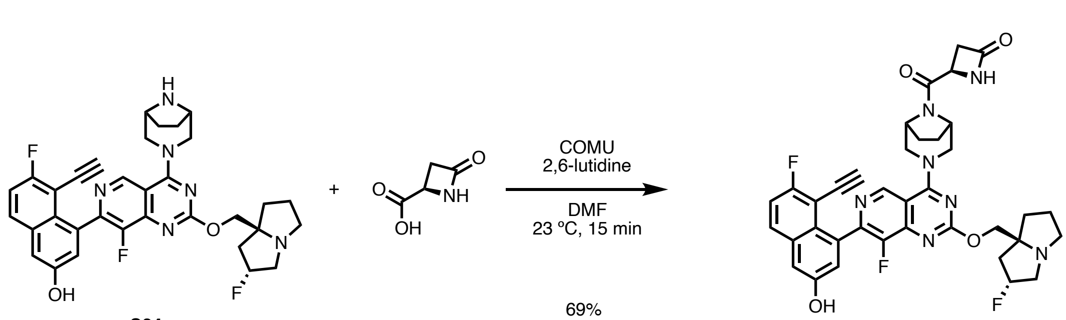


**S24** (*R*)-**S36**

**(*R*)-S36.** A 4-mL dram vial equipped with a stir bar was charged with (2R)-4-oxoazetidine-2-carboxylic acid (3.5 mg, 0.030 mmol, 1.5 equiv), COMU (12.8 mg, 0.030 mmol, 1.5 equiv). DMF (200 µL, 0.1 M) and 2,6-lutidine (4.6 µL, 2 equiv) were added via pipette. The mixture was stirred at room temperature for 10 min before **S24** (12.0 mg, 0.020 mmol) was added as a solid. The mixture was stirred for another 15 min at which point full conversion of the **S24** was apparent. The reaction mixture was purified by reverse-phase HPLC (5–95% MeCN-H2O with 0.1% TFA) to give the title compound as a pale yellow solid (9.7 mg, 0.014 mmol, 69% yield). $^{1}H$ NMR (400 MHz, DMSO) δ 10.88 (s, 1H), 10.19 (s, 1H), 9.17 – 9.06 (m, 1H), 8.40 – 8.28 (m, 1H), 7.99 (dd, *J* = 9.2, 5.9 Hz, 1H), 7.47 (t, *J* = 9.0 Hz, 1H), 7.41 (d, *J* = 2.6 Hz, 1H), 7.19 (d, *J* = 2.7 Hz, 1H), 5.71 – 5.42 (m, 2H), 4.81 – 4.36 (m, 8H), 3.94 – 3.58 (m, 6H), 3.38 – 3.18 (m, 2H), 3.07 (d, *J* = 14.4 Hz, 1H), 2.87 (d, *J* = 14.4 Hz, 1H), 2.68 – 2.53 (m, 1H), 2.42 – 2.27 (m, 1H), 2.18 (tq, *J* = 13.6, 6.2 Hz, 2H), 2.12 – 1.67 (m, 5H). $^{19}F$ NMR (376 MHz, DMSO) δ -110.60 (d, *J* = 7.5 Hz, 1F), -139.52 – -139.78 (m, 1F), -172.85 – -173.05 (m, 1F). Accurate MS (ESI-QTOF) calculated for $C_{37}H_{35}F_3N_7O_4$ $[M + H]^+$ 698.2702, found 698.2859.

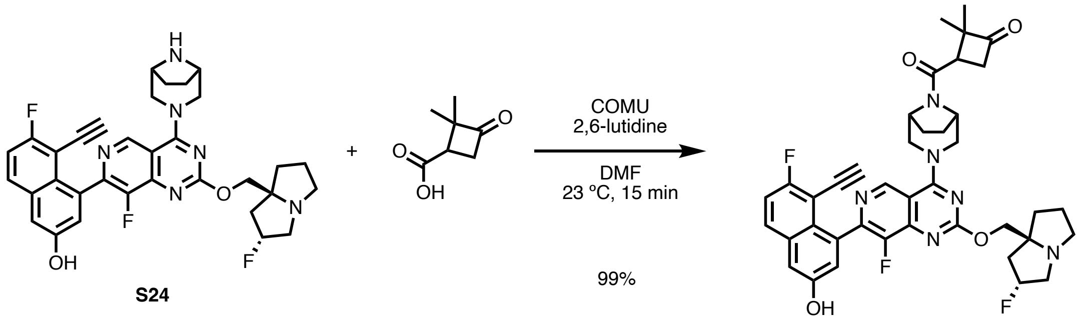


**S37.** A 4-mL dram-vial equipped with a stir bar was charged with 2,2-dimethyl-3-oxo-cyclobutanecarboxylic acid (8.5 mg, 0.060 mmol), COMU (25.7 mg, 0.060 mmol). DMF (0.250 mL) and 2,6-lutidine (0.02 mL, 0.150 mmol) were added. The mixture was stirred at room temperature for 10 min before **S24** (30.0 mg, 0.050 mmol) was added. The reaction mixture was further stirred for 30 min, at which point full consumption of the starting material was apparent. The reaction mixture was directly loaded onto a packed silica cartridge and purified by reverse-phase HPLC (5–95% MeCN-H2O) to give the title compound as a yellow solid (43.2 mg, 0.050 mmol, 99% yield). Accurate MS (ESI-QTOF) calculated for $C_{40}H_{40}F_3N_6O_4$ $[M + H]^+$ 725.3063, found 725.3246. $^{1}H$ NMR (400 MHz, DMSO) δ 10.69 (s, 1H), 10.20 (s, 1H), 9.22 – 9.09 (m, 1H), 8.00 (dd, *J* = 9.2, 5.9 Hz, 1H), 7.48 (t, *J* = 9.0 Hz, 1H), 7.42 (d, *J* = 2.5 Hz, 1H), 7.22 – 7.17 (m, 1H), 5.65 – 5.41 (m, 1H), 4.93 – 4.42 (m, 6H), 3.95 – 3.87 (m, 1H), 3.87 – 3.04 (m, 13H), 2.44 (s, 1H), 2.29 (d, *J* = 12.6 Hz, 1H), 2.22 – 1.71 (m, 7H), 1.41 – 1.31 (m, 3H), 1.25 – 0.91 (m, 3H). $^{13}C$ NMR (151 MHz, DMSO, two diastereomers) δ 211.76, 211.72, 173.35, 166.55, 166.53, 166.32, 166.28, 165.14, 165.13, 165.11, 165.10, 165.08, 162.89, 162.84, 162.82, 162.81, 161.26, 154.16, 154.14, 151.32, 149.68, 149.65, 149.62, 147.72, 147.69, 147.67, 147.64, 147.61, 147.59, 145.34, 145.31, 145.25, 145.22, 144.27, 144.23, 133.80, 133.76, 132.50, 130.66, 130.60, 125.14, 125.12, 125.07, 122.94, 116.12, 115.94, 111.68, 111.32, 111.25, 111.18, 103.83, 103.81, 103.72, 103.70, 91.41, 91.38, 91.37, 91.34, 74.95, 74.93, 62.78, 62.66, 59.48, 59.42, 59.34, 59.28, 57.40, 57.38, 54.89, 54.27, 54.23, 54.12, 54.07, 53.63, 51.50, 51.43, 51.30, 51.24, 48.61, 44.95, 44.59, 44.58,

44.56, 38.06, 38.02, 37.24, 37.22, 34.95, 27.46, 27.42, 27.36, 27.30, 25.92, 25.80, 25.07, 24.95, 24.55, 24.52, 22.71, 22.57, 22.55, 22.43, 22.35, 18.30, 18.15, 17.81, 17.80. $^{19}$F NMR (376 MHz, DMSO) δ -70.13 (d, *J* = 710 Hz, 6F, PF6), -110.54 – -110.64 (m, 1F), -139.62 – -139.90 (m, 1F), -172.86 (s, 1F). Accurate MS (ESI-QTOF) calculated for $C_{40}H_{40}F_3N_6O_4$ $[M + H]^+$ 725.3063, found 725.3246.

**Supplementary Reference**


58 Wang, X. *et al.* KRAS G12D Inhibitors. WO2021041671A1 (2021).

59 Wang, X. L. *et al.* Identification of MRTX1133, a Noncovalent, Potent, and Selective KRAS(G12D) Inhibitor. *Journal of Medicinal Chemistry* (2021). https://doi.org:10.1021/acs.jmedchem.1c01688

60 LeboucherDurand, M. A., Langlois, V. & Guerin, P. 4-carboxy-2-oxetanone as a new chiral precursor in the preparation of functionalized racemic or optically active poly(malic acid) derivatives. *Polymer Bulletin* **36**, 35-41 (1996). https://doi.org:Doi 10.1007/Bf00296005

61 Kawamura, S., Unno, Y., Asai, A., Arisawa, M. & Shuto, S. Design and synthesis of the stabilized analogs of belactosin A with the unnatural cis-cyclopropane structure. *Organic & Biomolecular Chemistry* **11**, 6615-6622 (2013). https://doi.org:10.1039/C3OB41338A

62 Tello-Aburto, R., Hallada, L. P., Niroula, D. & Rogelj, S. Total synthesis and absolute stereochemistry of the proteasome inhibitors cystargolides A and B. *Organic & Biomolecular Chemistry* **13**, 10127-10130 (2015). https://doi.org:10.1039/C5OB01821H

63 Belibel, R. & Barbaud, C. Synthesis of new optically active alpha, alpha ',beta-trisubstituted-beta-lactones as monomers for stereoregular biopolyesters. *Journal of Polymer Science, Part a: Polymer Chemistry* **53**, 2586-2597 (2015). https://doi.org:10.1002/pola.27724
